# Orbitrap Collision Cross Section Measurements Enhance Isomer Annotations in Lipidomics

**DOI:** 10.64898/2026.07.03.735735

**Authors:** Ziqin Ni, Konstantin Ayzikov, Alexander Makarov, Samuel Moore, David A. Gaul, Kyle L. Fort, Facundo M. Fernández

## Abstract

Despite advances in high-resolution mass spectrometry (HRMS), confident lipid annotation remains challenging due to the extensive chemical diversity of the lipidome and the prevalence of isomeric species. Ion mobility collision cross section (CCS) measurements provide structural information that complements HRMS; however, not all HRMS platforms can perform these measurements, necessitating a trade-off among mass resolution, accuracy, and robustness. Here, we introduce a method to infer lipid CCS values directly from liquid chromatography (LC)-Orbitrap MS experiments (*^Orbi^CCS*). We show that Orbitrap mass analyzer pressure readings, and therefore CCS values, are influenced by the LC gradient solvent composition, requiring correction using isotopically labeled internal standards injected post-column. We also show that hundreds of lipid features can be assigned *^Orbi^CCS* values in a single LC run, with average precision better than 1% and an accuracy of 1–2% relative to reference *^DT^CCS* and *^TIMS^CCS* values. This excellent CCS accuracy not only enables more reliable annotation of lipid species in complex mixtures by matching *^Orbi^CCS* values to reference databases but also accelerates lipid structural elucidation based on the unknown’s position in *^Orbi^CCS*-retention time-*m/z* space.

## Introduction

Lipids play a crucial role in cell membrane structure, signaling, and disease states such as cancer. However, lipidomes from biological systems can be highly complex, requiring molecular analysis tools with high sensitivity and selectivity. Orbitrap™ mass spectrometry (MS) is known for its reliability, ultrahigh mass resolution, excellent mass accuracy, and stability, making it an essential tool for analyzing complex lipid mixtures. While accurate Orbitrap *m/z* measurements can suggest potential lipid classes and molecular formulas, additional structural data is needed to enhance annotation confidence, differentiate isomers and isobars, and obtain more detailed structural insights. In addition to tandem mass spectrometry (MS/MS), ion mobility collision cross section (CCS or σ) values provide complementary molecular information by correlating with the rotationally averaged size and charge of gas-phase ions. Therefore, interest in CCS-assisted lipid structural elucidation has been increasing^1–3^. Typically, CCS values are obtained from specialized ion mobility (IM) mass spectrometry systems based on drift tube IMS (DTIMS)^4^, trapped ion mobility spectrometry (TIMS)^5^, traveling wave ion mobility spectrometry (TWIMS)^6^, and structures for lossless ion manipulations (SLIM)^7^. In these systems, ions are separated based on their mobilities in an inert gas, typically N_2_, under dynamic or static electric fields^8^. A lesser-known fact is that Orbitrap mass analyzers can also indirectly measure CCS values by collision-induced ion losses within the analyzer, offering an effective way to improve lipid annotation workflows in complex mixtures without additional IM hardware.

Following their electrostatic injection and trapping, ions in the Orbitrap mass analyzer are characterized by their high kinetic energies (1.5−3.6 keV/charge)^9^. In this high-energy regime, ion-neutral collisions are best described by the hard-sphere collision model^10,11^, where a single ion-neutral collision is usually enough to eject an ion from its ion packet, either through scattering or activation followed by decay. For smaller ions, such as the lipid species with *m/z* < 2000 considered here, deriving CCS values via the energetic hard-sphere collision model, assuming that the rate of ion loss is proportional to the number of ion-neutral collisions within the Orbitrap analyzer, is a reasonable approximation. However, recent studies have shown that ions with a mass exceeding one megadalton can survive collisions in this energy regime, exhibiting a gradual frequency shift^12^. For large ions, collisional energy can be more readily distributed into vibrational modes rather than causing dissociation^13^. Therefore, when determining CCS values for lower-charge states of large proteins (>17.5–232 kDa), the energetic hard-sphere collision model is insufficient, and the collisional energy must be adjusted to account for the fraction absorbed into vibrations^14^.

The number of ion-neutral collisions in the Orbitrap mass analyzer can be estimated based on the ions’ flight path length (*l*) and their mean free path (*λ*). Ion packets inside the analyzer oscillate at MHz frequencies along the center electrode after undergoing electrodynamic squeezing and trapping. The cumulative ion flight path (*l)* per second can be estimated as *l*= *Lf_r_*, where *L* is the average ion path length (typically 65.1 mm), and *f_r_* is axial oscillation frequency in kHz. The mean free path (*λ*) is defined as the average distance between consecutive ion-neutral collisions, which can be estimated as 1/nσ, where *n* is the number of background neutral gas molecules and σ is the ions’ collision cross section. The number of ion-neutral collisions per second can thus be determined as 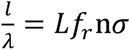. Following the energetic ion-neutral collision model, we expect the number of ion-neutral collisions to be equivalent to the number of ion loss events, which reflects a decay rate of −*d*(*Lf_r_*nσ)/*dt* from the starting number of ions N_0_. By substituting *n= P/kT* from the ideal gas law, the decay constant (*c*) can be rewritten as 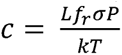. After rearranging this equation, we obtain:

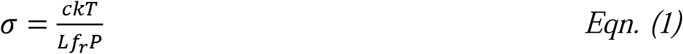

The conventional method for determining *c* involves fitting an exponential function to the transient signal obtained experimentally^15^. Using this approach, previous studies determined CCS values for proteins in the 8.5–17.5 kDa range, with an average difference of 7% compared to CCS values measured by IM techniques^16,17^. However, acquiring the transient signals requires either specialized software or the addition of an extra data-acquisition module to the existing Orbitrap instrument. An alternative method to estimate *c* is to use the full-width at half-maximum (FWHM) of the peak in the frequency domain^16^. This method has been successfully applied in Fourier Transform Ion Cyclotron Resonance MS^18^.

Here, we demonstrate an Orbitrap CCS measurement strategy that enables determination of lipid *^Orbi^CCS* values with high accuracy (1-2%) and precision (RSD < 1%). In this approach, the decay constant *c* is calculated as the ratio of the ion’s axial frequency and the monoisotopic peak resolving power (*R_p_*), *c* = *f_r_/R_p_*. Because this formula is exact only for transients significantly longer than the decay constant, for real-life transients with limited decay, internal calibrants must be used to deduce CCS. To improve the accuracy of *R_p_* estimation, the resolving power of all isotopic peaks with signal-to-noise ratios (*S/N*) above a predefined threshold was determined and averaged to enhance statistical robustness. Because lipids have lower CCS values than proteins, higher pressure was required in the Orbitrap mass analyzer to induce a measurable transient signal decay. To achieve such conditions, the rotational speed of the turbomolecular pump in the Orbitrap chamber was reduced to values that enabled the use of Eqn. (1) to derive *^Orbi^CCS* accurately. Any systematic bias in *^Orbi^CCS* measurements induced by pressure changes was corrected using appropriate CCS internal standards^17^.

## Materials and Methods

### Experimental Configurations

Analyses were performed on a research-grade Thermo Scientific™ Orbitrap Exploris™ 240 mass spectrometer coupled to a Vanquish Horizon UPLC system. The Orbitrap mass spectrometer was modified with a frequency controller to adjust the turbomolecular pump’s rotational speed. Because the pump speed governs the ultrahigh-vacuum (UHV) conditions within the Orbitrap chamber, reducing its speed directly increased analyzer pressure in a controlled fashion. A T-junction line before the electrospray ionization (ESI) source was setup to mix LC eluate with a continuous stream of the internal standard solution.

### Chemicals and sample preparation

This study analyzed SRM 1950 human plasma serum from the National Institute of Standards and Technology (NIST). Isotopically labeled lipid standards listed in Supplementary Table S5 were used as internal standards, and they were purchased from Avanti Polar Lipids (Alabaster, AL, USA). These standard materials were prepared using optima™ LC–MS-grade methanol, 2-isopropanol (IPA), water, acetonitrile (ACN), ammonium formate, and formic acid from Fisher Chemical (Fisher Scientific, Pittsburgh, PA, USA).

To prepare each solution sample for LC-MS experiments, the lipid internal standards stock solution mixture that were made in concentration listed in Supplementary Table S5, was diluted 60-fold in IPA. A 150 µL aliquot of the diluted lipid internal standards solution was mixed with 50 µL of SRM 1950 human serum, then centrifuged at 12,000 rpm for 5 min. The plasma extracts were transferred to 300 µL LC vials. IPA was used as the blank for all LC-MS analyses.

In the Orbitrap pressure corrected workflow, the diluted lipid internal standard solution was used to determine the exact pressure inside the Orbitrap chamber. The LC eluate was mixed with a continuous stream of the internal standard solution (5 µL min ¹) *via* a T-junction before entering the electrospray ionization (ESI) source.

### LC-MS Experiments

Blanks and SRM 1950 plasma extracts (2 µL injections) were separated on an Accucore™ C_30_ column (150 × 2.1 mm, 2.6 µm particle size). Mobile phase A was 10 mM ammonium formate in water/acetonitrile (40:60 v/v) with 0.1% formic acid, and mobile phase B was 10 mM ammonium formate in 2-isopropanol/acetonitrile (90:10 v/v) with 0.1% formic acid.

For the gradient LC experiment, the elution began at 20% B, ramped to 60% B at 1 min, 70% B at 5 min, 85% B at 5.5 min, 90% B at 8 min, and held at 100% B from 8.2 to 10.5 min, then returned to 20% B at 10.7 min until 12 min total runtime. For isocratic LC experiments, the mobile phase composition was held constant at 85% B for 6 min. The column oven was maintained at 50°C throughout the analysis.

For variable-pressure experiments, the turbopump speed was adjusted between 100% and 66% of the maximum rotational speed. Lower speeds were avoided to prevent the internal vacuum safety system from automatically shutting down the instrument.

Data collection commenced after the UHV pressure stabilized (∼10 min post-adjustment). LC-MS run queues began with solvent blanks followed by full-scan and DDA analyses at specified resolution settings. Full-scan MS data were acquired in positive ion mode from *m/z* 150–2000 at 180,000 FWHM mass resolution unless otherwise specified. Precursor ions were isolated with a 0.8 *m/z* window and fragmented by stepped higher-energy collisional dissociation (HCD) at normalized collision energies of 15%, 30%, 50%. MS/MS spectra were acquired at 15,000 resolution with a fixed duty cycle of 0.6 s.

### Data analysis

Spectral features were extracted using Thermo Scientific Compound Discoverer™ 3.3 SP3 through retention-time alignment, peak picking, feature grouping, integration, and gap filling. Tentative compound annotations were assigned *via* mzCloud and in-house *mzVault* libraries. The processed results were exported as .csv files containing molecular formulas, exact *m/z* values, retention times, and intensities for further processing. The average mass error was 1ppm.

### Data processing

Custom Python scripts were used to calculate the average *self-bunch* mass resolution for each feature using spectral data from the .RAW files. Self-bunching refers to coherent ion motion within the Orbitrap analyzer, which is essential for estimating ion-neutral collision rates. Ions with self-bunched signal-to-noise ratios (SNR) < 30 were excluded. The resolving power of isotopic peaks was averaged to improve statistical robustness within a single MS scan. The monoisotopic resolving power was further averaged over 100 scans centered on the centroid retention time. The average monoisotopic resolving power and ion-gauge pressure readings were used to calculate *^Orbi^CCS_LC_* and *^Orbi^CCS_LC,IS_*using Eqn. (1). The retention time-specific calibration factors, *a_RT_* and *b_RT_*, were derived by linear regression between *^Orbi^CCS_LC,IS_,* and *^DT^CCS_IS_*. The three lipid internal standards LPC 18:1(d7), SM d18:1/18:1(d9), TG15:0/18:1(d7)/15:0 have literature *^DT^CCS* values of 232.9, 286.3, and 312.1 Å^2^, respectively. The *^Orbi^CCS* values of lipid features were calibrated using *a_RT_* and *b_RT_* via Eqn. (2). Features with R² < 0.99 were removed.

### CCS accuracy and precision

We manually compiled *^TIMS^CCS* and *^DT^CCS* values from the literature. To evaluate the accuracy of *^Orbi^CCS*, lipid species annotated in a single LC–Orbitrap experiment were cross-referenced with lipid features in reference CCS databases using *m/z* matching with a mass error of less than 5 ppm. Only lipid features with consistent annotations between the reference databases and the Orbitrap measurements were filtered to evaluate the accuracy of *^Orbi^CCS* values. The final cross-referenced lipid features show a median mass error of ∼0.4 ppm.

To evaluate the precision of *^Orbi^CCS*, data processing was repeated for four independent LC-MS experiments of SRM 1950 plasma. Lipid features with *m/z* errors < 5 ppm (median error 0.1 ppm) and centroid RT ± 0.1s across all experiments were used to calculate the mean and standard deviation of *^Orbi^CCS* values. The list of matched lipid species is summarized in *Supplementary Table S2*.

## Results

### Effect of Pressure Changes during LC-MS Experiments

Although some lipidomics approaches avoid front-end separations, such as those used in shotgun lipidomics^19^, the large majority of experiments make use of ultrahigh performance liquid chromatography (UHPLC) to increase peak capacity and reduce ion suppression before MS detection. UHPLC experiments typically employ solvent gradients to maximize separation efficiency and solve the “general elution problem”. A combination of a heated mass spectrometer inlet system, desolvation gas, and a differentially pumped vacuum system removes the UHPLC solvent following electrospray ionization. Under typical LC-MS conditions, the Orbitrap mass analyzer operates in ultrahigh vacuum (10 ¹¹–10 ¹ mbar), thereby extending ion transient lifetimes and maximizing sensitivity for accurate and precise *m/z* determination. However, if the Orbitrap pressure is purposely elevated by reducing the turbopump speed, ions undergo more frequent collisions with background neutrals. The enhanced ion-neutral collision rate reduces spectral intensity while simultaneously encoding information about the ions’ structure in this environment (**Figure 1a**).

**Figure 1.**
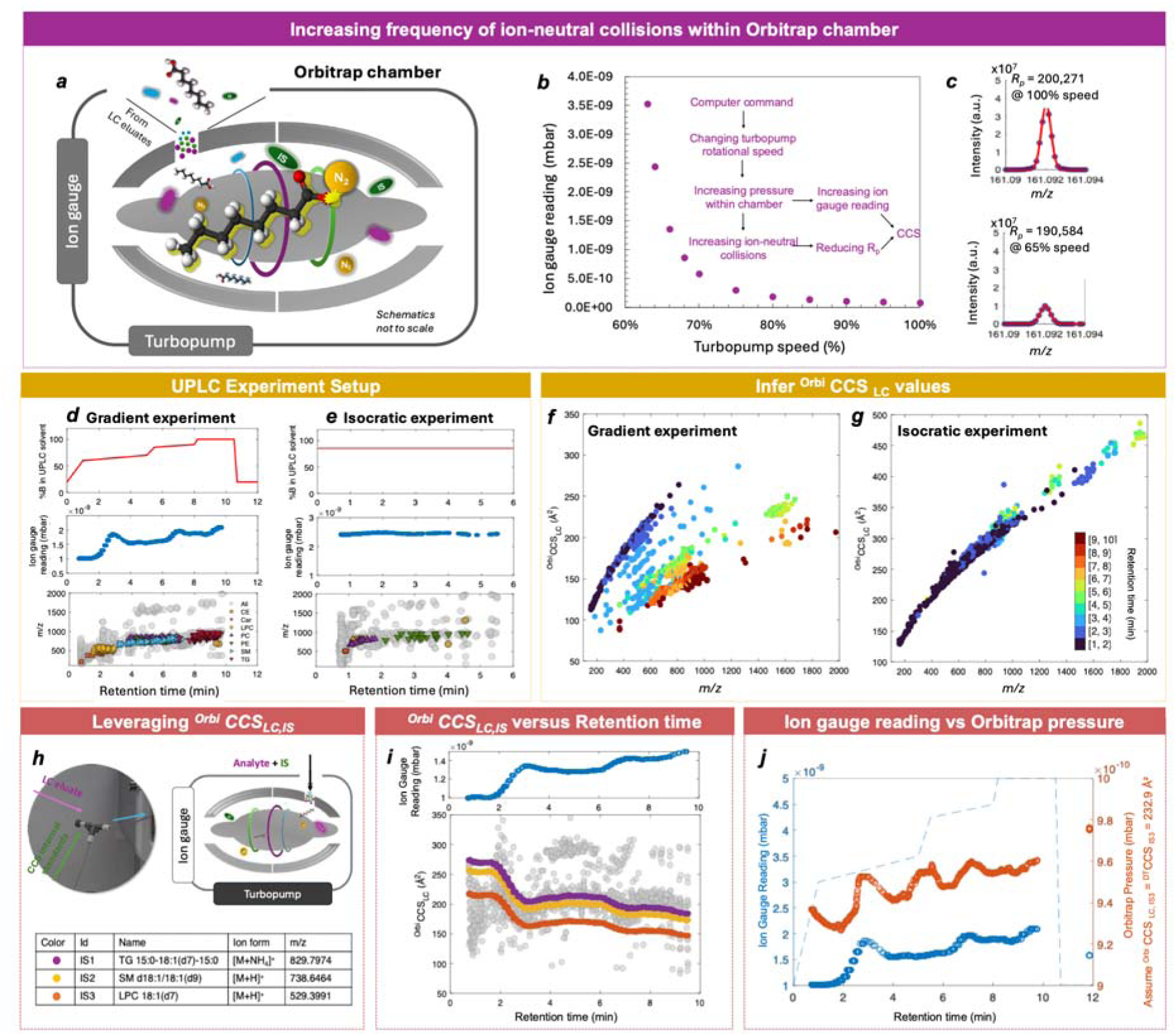
Pressure-dependent ion-neutral collisions and their impact on ^Orbi^CCS_LC_ measurements. (a) Schematic showing ions colliding with background N molecules while oscillating inside the Orbitrap chamber, which is enclosed in a high vacuum chamber. The instrument control software was modified to operate the turbomolecular pump (turbopump) at a user-defined speed. The ion gauge records pressure changes inside the Orbitrap. (b) Relationship between turbopump speed and ion gauge pressure. (c) Zoomed in [Ala +H]⁺ mass spectra from direct infusion (5 μL min⁻¹, 20 μM polyalanine) acquired at 240,000 nominal resolution across several turbopump speeds, showing changes in peak width, which are indicative of greater transient signal decay. LC-MS analysis of SRM 1950 human plasma reference material using a C stationary phase under gradient elution (d) and isocratic conditions at 85% solvent B (e). The red trace denotes the percentage of solvent B (top panel), and the blue dotted line indicates Orbitrap ion gauge pressure as a function of chromatographic retention time (middle panel). Different lipid classes eluted at distinct retention times (bottom panel). Panels (f) and (g) display the calculated *^Orbi^CCS_LC_* values for the lipids detected in the experiments described in (d) and (e), respectively. Data points are color-coded by chromatographic retention time. (h) Post-column infusion of isotopically labeled lipid standards via a T-junction upstream of the ESI ion source. (i) Retention-time-dependent changes in *^Orbi^CCS_LC,IS_*for three internal standards, correlated with ion gauge pressure. (j) Ion gauge pressure (blue) and back-calculated Orbitrap pressure (orange) at 66% turbopump speed, derived from LPC 18:1(d7) using Eqn. (1) (*^DT^CCS_N2_* = 232.9 Å²).

**Figure 1b** shows the effect of reducing the turbopump speed on the measured Orbitrap pressure, reaching 5.07 e-9 mbar (3.8e-9 Torr) at 63% of the maximum rotational speed. Lower speeds were avoided to prevent automatic instrument shutdown triggered by the internal vacuum safety system. **Figure 1c** shows the concomitant reduction in mass resolving power *R_p_* observed for the [Ala_2_+H]^+^ ion during direct infusion experiments (240,000 resolution setting, 75:25 water-acetonitrile solvent composition) conducted at 100% and 65% turbopump speeds. The decreasing *R_p_* indicated an increasing ion-neutral collision rate in the Orbitrap environment, demonstrating that the derived decay constants depend on collisional damping with background neutrals and can therefore be reliably used to calculate CCS values.

In addition to estimating the ion-neutral collision rate from single-peak resolving power (*Rp*) analysis, another critical parameter for deriving *^Orbi^CCS* values via Eqn (1) is the pressure inside the Orbitrap analyzer, *P_Orbi_*. Typically, the pressure inside the Orbitrap analyzer is inferred from the ion gauge reading using a set of external calibrants, assuming a linear, correctable relationship between *P_Orbi_* and *P* during experiments. This method has been used to determine *^Orbi^CCS* values of proteins in previous studies^16,17^. Interestingly, we observed subtle yet measurable pressure fluctuations in the Orbitrap chamber during individual LC runs. **Figure 1d** shows the pressure measured during an LC experiment at a turbopump speed of 66%. Importantly, pressure fluctuations correlated with the percentage of solvent B used in the LC gradient profile, with an approximate 2-minute time delay. The most significant pressure changes occurred when the B rate increased sharply from 20% to 60%. After % B stabilized, the chamber pressure gradually returned to normal under turbopump evacuation. In contrast, no notable pressure fluctuations were observed when conducting LC-MS experiments under isocratic LC conditions with 85% B (**Figure 1e**).

We first calculated lipid CCS values from pressure derived from ion gauge readings, referred to as *^Orbi^CCS_LC_*. As a result of the changing pressure during the LC-MS run, *^Orbi^CCS_LC_* of lipids features showed distinctive traces as a function of retention time (RT) (**Figure 1f**). In contrast, isocratic experiments with constant % B profiles yielded more similar *^Orbi^CCS_LC_*values across all RT at a specific *m/z*, resulting in a more consistent trend line in *^Orbi^CCS_LC_-m/z* space (**Figure 1g**). To further investigate the impact of changing ion gauge readings on *^Orbi^CCS_LC_*values, we merged the LC eluate with a solvent stream containing isotopically labeled lipid standards at a constant flow rate through a T-junction before the ESI ion source (**Figure 1h**). Continuous post-column injection of such standards allowed measuring their *^Orbi^CCS_LC,IS_* values at any given RT (**Figure 1i).** Temporal fluctuations in *^Orbi^CCS_LC,IS_* were systematic and inversely correlated with changes in ion gauge pressure readings. These findings, in contrast to previous assumptions that *P_Orbi_*is directly proportional to *P*, suggested that the LC solvent composition biased ion-gauge pressure readings and ultimately affected the ion-neutral collision rate during the LC experiment.

The apparent pressure inside the Orbitrap analyzer, *P_Orbi_*, can be back-calculated with known *^DT^CCS* values that have been reported in the literature and normalized to standard temperature and pressure conditions (T_o_ = 273.15K, and P_o_ = 1013 mbar) (**Figure 1j**). The back-calculated pressures were consistently lower than the ion gauge readings and showed distinct temporal fluctuation patterns. We note that *P_Orbi_*does not necessarily represent the absolute physical pressure within the Orbitrap analyzer, as the higher-energy *^Orbi^CCS* values of the internal standard would intrinsically differ from those of its low-energy drift-tube CCS counterpart. Nevertheless, the temporal variation in *P_Orbi_* provides a realistic measure of relative pressure fluctuations. *P_Orbi_* values showed exceptional stability, varying by less than ∼2% across LC experiments conducted at 68% turbopump speed. This contrasts with a 100% increase in ion gauge readings. Because ion gauges measure pressure from the ion current generated by filament-induced ionization of background neutrals, introducing volatile LC solvents such as isopropanol can amplify these currents due to their larger ionization cross sections and higher electron densities relative to N_2_. Moreover, these findings suggested that the elution profiles of the LC solvents can bias ion-gauge readings. To address this effect, pressure correction was applied in a retention-time-dependent manner to account for solvent-induced fluctuations, ensuring accurate pressure calibration within the analyzer.

### Pressure-corrected ^Orbi^CCS Workflow

The *^Orbi^CCS* workflow involves LC-MS data acquisition (**Figure 2, step 1)**, lipid feature annotation via Compound Discover (CD) (**step 2**), and computation of average monoisotopic *R_p_* via a custom-built Python script (**step 3.1**). Average monoisotopic *R_p_* is the average *R_p_* of isotopic peaks whose signal-to-noise ratio is above a predefined threshold value. To set up the instrument for continuous injection of internal CCS standards, a T-junction was added upstream of the ESI source, merging the LC eluate with a solvent stream containing isotopically labeled lipid standards (**Figure 1h**). The turbomolecular pump was set to the desired rotational speed prior to data acquisition. We note that the impact of pressure fluctuations on *^Orbi^CCS_LC_* values is more pronounced for ions with higher *m/z*. To better account for this *m/z*-dependent effect, we implemented a strategy to calibrate *^Orbi^CCS_LC_*values for detected lipids using three internal standards spanning *m/z* 500-900 Th. To measure the *^Orbi^CCS_LC_ value* of a selected lipid feature, the monoisotopic resolving power was calculated and averaged across 100 scans centered on the feature’s centroid retention time, yielding a 0.2-s temporal window (256 ms per scan at a resolution of 120,000). The *^Orbi^CCS_LC_*values of internal standards (*^Orbi^CCS_LC,IS_*) were then calculated from the same 100 scans within the same temporal window (**Figure 2, step 3.2**). Linear regression analysis between *^DT^CCS* values of internal standards and their corresponding *^Orbi^CCS_LC,IS_*was performed to derive RT-specific calibration factors, *a_RT_* and *b_RT_* (**Figure 2, step 3.3**). The values of *a_RT_* are highly correlated with the ion gauge reading, whereas the values of *b_RT_* are less so (**Supplementary Figure S6).** These calibration factors were then used to determine *^Orbi^CCS* values of the corresponding lipid feature, following:

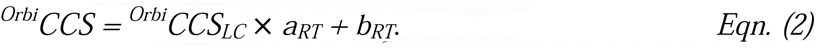

**Figure 2.**
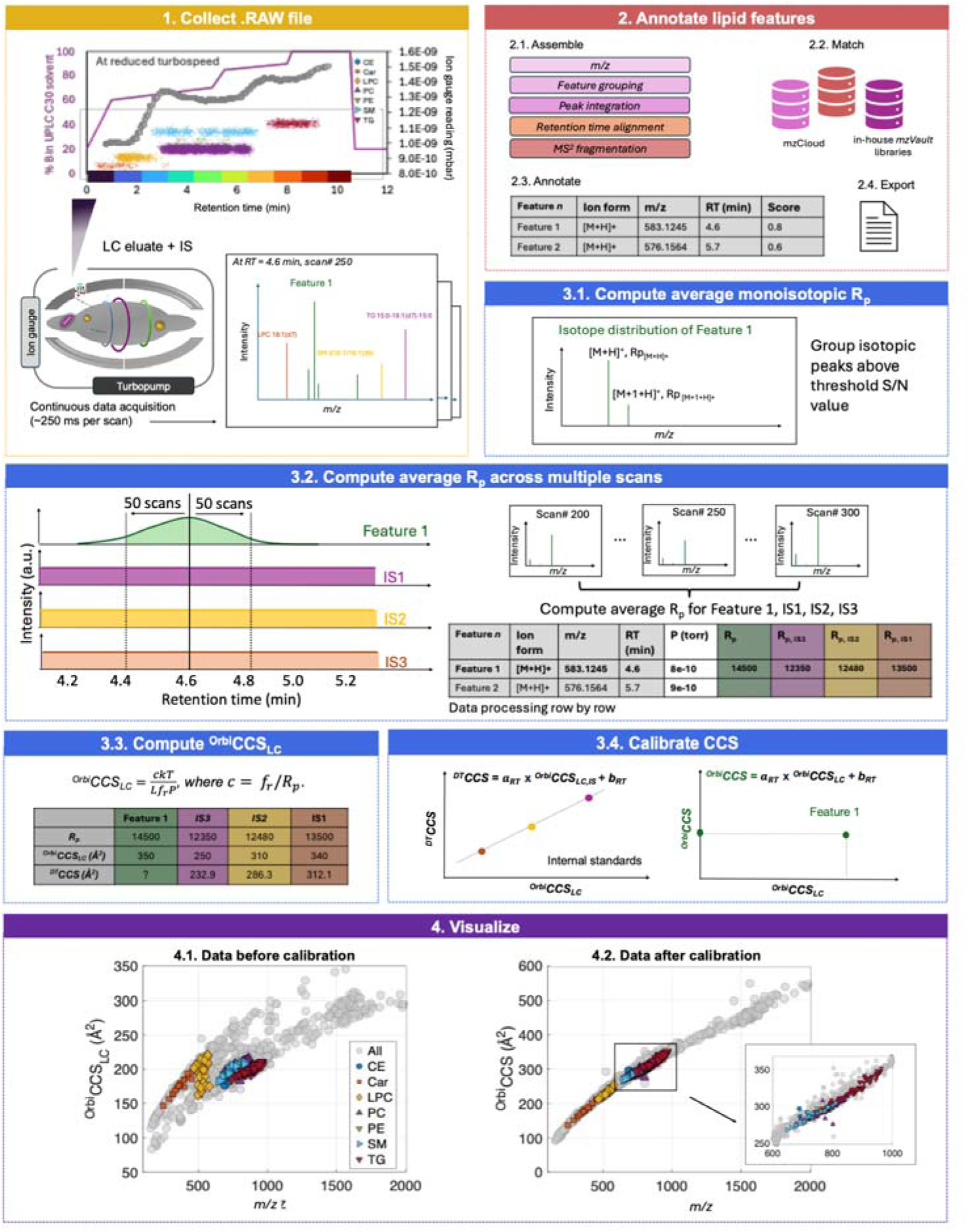
Workflow for ^Orbi^CCS Calculation. The ^Orbi^CCS workflow comprises data acquisition (step 1), lipid feature annotation using Compound Discoverer (CD, step 2), and post-processing with custom Python scripts (step 3). Step 3.1: The resolving power of isotopic peaks with signal-to-noise ratios above a defined threshold was identified and averaged to improve statistical robustness. Step 3.2: For each lipid feature, the monoisotopic resolving power was averaged over 100 scans centered on the feature’s centroid retention time, corresponding to a 0.2 s temporal window (256 ms per scan at a resolution of 120,000). Step 3.3: *^Orbi^CCS_LC_*values for the internal standards (LPC 18:1(d7), SM d18:1/18:1(d9), and TG 15:0/18:1(d7)/15:0) were calculated over the same scan window. Step 3.4: Linear regression of reference *^DT^CCS* values for the internal standards against their corresponding *^Orbi^CCS_LC_*values was used to derive retention-time-specific calibration factors *a_RT_* and *b_RT_*, which were applied to determine calibrated *^Orbi^CCS* values. Step 4: Visualization of annotated lipids and unannotated features in *^Orbi^CCS* space before (4.1) and after (4.2) CCS calibration.

As a result of this retention-time-specific calibration, *^Orbi^CCS* values were significantly compressed in *m/z*-CCS space compared with *^Orbi^CCS_LC_*(**Figure 2, step 4**). However, CCS separation of isobaric lipid species was still observed.

### Evaluating ^Orbi^CCS Accuracy and Precision

Using published results, we created an in-house database of drift tube IM CCS values (*^DT^CCS*) and trapped IM CCS values (*^TIMS^CCS*) for the lipids in SRM 1950 that were annotated at a Schymanski confidence level 3^20^. To evaluate the accuracy of *^Orbi^CCS*, lipid species detected in a single LC–Orbitrap experiment (pump speed 68% and resolution setting 240,000) were cross-referenced with lipid features in reference CCS databases using *m/z* values within 5 ppm. Only lipid features with consistent annotations across the reference databases and the Orbitrap measurements were retained for *^Orbi^CCS* accuracy evaluation (**Supplementary Tables S3 and S4**). The final set of cross-referenced lipid features exhibited a median mass error of <1 ppm. Matched species showed an average *^Orbi^CCS* deviation of 0.1% relative to *^TIMS^CCS* and -0.6% relative to *^DT^CCS* (**Figure 3a and d**). Lipid features at *m/z* < 500 exhibited the most significant deviation from reference CCS values (up to -10%). This mass range lies outside the selected internal standards and covers most Car lipids and some LPC lipids. A breakdown of matched lipid species across lipid classes revealed a class bias, which may reflect fundamental structural differences in lipid head groups in the environment where CCS values were determined (**Figure 3b and e**). Greater than 90% of matched features had less than ±5% error relative to reference values (**Figure 3c and f**). Accuracy was comparable across experiments conducted at different turbopump speeds, with results within 1% of the *^TIMS^CCS* reference values and 2% of the *^DT^CCS* reference values. The number of annotated features decreased slightly at lower turbopump speeds (**Figure 3g**). Experiments conducted at higher resolution settings showed higher accuracy and more annotated features (**Figure 3h**).

**Figure 3.**
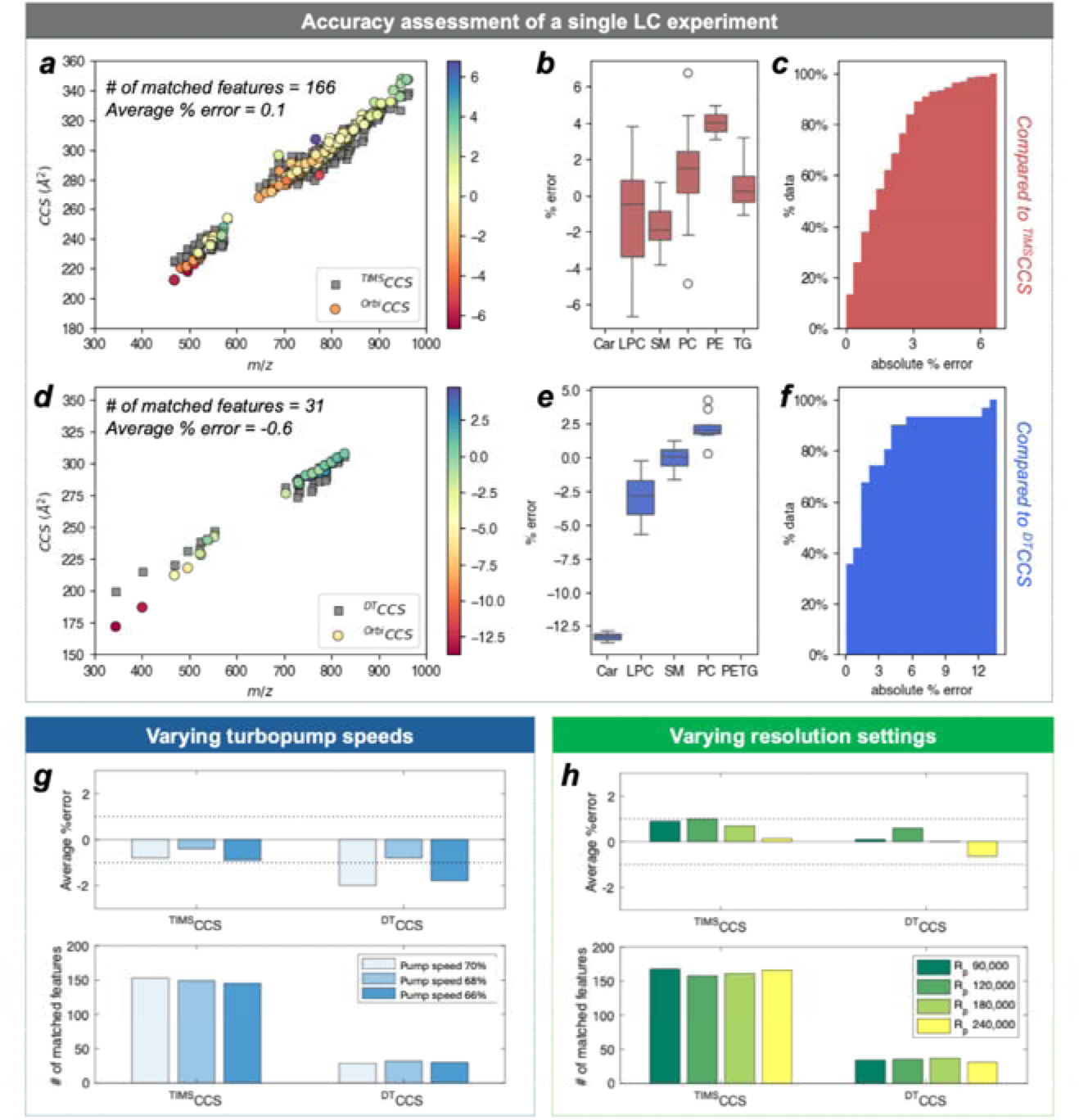
Evaluation of *^Orbi^CCS* Accuracy. (a-f) Comparison of *^Orbi^CCS* values to reference trapped IM (TIMS) collision cross section values *(^TIMS^CCS*) and drift tube (DT) IM collision cross section values (*^DT^CCS*). Annotated lipid species in LC-Orbitrap experiments were cross-referenced to CCS reference databases by matching *m/z* within 5 ppm error and checking for consistent lipid annotation. The final matched datasets show a median mass error of <1 ppm. The number of matched species and the mean percentage CCS error are indicated. (b, e) Average percentage errors of *^Orbi^CCS* values grouped by lipid class relative to *^TIMS^CCS* and *^DT^CCS* reference datasets. (c, f) Cumulative percentage of matched lipid species within specific error thresholds.

Lipid *^Orbi^CCS* values were highly reproducible with an average of 0.7 % (*RSD*) based on four independent LC experiments (**Figure 4a**; **Supplementary Table S2**). More than 90% of annotated lipid features had *RSD* <1% (**Figure 4c**). In particular, LPC, SM, and TG lipids showed median RSD < 0.5% (**Figure 4d-g**). No clear correlation was observed between measured RSD and the magnitude of *^Orbi^CCS* values, the number of unsaturations, or the number of carbon atoms in the sidechain. PC and PE lipids had the highest RSD, likely due to their significant overlap in RT and *m/z* values.

**Figure 4.**
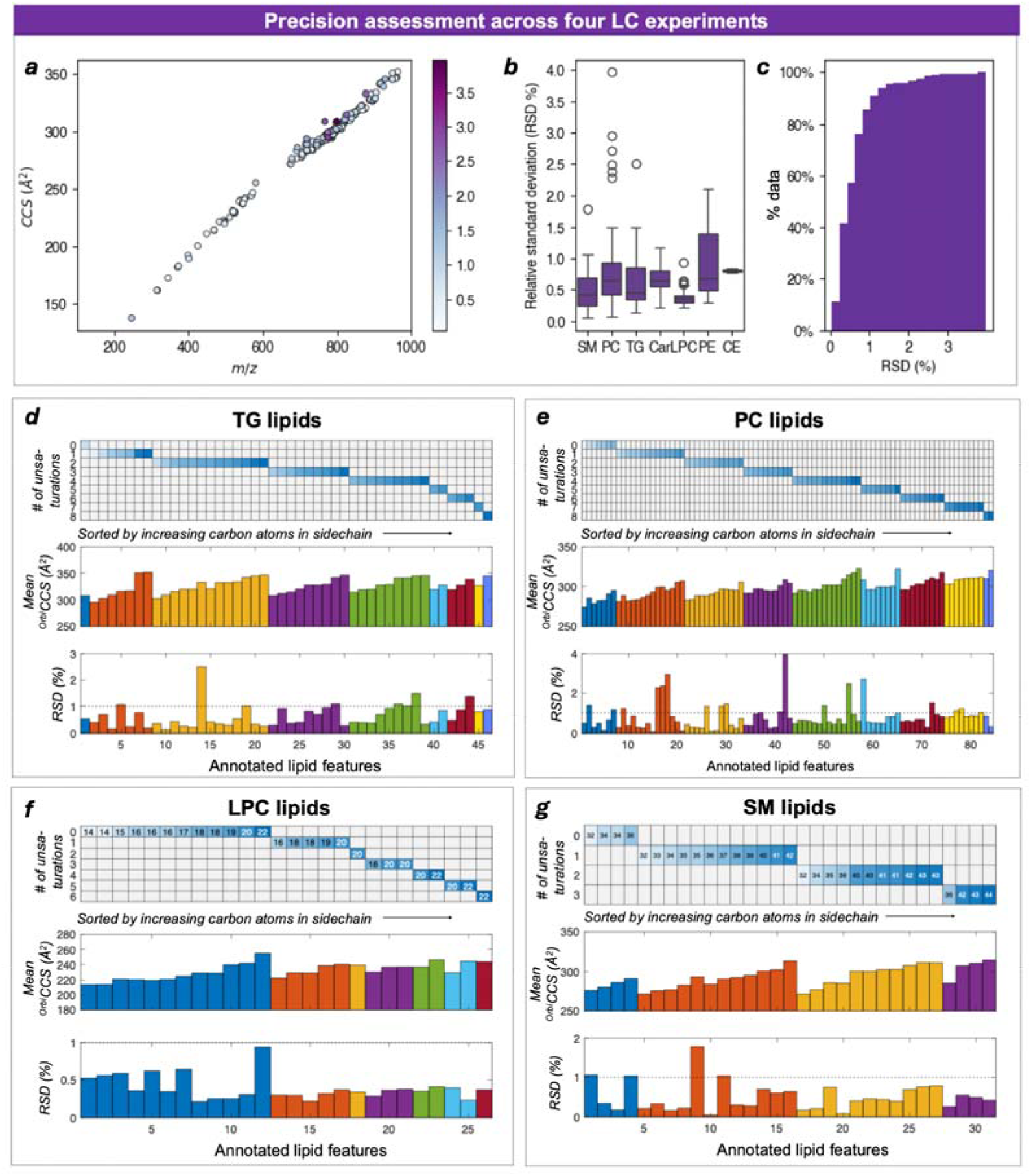
Evaluation of *^Orbi^CCS* Precision Across Multiple LC Experiments. (a) Reproducibility of *^Orbi^CCS* measurements expressed as relative standard deviation (RSD). (b). Calculated RSD broken down by lipid class. (c). Cumulative percentage of lipid species within specific RSD thresholds. Panels d-g summarize the mean *^Orbi^CCS* values and corresponding RSD of annotated lipid features grouped by lipid classes, TG, PC, LPC, and SM.

## Discussion

Over 2,000 features were detected in a single LC-Orbitrap experiment (turbopump speed at 68% and resolution set to 240,000). As expected, the chemical space was crowded when viewed in a single dimension (*m/z* or retention time) or even in two dimensions (*m/z* versus RT) (**Figure 5a**). Leveraging MS/MS information allowed the annotation of ∼20% of the detected lipids, leaving many features unassigned, as is typically the case with lipidomic datasets. Using the workflow presented in this study, we computed *^Orbi^CCS* values for all detected lipid features that exceeded the S/N threshold in the LC-MS run, thereby providing an additional dimension for feature annotation and decluttering spectral space. We cross-referenced *^Orbi^CCS* values against a *m/z*-matched CCS reference database, successfully matching 31 lipid species against *^DT^CCS* (*n*=921) and 166 lipid species against *^TIMS^CCS* (*n*=326) databases, with a maximum mass tolerance of 5 ppm (<1 ppm median mass error, **Figure 5b**). Among these matched species, 12 and 60 lipid species were matched with <1% difference in CCS values to *^DT^CCS* and *^TIMS^CCS* databases, respectively (***Supplementary Table S7&8)***. The number of annotations was constrained by the limited number of CCS values in the CCS reference databases used. However, with the continued expansion of reference CCS datasets, the coverage of *^Orbi^CCS*-based annotations could scale substantially. Thus, we expect that integrating *^Orbi^CCS* into annotation workflows will provide a powerful complementary metric for validating and refining lipid feature assignments across diverse databases.

**Figure 5.**
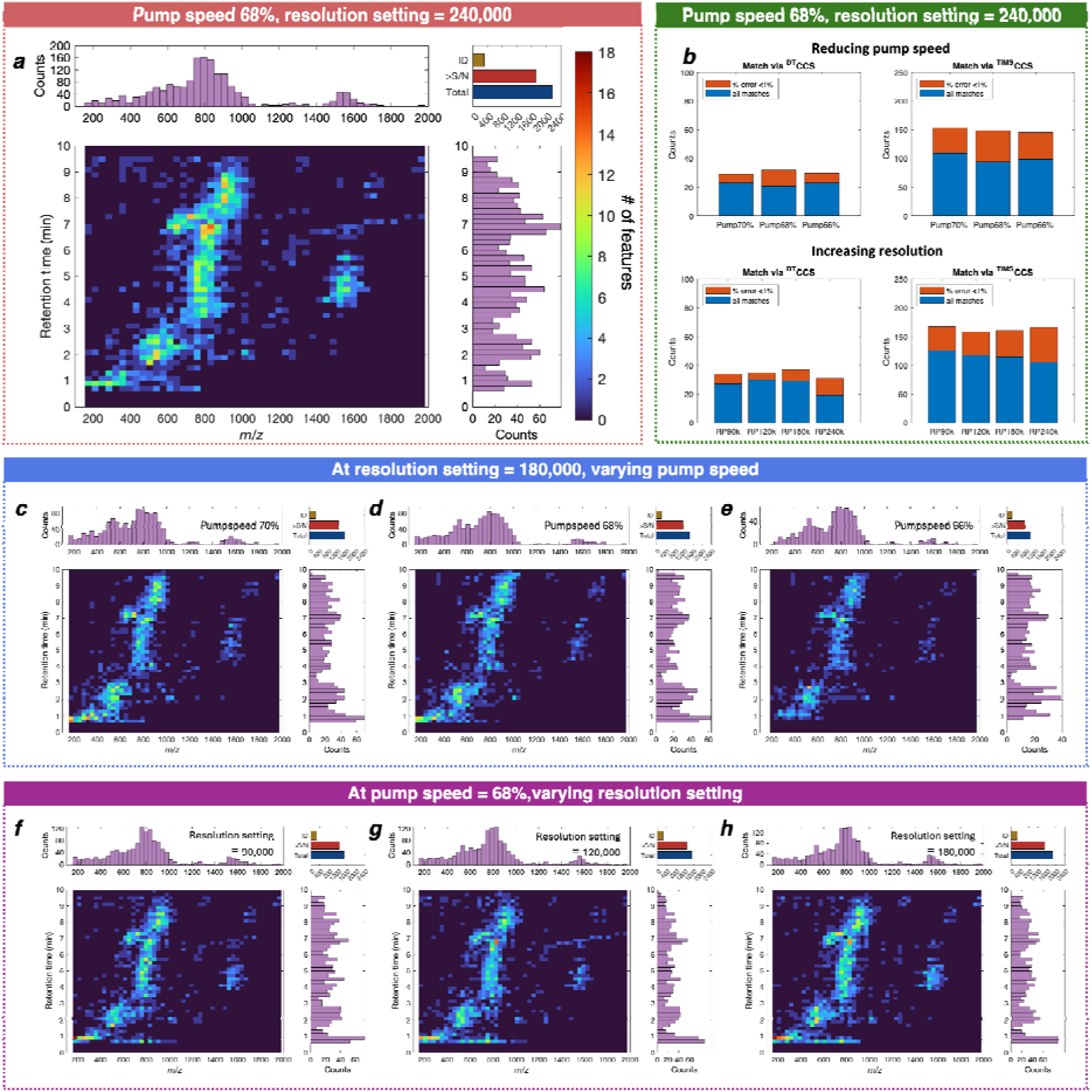
Impact of Turbopump speed and Orbitrap Resolution Settings on Annotations. (a) Heatmap showing the number of detected features by *m/z* and retention time in a single UHPLC-Orbitrap experiment (turbopump speed 68% and resolution 240,000). Histograms summarize feature counts by *m/z* and retention time. The inset bar graph reports the total number of detected features, the number of features above threshold S/N for ^Orbi^CCS calculation, and the number of features that were annotated at a Schymanski confidence level 3 (see^20^). (b) Bar graphs summarizing the total number of lipid features matched relative to reference CCS datasets (blue) based on *m/z* and high-confidence matches with <1% CCS error (orange), across different turbopump speeds and resolution settings. The bar graphs summarize these statistics in experiments conducted at different turbopump speeds and resolution settings. (c-h). Heatmaps as in panel (a) for single UHPLC-Orbitrap experiments acquired at different turbopump speeds and resolution settings.

We investigated how various experimental settings affected the number of *^Orbi^CCS* values successfully matched to reference CCS databases. **Figure 5b** summarizes the number of lipid annotations at different turbopump speeds and Orbitrap resolution settings, highlighting that differences were relatively minimal in the tested range. The feature count histograms showed that these minor feature losses occurred mainly in the *m/z* < 300 and *m/z* >1,400 regions (**Figure 5c-e**). These ions have either a higher axial oscillatory frequency or a higher CCS value. In contrast, as experimental resolution setting increased, more features were detected and annotated at a given turbo pump speed (**Figure f-h)**.

Beyond database matching, measuring *^Orbi^CCS* values enabled high-level interrogation of complex mixtures by mapping each lipid feature into a multidimensional space defined by RT, *m/z*, and *^Orbi^CCS*, thereby revealing local trends among structurally related lipids. Contextual structural relationships provide additional information to confirm lipid species annotations in complex mixtures. A case study is discussed here to exemplify how related lipid classes behave with respect to *^Orbi^CCS* in an Orbitrap LC-MS experiment (**Figure 6a**). PC lipids with different numbers of unsaturation sites or acyl chain carbon atoms showed distinct trendlines in *m/z*-RT space (**Figure 6a**). Deviation from such trendlines may indicate a variety of lipid modifications such as alternative adduct formation, oxidation, or related isomers. Oxidation of PC lipids, for example, tends to move the trendline towards lower *m/z* and higher RT. Sodiated adducts deviate from the general trendline, characterized by higher *^Orbi^CCS* values, longer retention times, and higher *m/z* values (**Figure 6a and 6b**). Lipid isomers can be recognized by their distinct RT but identical *m/z* values and are characterized by horizontal alignment in the *m/z*-RT plot (**Figure 6b**). Relative differences in *^Orbi^CCS* between isomers provide insights into their structures, aiding in isomer assignment. For example, three PC features were annotated as PC(36:2) in the narrow 770-810 Th *m/z* range and 4.9–5.2 min retention time window (**Figure 6c**). Three out of four LC-MS experiments showed that the later-eluting isomer had a larger *^Orbi^CCS* value than the other two species (***Supplementary Table S2***). Previous TIMS analysis of PC(36:2) isomers has shown that the later-eluting isomer is likely PC(18:1(6z)/18:1(6z)) with a larger *^TIMS^CCS* value, consistent with the observed *^Orbi^CCS* values and RT in this study (**Figure 6d** ^21^). Additionally, the *^Orbi^CCS* values for these PC(36:2) isomers matched the *^TIMS^CCS* measurements with an error of 1.5%. The remaining two [M+H]^+^ PC(36:2) isomers were indistinguishable by *^Orbi^CCS*, considering the observed %RSD in the experiments. Overall, these results exemplify how the relative positions of lipid features within the combined *^Orbi^CCS*-RT-*m/z* space can aid in identifying isomeric lipid species and how *^Orbi^CCS* adds a new molecular metric that can be used in LC-MS lipidomics measurements to better characterize datasets and improve annotation confidence.

**Figure 6.**
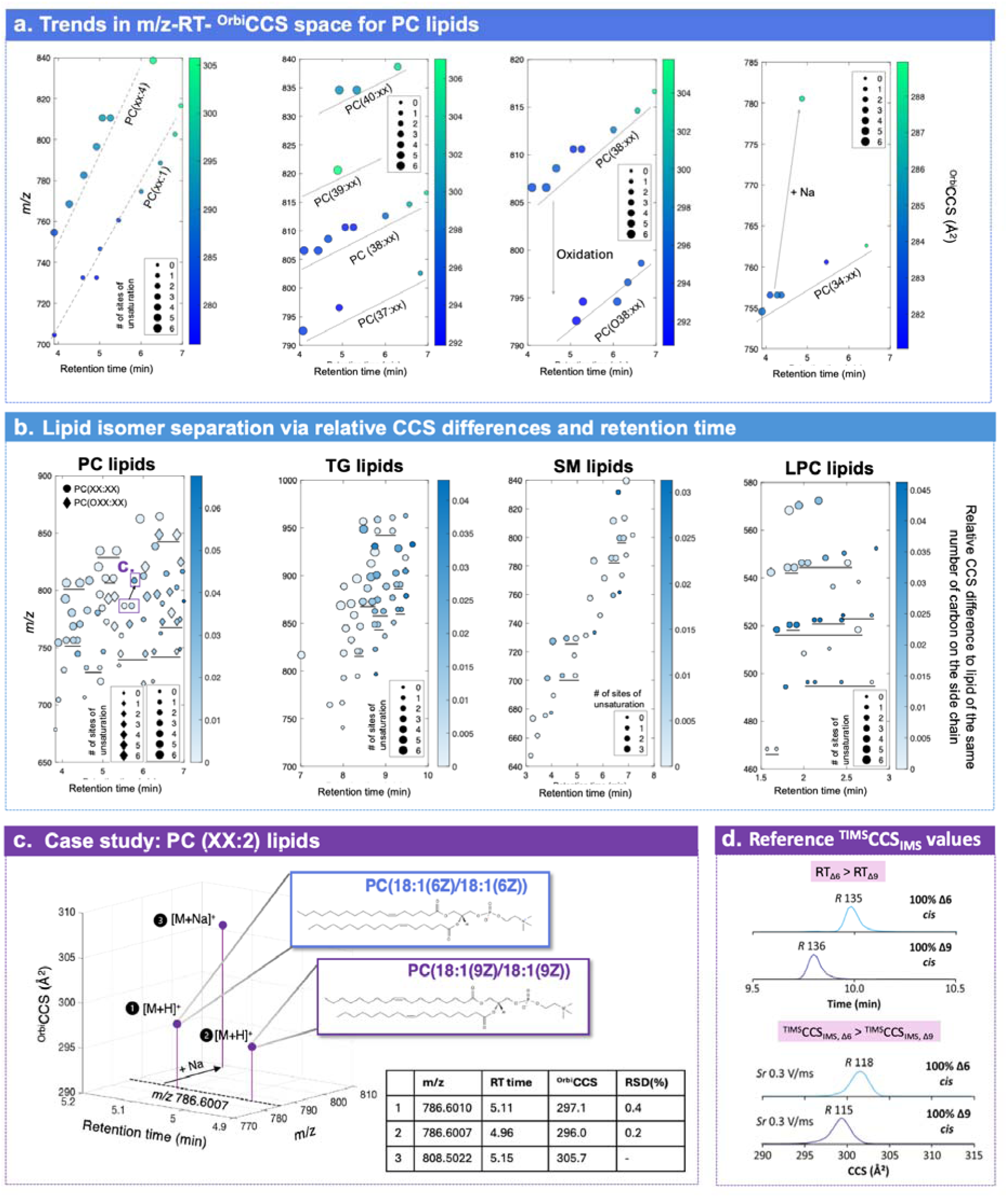
Elucidation of lipid structures using contextual RT-*m/z*–*^Orbi^CCS* relationships. Case study demonstrating observed trends in lipid structures within a single LC-MS experiment using complementary *m/z*, RT, and *^Orbi^CCS* information. (a) RT-*m/z* scatter plot of phosphatidylcholine (PC) lipids. The marker size represents the number of unsaturations, and color encodes *^Orbi^CCS* values. Systematic trendlines emerge as functions of the number of unsaturations and the carbon number of the acyl chains. Deviations from trendlines indicate structural variations such as oxidation, adduct formation, or isomerism. (b) Similar structural trendlines were observed in scatter plots of annotated PC, SM, LPC, and TG lipids. The marker size indicates the number of unsaturations. The color scale shows the relative OrbiCCS values for lipid features with the same number of carbon atoms. Isomers were underscored in the scatter plots. (c) Expanded view of a narrow RT (4.9–5.2 min) and *m/z* (770–810 Th) window highlighting three features annotated as PC(36:2), including two [M+H] isomers with identical *m/z* but distinct RT. (d) Comparison of *^Orbi^CCS* values for the two PC(36:2) isomers across replicate LC experiments showing the later-eluting isomer has larger *^Orbi^CCS*, consistent with reported *^TIMS^CCS* values and literature RT assignments.

## Supporting Information

See PDF attached.

## Acknowledgements

FMF and Z.N. acknowledge support by NIH R01CA218664, NIH R61CA281667, and the Georgia Institute of Technology. We also acknowledge support by the Petit Institute of Bioengineering and Bioscience Systems’ Mass Spectrometry Core Facility.

## Author Contributions

Z.N. and F.M.F designed the study. Z.N. and S.M. prepared the samples. K.F. and A.M. set up the instrument to run at reduced turbopump speeds. S.M. and D.G. contributed to optimizing the mass spectrometer parameters. Z.N. performed most of the experiments and analyzed the data. K.A. contributed to the initial Python script for data processing. All authors contributed to the discussion. Z.N. and F.M.F. wrote the manuscript. All authors discussed the results and commented on the manuscript.

