## Supplementary for "Orbitrap Collision Cross Section Measurements Enhance Isomer Annotations in Lipidomics": OrbiCCS_SI_070326.pdf

### Table of Contents

- Supplementary Table S1. Orbitrap resolution settings and corresponding transient durations.
- Supplementary Table S2. Summary of $^{Orbi}$CCS values for 202 annotated lipid species in SRM1950 plasma.
- Supplementary Table S3. Summary of lipid features with matched $^{Orbi}$CCS and $^{TIMS}$CCS values from a single LC-Orbitrap experiment (pump speed 68%, resolution setting 240,000). The same dataset present in Fig. 3a-f.
- Supplementary Table S4. Summary of lipid features with matched $^{Orbi}$CCS and $^{DT}$CCS values from a single LC-Orbitrap experiment (pump speed 68%, resolution setting 240,000). The same dataset present in Fig. 3a-f.
- Supplementary Table S5. Isotopically labeled lipid standards.
- Supplementary Figure S6. Slope and intercept values used to calibrate $^{Orbi}$CCS during LC-MS experiments.
- Supplementary Table S7. Summary of lipid features with matched $^{Orbi}$CCS to reference $^{TIMS}$CCS dataset for experiments conducted at a resolution of 240,000 and turbopump speed 68%.
- Supplementary Table S8. Summary of lipid features with matched $^{Orbi}$CCS to reference $^{DT}$CCS dataset for experiments conducted at a resolution of 240,000 and turbopump speed 68%.
- Supplementary Table S9. Summary of lipid features with matched $^{Orbi}$CCS to reference $^{TIMS}$CCS dataset for experiment conducted at resolution 180,000 and turbopump speed 68%.

- Supplementary Table S10. Summary of lipid features with matched ${}^{Orbi}CCS$ to reference ${}^{DT}CCS$ dataset for experiments conducted at a resolution of 180,000 and turbopump speed 68%.
- Supplementary Table S11. Summary of lipid features with matched ${}^{Orbi}CCS$ to reference ${}^{TIMS}CCS$ dataset for experiments conducted at a resolution of 120,000 and turbopump speed 68%.
- Supplementary Table S12. Summary of lipid features with matched ${}^{Orbi}CCS$ to reference ${}^{DT}CCS$ dataset for experiments conducted at a resolution of 120,000 and turbopump speed 68%.
- Supplementary Table S13. Summary of lipid features with matched ${}^{Orbi}CCS$ to reference ${}^{TIMS}CCS$ dataset for experiments conducted at a resolution of 90,000 and turbopump speed 68%.
- Supplementary Table S14. Summary of lipid features with matched ${}^{Orbi}CCS$ to reference ${}^{DT}CCS$ dataset for experiments conducted at a resolution of 90,000 and turbopump speed 68%.

**Supplementary Table S1. Orbitrap resolution settings and corresponding transient durations.** Summary of resolution settings used on the Thermo Fisher Orbitrap Exploris™ 240 mass spectrometer and the corresponding transient acquisition durations for a single scan.

| **Resolution setting at *m/z* 200** | **Duration of transient signal (ms)** |
| --- | --- |
| 60,000 | 128 |
| 90,000 | 192 |
| 120,000 | 256 |
| 180,000 | 384 |
| 240,000 | 512 |

**Supplementary Table S2**. **Summary of $^{Orbi}$CCS of 202 annotated lipid species in IPA extract of SRM1950 plasma.** Table reports lipid class, proposed annotation, experimental monoisotopic m/z value, retention time, $^{Orbi}$CCS values, and relevant statistics across four LC experiments conducted at 68% turbopump speed. Annotation was assigned based on the following criteria: 1) $MS^1$ and MS/MS spectra of the standard matched to the compound. 2) MS1 and MS/MS spectrum of the feature matched with library spectra.

| # | Name | Formula | *m/z* | Type | RT1 | RT2 | RT3 | RT4 | CCS1 | CCS2 | CCS3 | CCS4 | meanCCS | stdCCS | RSD(%) |
|---|---|---|---|---|---|---|---|---|---|---|---|---|---|---|---|
| 1 | Car(5:0) | C12 H23 N O4 | 246.1699 | Car | 0.82 | 0.81 | 0.84 | 0.83 | 139.2 | 136.2 | 135.5 | 136.4 | 136.8 | 1.6 | 1.2 |
| 2 | Car(10:0-OH) | C17 H33 N O5 | 314.2326 | Car | 0.92 | 0.92 | 0.93 | 0.94 | 163.0 | 159.7 | 161.2 | 160.9 | 161.2 | 1.3 | 0.8 |
| 3 | Car(10:0) | C17 H33 N O4 | 316.2482 | Car | 0.98 | 0.98 | 1.00 | 1.01 | 163.0 | 159.8 | 161.0 | 160.8 | 161.2 | 1.3 | 0.8 |
| 4 | Car(12:0) | C19 H37 N O4 | 344.2795 | Car | 1.21 | 1.21 | 1.22 | 1.23 | 172.5 | 171.6 | 171.8 | 172.1 | 172.0 | 0.4 | 0.2 |
| 5 | Car(14:1) | C21 H39 N O4 | 370.2951 | Car | 1.37 | 1.36 | 1.38 | 1.39 | 182.2 | 181.2 | 182.3 | 180.1 | 181.4 | 1.1 | 0.6 |
| 6 | Car(14:0) | C21 H41 N O4 | 372.3108 | Car | 1.60 | 1.59 | 1.63 | 1.65 | 183.6 | 182.8 | 183.1 | 180.9 | 182.6 | 1.2 | 0.6 |
| 7 | Car(16:1) | C23 H43 N O4 | 398.3265 | Car | 1.72 | 1.71 | 1.76 | 1.78 | 192.8 | 193.0 | 191.8 | 190.3 | 192.0 | 1.3 | 0.7 |
| 8 | Car(16:0) | C23 H45 N O4 | 400.3421 | Car | 2.01 | 2.01 | 2.04 | 2.05 | 190.7 | 188.6 | 189.7 | 187.2 | 189.1 | 1.5 | 0.8 |
| 9 | Car(18:2) | C25 H45 N O4 | 424.3422 | Car | 1.82 | 1.82 | 1.87 | 1.88 | 201.7 | 199.4 | 200.1 | 199.4 | 200.2 | 1.1 | 0.5 |
| 10 | Car(20:4) | C27 H45 N O4 | 448.3421 | Car | 1.79 | 1.79 | 1.84 | 1.86 | 211.0 | 211.3 | 210.8 | 210.3 | 210.8 | 0.4 | 0.2 |
| 11 | CE(20:4)+NH3 | C47 H79 N O2 | 690.6185 | CE | 8.91 | 8.91 | 8.91 | 8.92 | 281.8 | 280.7 | 283.8 | 285.7 | 283.0 | 2.2 | 0.8 |
| 12 | CE(22:6)+NH3 | C49 H79 N O2 | 714.6188 | CE | 8.70 | 8.70 | 8.70 | 8.70 | 286.8 | 287.3 | 291.4 | 291.1 | 289.2 | 2.4 | 0.8 |
| 13 | LPC(14:0)>LPC(14:0/0:0) _and_LPC(0:0/14:0) | C22 H46 N O7 P | 468.3085 | LPC | 1.62 | 1.62 | 1.66 | 1.67 | 215.1 | 214.2 | 213.9 | 212.4 | 213.9 | 1.1 | 0.5 |
| 14 | LPC(14:0)>LPC(14:0/0:0) _and_LPC(0:0/14:0) | C22 H46 N O7 P | 468.3085 | LPC | 1.52 | 1.52 | 1.55 | 1.57 | 215.4 | 214.5 | 214.0 | 212.5 | 214.1 | 1.2 | 0.6 |
| 15 | LPC(15:0)>LPC(15:0/0:0) _and_LPC(0:0/15:0) | C23 H48 N O7 P | 482.3242 | LPC | 1.76 | 1.76 | 1.81 | 1.82 | 220.9 | 220.7 | 223.3 | 220.4 | 221.3 | 1.3 | 0.6 |
| 16 | LPC(16:1)>LPC(16:1/0:0) _and_LPC(0:0/16:1) | C24 H48 N O7 P | 494.3242 | LPC | 1.64 | 1.64 | 1.68 | 1.69 | 223.3 | 222.1 | 223.3 | 222.1 | 222.7 | 0.7 | 0.3 |
| 17 | LPC(16:0)>LPC(16:0/0:0) _and_LPC(0:0/16:0) | C24 H50 N O7 P | 496.3396 | LPC | 2.08 | 2.07 | 2.10 | 2.11 | 221.7 | 220.5 | 221.3 | 219.9 | 220.8 | 0.8 | 0.4 |
| 18 | LPC(16:0)>LPC(16:0/0:0) _and_LPC(0:0/16:0) | C24 H50 N O7 P | 496.3398 | LPC | 2.76 | 2.76 | 2.77 | 2.77 | 221.2 | 220.7 | 219.3 | 218.2 | 219.8 | 1.4 | 0.6 |
| 19 | LPC(16:0)>LPC(16:0/0:0) _and_LPC(0:0/16:0) | C24 H50 N O7 P | 496.3399 | LPC | 1.99 | 1.99 | 2.02 | 2.03 | 221.7 | 220.4 | 221.8 | 220.5 | 221.1 | 0.8 | 0.3 |
| 20 | LPC(17:0)>LPC(17:0/0:0) _and_LPC(0:0/17:0) | C25 H52 N O7 P | 510.3555 | LPC | 2.25 | 2.25 | 2.26 | 2.22 | 226.4 | 225.4 | 225.9 | 223.1 | 225.2 | 1.5 | 0.6 |
| 21 | LPC(18:3)>LPC(18:3/0:0) _and_LPC(0:0/18:3) | C26 H48 N O7 P | 518.3242 | LPC | 1.64 | 1.64 | 1.68 | 1.65 | 230.4 | 230.0 | 231.6 | 230.8 | 230.7 | 0.7 | 0.3 |
| 22 | LPC(20:5)>LPC(20:5/0:0) _and_LPC(0:0/20:5) | C28 H48 N O7 P | 520.3397 | LPC | 1.86 | 1.86 | 1.82 | 1.84 | 229.3 | 229.2 | 231.2 | 230.1 | 230.0 | 0.9 | 0.4 |
| 23 | LPC(18:1)>LPC(18:1/0:0) _and_LPC(0:0/18:1) | C26 H52 N O7 P | 522.3552 | LPC | 2.07 | 2.07 | 2.09 | 2.11 | 229.2 | 229.2 | 230.7 | 229.7 | 229.7 | 0.7 | 0.3 |
| 24 | LPC(18:1)>LPC(18:1/0:0) _and_LPC(0:0/18:1) | C26 H52 N O7 P | 522.3554 | LPC | 2.14 | 2.14 | 2.16 | 2.17 | 229.2 | 229.4 | 230.3 | 229.3 | 229.5 | 0.5 | 0.2 |
| 25 | LPC(18:0)>LPC(18:0/0:0) _and_LPC(0:0/18:0) | C26 H54 N O7 P | 524.3710 | LPC | 2.39 | 2.40 | 2.40 | 2.40 | 229.2 | 229.8 | 229.8 | 228.9 | 229.4 | 0.5 | 0.2 |
| 26 | LPC(18:0)>LPC(18:0/0:0) _and_LPC(0:0/18:0) | C26 H54 N O7 P | 524.3713 | LPC | 2.76 | 2.73 | 2.77 | 2.77 | 229.3 | 229.9 | 229.2 | 228.4 | 229.2 | 0.6 | 0.3 |
| 27 | LPC(19:1)>LPC(19:1/0:0) _and_LPC(0:0/19:1) | C27 H54 N O7 P | 536.3712 | LPC | 2.29 | 2.29 | 2.30 | 2.31 | 238.2 | 237.6 | 239.2 | 239.0 | 238.5 | 0.8 | 0.3 |

| # | Name | Formula | *m/z* | Type | RT1 | RT2 | RT3 | RT4 | CCS1 | CCS2 | CCS3 | CCS4 | meanCCS | stdCCS | RSD(%) |
|---|---|---|---|---|---|---|---|---|---|---|---|---|---|---|---|
| 28 | LPC(19:0)>LPC(19:0/0:0) _and_LPC(0:0/19:0) | C27 H56 N O7 P | 538.3870 | LPC | 2.55 | 2.54 | 2.55 | 2.54 | 240.1 | 239.8 | 238.8 | 240.0 | 239.7 | 0.6 | 0.3 |
| 29 | LPC(20:4)>LPC(20:4/0:0) _and_LPC(0:0/20:4) | C28 H50 N O7 P | 544.3398 | LPC | 1.84 | 1.84 | 1.81 | 1.82 | 237.1 | 235.8 | 237.6 | 236.1 | 236.7 | 0.8 | 0.4 |
| 30 | LPC(20:3)>LPC(20:3/0:0) _and_LPC(0:0/20:3) | C28 H52 N O7 P | 546.3555 | LPC | 2.00 | 2.09 | 2.03 | 2.04 | 237.3 | 235.9 | 237.3 | 235.7 | 236.6 | 0.9 | 0.4 |
| 31 | LPC(20:3)>LPC(20:3/0:0) _and_LPC(0:0/20:3) | C28 H52 N O7 P | 546.3558 | LPC | 1.93 | 1.93 | 1.96 | 1.98 | 237.6 | 236.1 | 237.7 | 236.2 | 236.9 | 0.9 | 0.4 |
| 32 | LPC(20:2)>LPC(20:2/0:0) _and_LPC(0:0/20:2) | C28 H54 N O7 P | 548.3713 | LPC | 2.21 | 2.21 | 2.23 | 2.23 | 240.4 | 238.7 | 239.5 | 238.8 | 239.3 | 0.8 | 0.3 |
| 33 | LPC(20:1)>LPC(20:1/0:0) _and_LPC(0:0/20:1) | C28 H56 N O7 P | 550.3867 | LPC | 2.42 | 2.42 | 2.42 | 2.43 | 239.9 | 239.3 | 240.3 | 241.4 | 240.2 | 0.9 | 0.4 |
| 34 | LPC(20:0)>LPC(20:0/0:0) _and_LPC(0:0/20:0) | C28 H58 N O7 P | 552.4025 | LPC | 2.72 | 2.72 | 2.73 | 2.74 | 241.8 | 240.8 | 241.3 | 242.6 | 241.6 | 0.8 | 0.3 |
| 35 | LPC(22:6)>LPC(22:6/0:0) _and_LPC(0:0/22:6) | C30 H50 N O7 P | 568.3398 | LPC | 1.77 | 1.77 | 1.76 | 1.77 | 244.7 | 243.1 | 243.9 | 242.6 | 243.6 | 0.9 | 0.4 |
| 36 | LPC(22:5)>LPC(22:5/0:0) _and_LPC(0:0/22:5) | C30 H52 N O7 P | 570.3554 | LPC | 1.90 | 1.90 | 1.94 | 1.95 | 244.7 | 243.6 | 244.9 | 244.3 | 244.4 | 0.6 | 0.2 |
| 37 | LPC(22:4)>LPC(22:4/0:0) _and_LPC(0:0/22:4) | C30 H54 N O7 P | 572.3711 | LPC | 2.13 | 2.13 | 2.15 | 2.16 | 245.5 | 245.7 | 246.5 | 247.8 | 246.4 | 1.0 | 0.4 |
| 38 | LPC(22:0)>LPC(22:0/0:0) _and_LPC(0:0/22:0) | C30 H62 N O7 P | 580.4340 | LPC | 3.12 | 3.13 | 3.13 | 3.13 | 256.9 | 252.0 | 256.9 | 254.0 | 255.0 | 2.4 | 0.9 |
| 39 | PC(28:0) | C36 H72 N O8 P | 678.5070 | PC | 3.53 | 3.53 | 3.53 | 3.53 | 275.1 | 274.6 | 273.0 | 276.3 | 274.7 | 1.4 | 0.5 |
| 40 | PC(O-30:0) | C38 H78 N O7 P | 692.5586 | PC | 4.55 | 4.55 | 4.57 | 4.56 | 289.6 | 289.3 | 282.9 | 282.3 | 286.0 | 3.9 | 1.4 |
| 41 | PC(30:1) | C38 H74 N O8 P | 704.5226 | PC | 3.58 | 3.58 | 3.59 | 3.59 | 281.2 | 283.1 | 281.9 | 282.0 | 282.1 | 0.8 | 0.3 |
| 42 | PC(30:0) | C38 H76 N O8 P | 706.5380 | PC | 4.10 | 4.10 | 4.10 | 4.09 | 279.1 | 280.0 | 278.4 | 278.1 | 278.9 | 0.8 | 0.3 |
| 43 | PC(O-32:2) | C40 H78 N O7 P | 716.5583 | PC | 4.54 | 4.53 | 4.54 | 4.53 | 284.6 | 284.5 | 282.7 | 282.3 | 283.5 | 1.2 | 0.4 |
| 44 | PC(31:1) | C39 H76 N O8 P | 718.5383 | PC | 3.86 | 3.86 | 3.87 | 3.88 | 292.1 | 291.8 | 286.4 | 285.3 | 288.9 | 3.5 | 1.2 |
| 45 | PC(O-32:1) | C40 H80 N O7 P | 718.5748 | PC | 5.26 | 5.25 | 5.27 | 5.27 | 283.4 | 283.5 | 281.4 | 281.8 | 282.5 | 1.1 | 0.4 |
| 46 | PC(O-32:1) | C40 H80 N O7 P | 718.5751 | PC | 4.66 | 4.66 | 4.68 | 4.66 | 285.8 | 285.1 | 280.7 | 282.3 | 283.5 | 2.4 | 0.8 |
| 47 | PC(O-32:0) | C40 H82 N O7 P | 720.5905 | PC | 5.38 | 5.37 | 5.40 | 5.38 | 283.8 | 284.3 | 281.6 | 281.7 | 282.9 | 1.4 | 0.5 |
| 48 | PC(32:2) | C40 H76 N O8 P | 730.5384 | PC | 3.70 | 3.69 | 3.70 | 3.70 | 284.0 | 285.4 | 283.9 | 283.4 | 284.2 | 0.9 | 0.3 |
| 49 | PC(32:1) | C40 H78 N O8 P | 732.5540 | PC | 4.18 | 4.18 | 4.18 | 4.17 | 284.4 | 285.3 | 283.7 | 283.4 | 284.2 | 0.9 | 0.3 |
| 50 | PC(32:0) | C40 H80 N O8 P | 734.5698 | PC | 4.83 | 4.82 | 4.82 | 4.82 | 283.4 | 283.8 | 282.9 | 283.0 | 283.3 | 0.4 | 0.1 |
| 51 | PC(33:2) | C41 H78 N O8 P | 744.5543 | PC | 4.00 | 4.00 | 4.01 | 4.01 | 288.8 | 290.0 | 288.4 | 288.1 | 288.8 | 0.8 | 0.3 |
| 52 | PC(O-34:2) | C42 H82 N O7 P | 744.5898 | PC | 5.32 | 5.32 | 5.33 | 5.32 | 289.0 | 289.7 | 288.1 | 288.3 | 288.8 | 0.7 | 0.3 |
| 53 | PC(O-34:2) | C42 H82 N O7 P | 744.5905 | PC | 4.83 | 4.82 | 4.82 | 4.82 | 292.2 | 291.4 | 285.4 | 285.0 | 288.5 | 3.8 | 1.3 |
| 54 | PC(33:1) | C41 H80 N O8 P | 746.5697 | PC | 4.52 | 4.52 | 4.53 | 4.53 | 285.9 | 287.6 | 286.8 | 287.2 | 286.9 | 0.7 | 0.3 |
| 55 | PC(O-34:1) | C42 H84 N O7 P | 746.6063 | PC | 5.44 | 5.43 | 5.45 | 5.46 | 289.9 | 290.8 | 288.2 | 287.9 | 289.2 | 1.4 | 0.5 |
| 56 | PC(O-34:0) | C42 H86 N O7 P | 748.6221 | PC | 6.31 | 6.31 | 6.31 | 6.30 | 291.5 | 293.0 | 289.4 | 288.7 | 290.6 | 1.9 | 0.7 |
| 57 | PC(34:4) | C42 H76 N O8 P | 754.5384 | PC | 3.62 | 3.61 | 3.62 | 3.62 | 292.8 | 292.5 | 290.7 | 290.0 | 291.5 | 1.4 | 0.5 |
| 58 | PC(34:3) | C42 H78 N O8 P | 756.5539 | PC | 3.77 | 3.76 | 3.77 | 3.78 | 292.6 | 292.5 | 291.0 | 290.2 | 291.6 | 1.2 | 0.4 |
| 59 | PC(34:3) | C42 H78 N O8 P | 756.5540 | PC | 3.89 | 3.89 | 3.90 | 3.90 | 292.6 | 292.4 | 291.2 | 290.2 | 291.6 | 1.1 | 0.4 |
| 60 | PC(34:2) | C42 H80 N O8 P | 758.5693 | PC | 4.32 | 4.32 | 4.32 | 4.33 | 290.2 | 290.0 | 290.4 | 290.7 | 290.3 | 0.3 | 0.1 |
| 61 | PC(34:1) | C42 H82 N O8 P | 760.5850 | PC | 4.89 | 4.89 | 4.88 | 4.88 | 293.1 | 293.1 | 293.4 | 292.9 | 293.1 | 0.2 | 0.1 |
| 62 | PC(34:0) | C42 H84 N O8 P | 762.6010 | PC | 5.70 | 5.70 | 5.69 | 5.69 | 297.3 | 297.8 | 293.5 | 290.6 | 294.8 | 3.4 | 1.2 |
| 63 | PC(O-36:5) | C44 H80 N O7 P | 766.5748 | PC | 4.23 | 4.23 | 4.23 | 4.21 | 320.3 | 300.3 | 307.2 | 307.0 | 308.7 | 8.4 | 2.7 |
| 64 | PC(35:4) | C43 H78 N O8 P | 768.5538 | PC | 5.19 | 5.19 | 5.20 | 5.20 | 296.5 | 297.6 | 294.4 | 293.4 | 295.5 | 1.9 | 0.7 |
| 65 | PC(35:4) | C43 H78 N O8 P | 768.5540 | PC | 3.89 | 3.89 | 3.90 | 3.90 | 296.7 | 296.9 | 294.1 | 293.0 | 295.2 | 1.9 | 0.6 |

| # | Name | Formula | *m/z* | Type | RT1 | RT2 | RT3 | RT4 | CCS1 | CCS2 | CCS3 | CCS4 | meanCCS | stdCCS | RSD(%) |
|---|---|---|---|---|---|---|---|---|---|---|---|---|---|---|---|
| 66 | PC(O-36:4) | C44 H82 N O7 P | 768.5903 | PC | 4.69 | 4.69 | 4.70 | 4.70 | 294.1 | 294.8 | 292.6 | 290.7 | 293.0 | 1.8 | 0.6 |
| 67 | PC(35:3) | C43 H80 N O8 P | 770.5691 | PC | 4.05 | 4.05 | 4.05 | 4.04 | 298.9 | 300.5 | 295.3 | 294.7 | 297.3 | 2.8 | 0.9 |
| 68 | PC(35:3) | C43 H80 N O8 P | 770.5695 | PC | 4.17 | 4.16 | 4.16 | 4.15 | 299.0 | 300.5 | 294.7 | 294.7 | 297.2 | 3.0 | 1.0 |
| 69 | PC(O-36:3) | C44 H84 N O7 P | 770.6058 | PC | 5.03 | 5.02 | 5.03 | 5.03 | 296.6 | 296.5 | 294.2 | 292.4 | 294.9 | 2.0 | 0.7 |
| 70 | PC(35:2) | C43 H82 N O8 P | 772.5850 | PC | 4.57 | 4.57 | 4.58 | 4.58 | 293.2 | 293.2 | 292.5 | 292.5 | 292.9 | 0.4 | 0.1 |
| 71 | PC(O-36:2) | C44 H86 N O7 P | 772.6216 | PC | 5.72 | 5.71 | 5.71 | 5.70 | 299.5 | 300.0 | 296.3 | 291.4 | 296.8 | 3.9 | 1.3 |
| 72 | PC(O-36:2) | C44 H86 N O7 P | 772.6217 | PC | 6.22 | 6.22 | 6.22 | 6.22 | 300.0 | 299.8 | 295.5 | 290.9 | 296.6 | 4.3 | 1.4 |
| 73 | PC(35:1) | C43 H84 N O8 P | 774.6007 | PC | 5.19 | 5.19 | 5.18 | 5.19 | 303.7 | 305.9 | 293.1 | 293.0 | 298.9 | 6.8 | 2.3 |
| 74 | PC(35:1) | C43 H84 N O8 P | 774.6008 | PC | 5.32 | 5.32 | 5.33 | 5.32 | 304.4 | 306.3 | 293.4 | 292.8 | 299.3 | 7.1 | 2.4 |
| 75 | PC(O-36:1) | C44 H88 N O7 P | 774.6370 | PC | 6.37 | 6.37 | 6.36 | 6.36 | 300.9 | 301.8 | 293.3 | 283.0 | 294.8 | 8.7 | 3.0 |
| 76 | PC(36:6) | C44 H76 N O8 P | 778.5382 | PC | 3.53 | 3.52 | 3.53 | 3.50 | 297.2 | 297.6 | 295.0 | 293.9 | 295.9 | 1.8 | 0.6 |
| 77 | PC(36:6) | C44 H76 N O8 P | 778.5384 | PC | 3.36 | 3.36 | 3.36 | 3.36 | 297.3 | 297.5 | 294.6 | 293.7 | 295.8 | 1.9 | 0.6 |
| 78 | PC(36:5) | C44 H78 N O8 P | 780.5538 | PC | 3.68 | 3.68 | 3.69 | 3.68 | 297.1 | 298.1 | 295.5 | 294.3 | 296.2 | 1.7 | 0.6 |
| 79 | PC(36:5) | C44 H78 N O8 P | 780.5539 | PC | 3.82 | 3.82 | 3.83 | 3.83 | 297.0 | 297.9 | 295.7 | 294.4 | 296.2 | 1.5 | 0.5 |
| 80 | PC(O-37:5) | C45 H82 N O7 P | 780.5900 | PC | 6.48 | 6.47 | 6.47 | 6.47 | 298.2 | 299.3 | 301.6 | 298.6 | 299.4 | 1.5 | 0.5 |
| 81 | PC(O-37:5) | C45 H82 N O7 P | 780.5903 | PC | 6.21 | 6.21 | 6.22 | 6.22 | 298.5 | 299.4 | 301.8 | 299.0 | 299.7 | 1.4 | 0.5 |
| 82 | PC(36:4) | C44 H80 N O8 P | 782.5693 | PC | 3.88 | 3.88 | 3.89 | 3.88 | 297.1 | 298.1 | 295.8 | 294.5 | 296.4 | 1.6 | 0.5 |
| 83 | PC(36:4) | C44 H80 N O8 P | 782.5694 | PC | 4.21 | 4.22 | 4.22 | 4.21 | 296.8 | 297.3 | 295.1 | 294.4 | 295.9 | 1.4 | 0.5 |
| 84 | PC(36:3) | C44 H82 N O8 P | 784.5848 | PC | 4.38 | 4.39 | 4.39 | 4.38 | 293.1 | 294.3 | 294.8 | 294.4 | 294.2 | 0.7 | 0.2 |
| 85 | PC(36:3) | C44 H82 N O8 P | 784.5851 | PC | 4.51 | 4.51 | 4.52 | 4.54 | 294.6 | 295.8 | 294.4 | 293.6 | 294.6 | 0.9 | 0.3 |
| 86 | PC(36:2) | C44 H84 N O8 P | 786.6007 | PC | 5.12 | 5.11 | 5.12 | 5.11 | 295.8 | 297.1 | 295.5 | 294.2 | 295.6 | 1.2 | 0.4 |
| 87 | PC(36:2) | C44 H84 N O8 P | 786.6007 | PC | 4.96 | 4.96 | 4.95 | 4.96 | 295.0 | 296.0 | 295.2 | 294.2 | 295.1 | 0.7 | 0.2 |
| 88 | PC(36:1) | C44 H86 N O8 P | 788.6165 | PC | 5.77 | 5.77 | 5.77 | 5.77 | 298.6 | 300.1 | 297.1 | 294.3 | 297.5 | 2.5 | 0.8 |
| 89 | PC(37:6) | C45 H78 N O8 P | 792.5536 | PC | 3.76 | 3.76 | 3.76 | 3.75 | 300.9 | 301.9 | 302.0 | 298.3 | 300.8 | 1.7 | 0.6 |
| 90 | PC(O-38:5) | C46 H84 N O7 P | 794.6058 | PC | 4.72 | 4.73 | 4.73 | 4.73 | 300.5 | 301.8 | 297.6 | 296.6 | 299.1 | 2.4 | 0.8 |
| 91 | PC(O-38:5) | C46 H84 N O7 P | 794.6059 | PC | 5.39 | 5.40 | 5.40 | 5.41 | 301.6 | 303.2 | 299.2 | 297.7 | 300.4 | 2.5 | 0.8 |
| 92 | PC(O-38:4) | C46 H86 N O7 P | 796.6217 | PC | 5.57 | 5.57 | 5.57 | 5.57 | 304.1 | 306.7 | 299.4 | 297.8 | 302.0 | 4.1 | 1.4 |
| 93 | PC(37:3) | C45 H84 N O8 P | 798.6002 | PC | 4.90 | 4.90 | 4.89 | 4.91 | 303.8 | 303.5 | 300.8 | 297.0 | 301.3 | 3.1 | 1.0 |
| 94 | PC(O-38:3) | C46 H88 N O7 P | 798.6372 | PC | 5.92 | 5.92 | 5.94 | 5.93 | 319.1 | 319.9 | 299.8 | 297.0 | 308.9 | 12.3 | 4.0 |
| 95 | PC(37:1) | C45 H88 N O8 P | 802.6322 | PC | 6.20 | 6.21 | 6.22 | 6.22 | 307.1 | 306.4 | 305.3 | 303.1 | 305.5 | 1.8 | 0.6 |
| 96 | PC(38:7) | C46 H78 N O8 P | 804.5538 | PC | 3.55 | 3.55 | 3.56 | 3.56 | 304.8 | 305.6 | 301.6 | 300.6 | 303.1 | 2.4 | 0.8 |
| 97 | PC(38:7) | C46 H78 N O8 P | 804.5539 | PC | 3.45 | 3.45 | 3.46 | 3.46 | 304.9 | 305.5 | 301.3 | 300.5 | 303.1 | 2.5 | 0.8 |
| 98 | PC(38:6) | C46 H80 N O8 P | 806.5694 | PC | 4.06 | 4.06 | 4.06 | 4.05 | 304.4 | 305.7 | 302.1 | 301.0 | 303.3 | 2.1 | 0.7 |
| 99 | PC(38:6) | C46 H80 N O8 P | 806.5695 | PC | 3.79 | 3.78 | 3.80 | 3.79 | 304.3 | 305.6 | 302.0 | 301.0 | 303.2 | 2.1 | 0.7 |
| 100 | PC(38:4) | C46 H84 N O8 P | 810.6006 | PC | 4.56 | 4.57 | 4.57 | 4.58 | 302.5 | 304.1 | 300.6 | 300.4 | 301.9 | 1.7 | 0.6 |
| 101 | PC(38:4) | C46 H84 N O8 P | 810.6008 | PC | 5.00 | 5.00 | 4.99 | 4.98 | 302.4 | 303.7 | 301.3 | 300.6 | 302.0 | 1.4 | 0.5 |
| 102 | PC(38:4) | C46 H84 N O8 P | 810.6008 | PC | 4.72 | 4.73 | 4.73 | 4.73 | 303.0 | 304.7 | 300.6 | 300.0 | 302.1 | 2.2 | 0.7 |
| 103 | PC(38:3) | C46 H86 N O8 P | 812.6165 | PC | 5.33 | 5.33 | 5.34 | 5.35 | 304.8 | 307.1 | 303.3 | 301.6 | 304.2 | 2.3 | 0.8 |
| 104 | PC(38:2) | C46 H88 N O8 P | 814.6321 | PC | 5.89 | 5.89 | 5.90 | 5.89 | 307.2 | 307.9 | 304.9 | 302.8 | 305.7 | 2.3 | 0.8 |
| 105 | PC(38:1) | C46 H90 N O8 P | 816.6475 | PC | 6.51 | 6.51 | 6.52 | 6.51 | 307.2 | 307.7 | 306.8 | 306.9 | 307.2 | 0.4 | 0.1 |
| 106 | PC(O-40:7) | C48 H84 N O7 P | 818.6052 | PC | 4.52 | 4.52 | 4.54 | 4.53 | 311.3 | 312.0 | 307.3 | 304.7 | 308.8 | 3.4 | 1.1 |
| 107 | PC(O-40:7) | C48 H84 N O7 P | 818.6057 | PC | 5.17 | 5.18 | 5.18 | 5.19 | 312.1 | 313.2 | 308.5 | 304.9 | 309.7 | 3.8 | 1.2 |
| 108 | PC(39:6) | C47 H82 N O8 P | 820.5850 | PC | 4.41 | 4.41 | 4.42 | 4.42 | 307.5 | 308.1 | 309.5 | 307.7 | 308.2 | 0.9 | 0.3 |
| 109 | PC(O-40:6) | C48 H86 N O7 P | 820.6210 | PC | 5.32 | 5.33 | 5.34 | 5.35 | 314.2 | 315.0 | 309.2 | 305.1 | 310.9 | 4.6 | 1.5 |
| 110 | PC(39:4) | C47 H86 N O8 P | 824.6166 | PC | 5.43 | 5.44 | 5.45 | 5.44 | 310.3 | 310.1 | 310.2 | 307.4 | 309.5 | 1.4 | 0.5 |
| 111 | PC(O-40:4) | C48 H90 N O7 P | 824.6524 | PC | 6.45 | 6.45 | 6.45 | 6.44 | 321.0 | 321.5 | 311.4 | 305.3 | 314.8 | 7.8 | 2.5 |
| 112 | PC(40:8) | C48 H80 N O8 P | 830.5695 | PC | 3.67 | 3.67 | 3.68 | 3.67 | 312.2 | 312.5 | 309.1 | 307.1 | 310.2 | 2.6 | 0.8 |
| 113 | PC(40:7) | C48 H82 N O8 P | 832.5845 | PC | 4.19 | 4.19 | 4.19 | 4.20 | 312.5 | 312.6 | 308.9 | 307.5 | 310.3 | 2.6 | 0.8 |

| # | Name | Formula | *m/z* | Type | RT1 | RT2 | RT3 | RT4 | CCS1 | CCS2 | CCS3 | CCS4 | meanCCS | stdCCS | RSD(%) |
|---|---|---|---|---|---|---|---|---|---|---|---|---|---|---|---|
| 114 | PC(40:7) | C48 H82 N O8 P | 832.5849 | PC | 4.12 | 4.11 | 4.11 | 4.10 | 312.5 | 312.8 | 309.5 | 307.3 | 310.5 | 2.6 | 0.9 |
| 115 | PC(40:7) | C48 H82 N O8 P | 832.5849 | PC | 3.93 | 3.93 | 3.92 | 3.94 | 312.5 | 313.0 | 309.4 | 307.4 | 310.6 | 2.6 | 0.8 |
| 116 | PC(40:6) | C48 H84 N O8 P | 834.6007 | PC | 4.80 | 4.79 | 4.80 | 4.79 | 311.6 | 312.0 | 307.4 | 306.4 | 309.3 | 2.8 | 0.9 |
| 117 | PC(42:7) | C50 H86 N O8 P | 838.6320 | PC | 5.60 | 5.59 | 5.59 | 5.58 | 313.9 | 314.8 | 309.9 | 308.3 | 311.7 | 3.1 | 1.0 |
| 118 | PC(42:4) | C50 H92 N O8 P | 848.6524 | PC | 6.24 | 6.24 | 6.25 | 6.24 | 319.3 | 319.4 | 316.6 | 313.1 | 317.1 | 2.9 | 0.9 |
| 119 | PC(O-42:6) | C50 H90 N O7 P | 848.6525 | PC | 5.67 | 5.66 | 5.66 | 5.65 | 318.7 | 319.9 | 317.5 | 313.8 | 317.5 | 2.6 | 0.8 |
| 120 | PC(42:8) | C50 H84 N O8 P | 858.6008 | PC | 4.12 | 4.12 | 4.11 | 4.10 | 319.7 | 322.0 | 319.8 | 320.2 | 320.4 | 1.1 | 0.3 |
| 121 | PC(42:5) | C50 H90 N O8 P | 864.6475 | PC | 5.83 | 5.83 | 5.83 | 5.83 | 323.7 | 325.6 | 322.8 | 318.3 | 322.6 | 3.1 | 1.0 |
| 122 | PC(42:4) | C50 H92 N O8 P | 866.6639 | PC | 6.63 | 6.63 | 6.63 | 6.63 | 323.5 | 325.0 | 323.1 | 320.3 | 323.0 | 2.0 | 0.6 |
| 123 | PE(O-36:3)>PE(O-18:1/18:2) | C41 H78 N O7 P | 728.5593 | PE | 5.60 | 5.60 | 5.60 | 5.59 | 288.4 | 288.0 | 288.0 | 286.5 | 287.7 | 0.8 | 0.3 |
| 124 | PE(O-38:6)>PE(O-18:2/20:4)_and_PE(O-16:1/22:5) | C43 H76 N O7 P | 750.5431 | PE | 4.68 | 4.68 | 4.69 | 4.68 | 293.6 | 294.9 | 293.7 | 290.2 | 293.1 | 2.0 | 0.7 |
| 125 | PE(O-40:4) | C45 H84 N O7 P | 782.6045 | PE | 5.11 | 5.09 | 5.11 | 5.11 | 295.7 | 295.4 | 308.9 | 299.9 | 300.0 | 6.3 | 2.1 |
| 126 | SM(d32:2) | C37 H73 N2 O6 P | 673.5281 | SM | 3.09 | 3.08 | 3.10 | 3.10 | 272.0 | 271.6 | 270.9 | 271.8 | 271.6 | 0.5 | 0.2 |
| 127 | SM(d32:1) | C37 H75 N2 O6 P | 675.5438 | SM | 3.49 | 3.48 | 3.49 | 3.50 | 272.1 | 271.5 | 271.0 | 272.3 | 271.7 | 0.6 | 0.2 |
| 128 | SM(d32:0) | C37 H77 N2 O6 P | 677.5598 | SM | 3.69 | 3.69 | 3.69 | 3.70 | 279.4 | 278.2 | 273.4 | 274.2 | 276.3 | 2.9 | 1.1 |
| 129 | SM(d33:1) | C38 H77 N2 O6 P | 689.5595 | SM | 3.76 | 3.76 | 3.76 | 3.76 | 275.5 | 277.3 | 275.2 | 276.0 | 276.0 | 0.9 | 0.3 |
| 130 | SM(d34:2) | C39 H77 N2 O6 P | 701.5595 | SM | 3.57 | 3.57 | 3.58 | 3.58 | 277.2 | 278.1 | 276.8 | 276.9 | 277.2 | 0.6 | 0.2 |
| 131 | SM(d34:1) | C39 H79 N2 O6 P | 703.5749 | SM | 4.06 | 4.07 | 4.06 | 4.06 | 276.8 | 277.6 | 276.7 | 276.6 | 276.9 | 0.4 | 0.2 |
| 132 | SM(d34:0) | C39 H81 N2 O6 P | 705.5905 | SM | 4.31 | 4.32 | 4.32 | 4.33 | 281.2 | 281.2 | 279.7 | 279.5 | 280.4 | 1.0 | 0.3 |
| 133 | SM(d35:2) | C40 H79 N2 O6 P | 715.5749 | SM | 3.86 | 3.86 | 3.84 | 3.83 | 288.6 | 286.7 | 285.3 | 283.5 | 286.0 | 2.1 | 0.8 |
| 134 | SM(d35:1) | C40 H81 N2 O6 P | 717.5906 | SM | 4.43 | 4.43 | 4.44 | 4.43 | 282.5 | 283.3 | 282.2 | 283.5 | 282.9 | 0.6 | 0.2 |
| 135 | SM(d35:1) | C40 H81 N2 O6 P | 717.5908 | SM | 4.26 | 4.27 | 4.27 | 4.28 | 297.4 | 298.6 | 287.4 | 291.4 | 293.7 | 5.3 | 1.8 |
| 136 | SM(d34:0-OH) | C39 H81 N2 O7 P | 721.5858 | SM | 3.74 | 3.74 | 3.76 | 3.75 | 285.9 | 287.1 | 286.3 | 286.3 | 286.4 | 0.5 | 0.2 |
| 137 | SM(d36:3) | C41 H79 N2 O6 P | 727.5753 | SM | 3.72 | 3.72 | 3.72 | 3.72 | 284.3 | 285.3 | 286.0 | 285.7 | 285.3 | 0.7 | 0.3 |
| 138 | SM(d36:2) | C41 H81 N2 O6 P | 729.5906 | SM | 4.19 | 4.19 | 4.19 | 4.19 | 285.1 | 285.7 | 285.3 | 285.4 | 285.4 | 0.2 | 0.1 |
| 139 | SM(d36:1) | C41 H83 N2 O6 P | 731.6065 | SM | 4.81 | 4.81 | 4.81 | 4.81 | 284.0 | 284.2 | 284.3 | 284.1 | 284.1 | 0.1 | 0.0 |
| 140 | SM(d36:0) | C41 H85 N2 O6 P | 733.6220 | SM | 5.12 | 5.11 | 5.12 | 5.11 | 294.4 | 293.3 | 287.8 | 289.9 | 291.4 | 3.0 | 1.0 |
| 141 | SM(d37:1) | C42 H85 N2 O6 P | 745.6220 | SM | 5.27 | 5.28 | 5.30 | 5.29 | 291.7 | 293.9 | 286.7 | 290.8 | 290.8 | 3.0 | 1.0 |
| 142 | SM(d38:1) | C43 H87 N2 O6 P | 759.6376 | SM | 5.77 | 5.77 | 5.77 | 5.77 | 291.9 | 292.9 | 293.9 | 292.3 | 292.7 | 0.9 | 0.3 |
| 143 | SM(d39:1) | C44 H89 N2 O6 P | 773.6532 | SM | 6.22 | 6.21 | 6.23 | 6.23 | 294.9 | 296.5 | 295.7 | 294.7 | 295.4 | 0.8 | 0.3 |
| 144 | SM(d40:2) | C45 H89 N2 O6 P | 785.6533 | SM | 5.69 | 5.69 | 5.68 | 5.68 | 300.8 | 301.7 | 300.3 | 298.8 | 300.4 | 1.2 | 0.4 |
| 145 | SM(d40:2) | C45 H89 N2 O6 P | 785.6534 | SM | 5.90 | 5.90 | 5.91 | 5.91 | 301.0 | 301.6 | 299.9 | 298.5 | 300.3 | 1.4 | 0.5 |
| 146 | SM(d40:1) | C45 H91 N2 O6 P | 787.6688 | SM | 6.53 | 6.53 | 6.53 | 6.54 | 302.7 | 302.4 | 299.5 | 298.5 | 300.8 | 2.1 | 0.7 |
| 147 | SM(d41:2) | C46 H91 N2 O6 P | 799.6690 | SM | 6.36 | 6.35 | 6.35 | 6.34 | 303.9 | 303.6 | 302.0 | 301.1 | 302.6 | 1.3 | 0.4 |
| 148 | SM(d41:2) | C46 H91 N2 O6 P | 799.6690 | SM | 6.09 | 6.10 | 6.11 | 6.10 | 303.9 | 303.7 | 302.2 | 301.4 | 302.8 | 1.2 | 0.4 |
| 149 | SM(d41:1) | C46 H93 N2 O6 P | 801.6847 | SM | 6.75 | 6.75 | 6.75 | 6.75 | 303.9 | 304.1 | 301.5 | 300.4 | 302.5 | 1.8 | 0.6 |
| 150 | SM(d42:3) | C47 H91 N2 O6 P | 811.6686 | SM | 5.79 | 5.80 | 5.80 | 5.80 | 307.4 | 308.7 | 306.9 | 304.6 | 306.9 | 1.7 | 0.6 |
| 151 | SM(d42:2) | C47 H93 N2 O6 P | 813.6845 | SM | 6.47 | 6.47 | 6.46 | 6.46 | 308.6 | 309.2 | 306.8 | 304.5 | 307.3 | 2.1 | 0.7 |
| 152 | SM(d43:3) | C48 H93 N2 O6 P | 825.6845 | SM | 6.23 | 6.24 | 6.24 | 6.24 | 310.7 | 311.9 | 309.7 | 308.3 | 310.2 | 1.5 | 0.5 |
| 153 | SM(d43:2) | C48 H95 N2 O6 P | 827.7000 | SM | 6.69 | 6.70 | 6.70 | 6.70 | 311.2 | 313.8 | 310.2 | 308.1 | 310.8 | 2.4 | 0.8 |
| 154 | SM(d43:2) | C48 H95 N2 O6 P | 827.7001 | SM | 6.82 | 6.81 | 6.82 | 6.81 | 311.3 | 313.7 | 309.9 | 307.8 | 310.7 | 2.5 | 0.8 |
| 155 | SM(t42:1)_or_SM(d42:1-OH) | C47 H95 N2 O7 P | 831.6951 | SM | 6.07 | 6.07 | 6.08 | 6.08 | 314.2 | 315.0 | 311.6 | 310.8 | 312.9 | 2.0 | 0.6 |
| 156 | SM(d44:3) | C49 H95 N2 O6 P | 839.6997 | SM | 6.56 | 6.56 | 6.57 | 6.56 | 314.7 | 315.6 | 314.0 | 312.4 | 314.2 | 1.3 | 0.4 |

| # | Name | Formula | *m/z* | Type | RT1 | RT2 | RT3 | RT4 | CCS1 | CCS2 | CCS3 | CCS4 | meanCCS | stdCCS | RSD(%) |
|---|---|---|---|---|---|---|---|---|---|---|---|---|---|---|---|
| 157 | TG(44:1)>TG(10:0_16:0_18:1) + NH3 | C47 H91 N O6 | 766.6921 | TG | 7.76 | 7.75 | 7.76 | 7.76 | 295.2 | 296.3 | 298.1 | 296.8 | 296.6 | 1.2 | 0.4 |
| 158 | TG(46:2) + NH3 | C49 H93 N O6 | 792.7076 | TG | 7.79 | 7.79 | 7.79 | 7.79 | 301.6 | 302.3 | 304.2 | 302.8 | 302.7 | 1.1 | 0.4 |
| 159 | TG(46:1)>TG(12:0_16:0_18:1)_and_TG(10:0_18:0_18:1)_and_TG(14:0_16:0_16:1) + NH3 | C49 H95 N O6 | 794.7235 | TG | 8.07 | 8.07 | 8.07 | 8.07 | 300.7 | 302.1 | 304.4 | 305.2 | 303.1 | 2.1 | 0.7 |
| 160 | TG(46:0)>TG(14:0_16:0_16:0)_and_TG(12:0_16:0_18:0) + NH3 | C49 H97 N O6 | 796.7390 | TG | 8.44 | 8.43 | 8.44 | 8.44 | 309.3 | 309.5 | 305.9 | 307.9 | 308.1 | 1.7 | 0.5 |
| 161 | TG(48:3) + NH3 | C51 H95 N O6 | 818.7235 | TG | 7.80 | 7.80 | 7.80 | 7.81 | 308.3 | 309.7 | 308.6 | 307.4 | 308.5 | 0.9 | 0.3 |
| 162 | TG(48:2)>TG(14:0_16:0_18:2)_and_TG(14:0_16:1_18:1) + NH3 | C51 H97 N O6 | 820.7390 | TG | 8.11 | 8.11 | 8.11 | 8.10 | 308.6 | 309.3 | 309.0 | 309.6 | 309.1 | 0.4 | 0.1 |
| 163 | TG(48:1)>TG(16:0_16:0_16:1)_and_TG(14:0_16:0_18:1) + NH3 | C51 H99 N O6 | 822.7546 | TG | 8.42 | 8.42 | 8.43 | 8.42 | 308.9 | 309.4 | 308.8 | 309.9 | 309.2 | 0.5 | 0.2 |
| 164 | TG(49:3) + NH3 | C52 H97 N O6 | 832.7390 | TG | 7.95 | 7.95 | 7.96 | 7.96 | 312.3 | 315.1 | 311.5 | 308.1 | 311.8 | 2.9 | 0.9 |
| 165 | TG(49:1)>TG(15:0_16:0_18:0)_and_TG(16:0_16:0_17:1)_and_TG(16:0_16:1_17:0) + NH3 | C52 H101 N O6 | 836.7702 | TG | 8.61 | 8.61 | 8.62 | 8.61 | 316.2 | 319.5 | 316.1 | 311.3 | 315.8 | 3.4 | 1.1 |
| 166 | TG(50:4)>TG(16:1_16:1_18:2)_and_TG(16:1_16:1_18:2) + NH3 | C53 H97 N O6 | 844.7388 | TG | 7.86 | 7.84 | 7.85 | 7.86 | 314.2 | 315.4 | 315.0 | 312.4 | 314.2 | 1.3 | 0.4 |
| 167 | TG(50:3) + NH3 | C53 H99 N O6 | 846.7545 | TG | 8.13 | 8.13 | 8.12 | 8.12 | 314.2 | 316.2 | 314.8 | 313.4 | 314.7 | 1.2 | 0.4 |
| 168 | TG(50:2)>TG(16:0_16:1_18:1)_and_TG(14:0_18:1_18:1) + NH3 | C53 H101 N O6 | 848.7701 | TG | 8.42 | 8.42 | 8.42 | 8.42 | 314.2 | 317.4 | 315.3 | 314.7 | 315.4 | 1.4 | 0.4 |
| 169 | TG(50:1)>TG(16:0_16:0_18:1) + NH3 | C53 H103 N O6 | 850.7858 | TG | 8.81 | 8.81 | 8.81 | 8.81 | 315.2 | 316.8 | 316.8 | 316.8 | 316.4 | 0.8 | 0.2 |
| 170 | TG(51:4)>TG(16:1_17:1_18:2)_and_TG(15:0_18:2_18:2)_and_TG(15:1_18:1_18:2) + NH3 | C54 H99 N O6 | 858.7545 | TG | 8.00 | 8.01 | 8.01 | 8.01 | 317.7 | 319.8 | 320.0 | 317.7 | 318.8 | 1.3 | 0.4 |
| 171 | TG(51:2) M + NH3 | C54 H103 N O6 | 862.7854 | TG | 7.79 | 7.79 | 7.79 | 7.79 | 318.9 | 320.0 | 320.9 | 319.6 | 319.9 | 0.8 | 0.3 |
| 172 | TG(51:2)>TG(15:0_18:1_18:1)_and_TG(16:0_17:1_18:1)_and_TG(16:1_17:0_18:1) + NH3 | C54 H103 N O6 | 862.7858 | TG | 8.60 | 8.60 | 8.60 | 8.60 | 318.8 | 320.2 | 320.2 | 320.2 | 319.8 | 0.7 | 0.2 |
| 173 | TG(52:6) M + NH3 | C55 H97 N O6 | 868.7390 | TG | 7.65 | 7.65 | 7.74 | 7.74 | 319.6 | 319.4 | 320.2 | 316.7 | 319.0 | 1.6 | 0.5 |
| 174 | TG(52:5)>TG(16:0_18:2_18:3) + NH3 | C55 H99 N O6 | 870.7545 | TG | 7.93 | 7.94 | 7.94 | 7.94 | 319.6 | 320.4 | 320.2 | 317.4 | 319.4 | 1.4 | 0.4 |
| 175 | TG(52:4)>TG(16:0_16:0_20:4)_and_TG(16:1_18:1_18:2)_and_TG(16:0_18:1_18:2) + NH3 | C55 H101 N O6 | 872.7702 | TG | 8.18 | 8.18 | 8.17 | 8.18 | 319.8 | 320.8 | 320.1 | 317.9 | 319.7 | 1.2 | 0.4 |
| 176 | TG(52:3)>TG(16:0_18:1_18:2)_and_TG(16:1_18:1_18:1) + NH3 | C55 H103 N O6 | 874.7858 | TG | 8.48 | 8.47 | 8.48 | 8.48 | 320.6 | 321.5 | 320.5 | 318.3 | 320.2 | 1.3 | 0.4 |

| # | Name | Formula | *m/z* | Type | RT1 | RT2 | RT3 | RT4 | CCS1 | CCS2 | CCS3 | CCS4 | meanCCS | stdCCS | RSD(%) |
|---|---|---|---|---|---|---|---|---|---|---|---|---|---|---|---|
| 177 | TG(52:2)>TG(16:0_18:1_18:1) + NH3 | C55 H105 N O6 | 876.8014 | TG | 7.79 | 7.80 | 7.80 | 7.80 | 337.4 | 337.1 | 336.8 | 320.4 | 333.0 | 8.3 | 2.5 |
| 178 | TG(52:2)>TG(16:0_18:1_18:1) + NH3 | C55 H105 N O6 | 876.8015 | TG | 8.80 | 8.79 | 8.80 | 8.79 | 320.5 | 322.4 | 321.0 | 319.1 | 320.8 | 1.4 | 0.4 |
| 179 | TG(53:4)>TG(17:0_18:2_18:2)_and_TG(17:1_18:1_18:2) + NH3 | C56 H103 N O6 | 886.7856 | TG | 8.30 | 8.30 | 8.30 | 8.30 | 327.2 | 329.7 | 328.8 | 324.6 | 327.6 | 2.2 | 0.7 |
| 180 | TG(53:3)>TG(17:0_18:1_18:2)_and_TG(17:1_18:1_18:1)_and_TG(16:1_18:1_19:1)_and_TG(16:0_18:1_19:2) + NH3 | C56 H105 N O6 | 888.8013 | TG | 7.80 | 7.80 | 7.81 | 7.81 | 328.0 | 330.1 | 329.6 | 324.3 | 328.0 | 2.7 | 0.8 |
| 181 | TG(53:3)>TG(17:0_18:1_18:2)_and_TG(17:1_18:1_18:1)_and_TG(16:1_18:1_19:1)_and_TG(16:0_18:1_19:2) + NH3 | C56 H105 N O6 | 888.8014 | TG | 8.66 | 8.66 | 8.67 | 8.66 | 327.9 | 330.2 | 328.8 | 324.8 | 327.9 | 2.3 | 0.7 |
| 182 | TG(53:2) + NH3 | C56 H107 N O6 | 890.8169 | TG | 8.82 | 8.81 | 8.81 | 8.80 | 331.6 | 334.1 | 331.6 | 332.4 | 332.4 | 1.2 | 0.3 |
| 183 | TG(53:2) + NH3 | C56 H107 N O6 | 890.8170 | TG | 8.99 | 8.99 | 9.00 | 9.00 | 331.6 | 333.9 | 332.1 | 331.9 | 332.4 | 1.0 | 0.3 |
| 184 | TG(54:7)>TG(16:0_16:1_22:6)_and_TG(16:0_18:2_20:5) + NH3 | C57 H99 N O6 | 894.7543 | TG | 7.84 | 7.84 | 7.85 | 7.84 | 326.2 | 328.1 | 328.5 | 322.9 | 326.4 | 2.6 | 0.8 |
| 185 | TG(54:6)>TG(16:0_18:2_20:4) + NH3 | C57 H101 N O6 | 896.7701 | TG | 8.07 | 8.06 | 8.06 | 8.07 | 326.8 | 329.5 | 329.5 | 323.6 | 327.4 | 2.8 | 0.9 |
| 186 | TG(54:5)>TG(18:1_18:2_18:2)_and_TG(16:0_16:0_22:5)_and_TG(16:0_18:1_20:4) + NH3 | C57 H103 N O6 | 898.7857 | TG | 8.36 | 8.35 | 8.36 | 8.36 | 327.4 | 329.9 | 329.5 | 324.0 | 327.7 | 2.7 | 0.8 |
| 187 | TG(54:4)>TG(18:1_18:1_18:2) + NH3 | C57 H105 N O6 | 900.8012 | TG | 8.47 | 8.46 | 8.47 | 8.47 | 329.0 | 331.5 | 329.9 | 324.4 | 328.7 | 3.0 | 0.9 |
| 188 | TG(54:3)>TG(18:0_18:1_18:2)_and_TG(18:1/18:1/18:1)_and_TG(16:0_18:2_20:1) + NH3 | C57 H107 N O6 | 902.8169 | TG | 8.87 | 8.87 | 8.87 | 8.87 | 330.3 | 331.8 | 329.7 | 324.7 | 329.1 | 3.1 | 0.9 |
| 189 | TG(54:2)>TG(16:0_18:1_20:1)_and_TG(18:0_18:1_18:1) + NH3 | C57 H109 N O6 | 904.8326 | TG | 9.20 | 9.20 | 9.20 | 9.20 | 334.5 | 335.1 | 334.8 | 331.2 | 333.9 | 1.8 | 0.5 |
| 190 | TG(55:4)>TG(18:1_18:2_19:1)_and_TG(18:2_18:2_19:0)_and_TG(18:1_18:1_19:2) + NH3 | C58 H107 N O6 | 914.8167 | TG | 7.85 | 7.83 | 7.85 | 7.86 | 342.9 | 344.0 | 342.1 | 335.8 | 341.2 | 3.7 | 1.1 |
| 191 | TG(55:4)>TG(18:1_18:2_19:1)_and_TG(18:2_18:2_19:0)_and_TG(18:1_18:1_19:2) + NH3 | C58 H107 N O6 | 914.8171 | TG | 8.65 | 8.65 | 8.65 | 8.65 | 343.0 | 344.1 | 341.5 | 336.4 | 341.2 | 3.4 | 1.0 |
| 192 | TG(55:3)>TG(18:1_18:1_19:1)_and_TG(18:1_18:2_19:0) + NH3 | C58 H109 N O6 | 916.8328 | TG | 8.97 | 8.97 | 8.98 | 8.98 | 343.9 | 344.6 | 342.0 | 336.3 | 341.7 | 3.7 | 1.1 |
| 193 | TG(55:2)>TG(18:1_18:1_19:0) + NH3 | C58 H111 N O6 | 918.8513 | TG | 8.81 | 8.82 | 8.81 | 8.81 | 344.0 | 344.9 | 343.0 | 337.3 | 342.3 | 3.4 | 1.0 |

| # | Name | Formula | *m/z* | Type | RT1 | RT2 | RT3 | RT4 | CCS1 | CCS2 | CCS3 | CCS4 | meanCCS | stdCCS | RSD(%) |
|---|---|---|---|---|---|---|---|---|---|---|---|---|---|---|---|
| 194 | TG(56:6)>TG(18:1_18:1_20:4)_and_TG(16:0_18:1_22:5)_and_TG(18:1_18:2_20:3) + NH3 | C59 H105 N O6 | 924.8013 | TG | 8.35 | 8.36 | 8.36 | 8.36 | 339.8 | 341.2 | 342.8 | 332.3 | 339.0 | 4.6 | 1.4 |
| 195 | TG(56:4)>TG(18:1_18:2_20:1)_and_TG(18:1_18:1_20:2) + NH3 | C59 H109 N O6 | 928.8322 | TG | 8.85 | 8.84 | 8.84 | 8.84 | 347.0 | 348.4 | 348.8 | 337.9 | 345.5 | 5.1 | 1.5 |
| 196 | TG(57:2) + NH3 | C60 H115 N O6 | 946.8795 | TG | 8.79 | 8.79 | 8.80 | 8.80 | 345.3 | 345.6 | 347.3 | 344.5 | 345.7 | 1.2 | 0.3 |
| 197 | TG(58:8) + NH3 | C61 H105 N O6 | 948.8010 | TG | 8.22 | 8.22 | 8.21 | 8.20 | 345.6 | 347.2 | 347.1 | 340.8 | 345.2 | 3.0 | 0.9 |
| 198 | TG(57:1)>TG(16:0_18:1_23:0)_and_TG(18:0_18:1_21:0) + NH3 | C60 H117 N O6 | 948.8952 | TG | 9.22 | 9.22 | 9.22 | 9.22 | 350.4 | 350.6 | 354.0 | 347.6 | 350.7 | 2.6 | 0.8 |
| 199 | TG(58:4) + NH3 | C61 H113 N O6 | 956.8637 | TG | 8.18 | 8.18 | 8.18 | 8.18 | 344.4 | 345.7 | 347.2 | 346.0 | 345.9 | 1.2 | 0.3 |
| 200 | TG(58:3) + NH3 | C61 H115 N O6 | 958.8795 | TG | 8.49 | 8.47 | 8.49 | 8.48 | 345.4 | 346.3 | 347.6 | 347.2 | 346.6 | 1.0 | 0.3 |
| 201 | TG(58:2) + NH3 | C61 H117 N O6 | 960.8951 | TG | 8.79 | 8.79 | 8.80 | 8.79 | 346.0 | 346.2 | 347.9 | 347.2 | 346.8 | 0.9 | 0.3 |
| 202 | TG(58:1)>TG(16:0_18:1_24:0)_and_TG(16:0_16:0_26:1)_and_TG(18:0_18:1_22:0) + NH3 | C61 H119 N O6 | 962.9108 | TG | 9.22 | 9.23 | 9.22 | 9.23 | 351.5 | 351.6 | 352.8 | 351.4 | 351.8 | 0.7 | 0.2 |

**Supplementary Table S3. Summary of lipid features with matched $^{Orbi}$CCS and $^{TIMS}$CCS values from a single LC-Orbitrap experiment (pump speed 68%, resolution setting 240,000). The same dataset present in Fig. 3a-f.**

| # | Name | Type | m/z_this study | m/z_database | Error_ppm | RT (min) | Orbi_CCS | TIMS_CCS | % error |
|---|---|---|---|---|---|---|---|---|---|
| 1 | CE(20:4)M+NH3 | CE | 690.6186 | 690.6187 | 0.1 | 8.92 | 280.7 | 294 | -4.5 |
| 2 | CE(22:6)M+NH3 | CE | 714.6188 | 714.6185 | 0.7 | 8.57 | 287.3 | 295.9 | -2.9 |
| 3 | LPC(14:0)>LPC(14:0/0:0)_and_LPC(0:0/14:0) | LPC | 468.3085 | 468.3084 | 0.6 | 8.7 | 214.2 | 224.6 | -4.6 |
| 4 | LPC(14:0)>LPC(14:0/0:0)_and_LPC(0:0/14:0) | LPC | 468.3085 | 468.3087 | 0.1 | 1.67 | 214.5 | 225.5 | -4.9 |
| 5 | LPC(15:0)>LPC(15:0/0:0)_and_LPC(0:0/15:0) | LPC | 482.3241 | 482.3245 | 0.4 | 1.57 | 220.1 | 223.3 | -1.4 |
| 6 | LPC(15:0)>LPC(15:0/0:0)_and_LPC(0:0/15:0) | LPC | 482.3243 | 482.3243 | 0.4 | 1.82 | 220.7 | 228.3 | -3.3 |
| 7 | LPC(16:1)>LPC(16:1/0:0)_and_LPC(0:0/16:1) | LPC | 494.3242 | 494.3244 | 1.2 | 2.03 | 222.1 | 228.5 | -2.8 |
| 8 | LPC(16:1)>LPC(16:1/0:0)_and_LPC(0:0/16:1) | LPC | 494.3242 | 494.3243 | 0.5 | 2.77 | 221.7 | 227.1 | -2.4 |
| 9 | LPC(16:0)>LPC(16:0/0:0)_and_LPC(0:0/16:0) | LPC | 496.3399 | 496.3402 | 0.2 | 1.69 | 220.4 | 233.7 | -5.7 |
| 10 | LPC(16:0)>LPC(16:0/0:0)_and_LPC(0:0/16:0) | LPC | 496.3399 | 496.3405 | 0.3 | 1.78 | 220.7 | 232.1 | -4.9 |
| 11 | LPC(17:0)>LPC(17:0/0:0)_and_LPC(0:0/17:0) | LPC | 510.3556 | 510.3555 | 0.4 | 2.22 | 225.4 | 236.4 | -4.7 |
| 12 | LPC(18:3)>LPC(18:3/0:0)_and_LPC(0:0/18:3) | LPC | 518.3242 | 518.3242 | 0.1 | 2 | 230.0 | 226.1 | 1.7 |
| 13 | LPC(20:5)>LPC(20:5/0:0)_and_LPC(0:0/20:5) | LPC | 520.3397 | 520.3405 | 1.2 | 2.4 | 229.2 | 229.1 | 0.0 |
| 14 | LPC(18:2)>LPC(18:2/0:0)_and_LPC(0:0/18:2) | LPC | 520.3399 | 520.3402 | 0.4 | 2.77 | 229.6 | 231.2 | -0.7 |
| 15 | LPC(18:1)>LPC(18:1/0:0)_and_LPC(0:0/18:1) | LPC | 522.3552 | 522.3562 | 1.1 | 2.78 | 229.2 | 235.2 | -2.6 |
| 16 | LPC(18:1)>LPC(18:1/0:0)_and_LPC(0:0/18:1) | LPC | 522.3555 | 522.3554 | 0.2 | 2.17 | 229.4 | 236.9 | -3.2 |
| 17 | LPC(18:0)>LPC(18:0/0:0)_and_LPC(0:0/18:0) | LPC | 524.3710 | 524.3717 | 1.5 | 1.91 | 229.8 | 240.1 | -4.3 |
| 18 | LPC(18:0)>LPC(18:0/0:0)_and_LPC(0:0/18:0) | LPC | 524.3713 | 524.3715 | 0.7 | 1.84 | 229.9 | 241 | -4.6 |
| 19 | LPC(19:1)>LPC(19:1/0:0)_and_LPC(0:0/19:1) | LPC | 536.3712 | 536.3720 | 0.0 | 1.69 | 237.6 | 237.8 | -0.1 |
| 20 | LPC(19:0)>LPC(19:0/0:0)_and_LPC(0:0/19:0) | LPC | 538.3870 | 538.3872 | 1.1 | 1.65 | 239.8 | 243.2 | -1.4 |
| 21 | LPC(20:4)>LPC(20:4/0:0)_and_LPC(0:0/20:4) | LPC | 544.3398 | 544.3399 | 0.4 | 2.54 | 235.8 | 232.6 | 1.4 |
| 22 | LPC(20:4)>LPC(20:4/0:0)_and_LPC(0:0/20:4) | LPC | 544.3398 | 544.3397 | 1.2 | 2.31 | 236.3 | 233.8 | 1.1 |
| 23 | LPC(20:3)>LPC(20:3/0:0)_and_LPC(0:0/20:3) | LPC | 546.3553 | 546.3552 | 0.1 | 2.74 | 235.9 | 233.8 | 0.9 |
| 24 | LPC(20:3)>LPC(20:3/0:0)_and_LPC(0:0/20:3) | LPC | 546.3559 | 546.3547 | 0.1 | 2.43 | 236.1 | 235.4 | 0.3 |
| 25 | LPC(20:2)>LPC(20:2/0:0)_and_LPC(0:0/20:2) | LPC | 548.3712 | 548.3708 | 0.8 | 2.23 | 238.7 | 236.5 | 0.9 |
| 26 | LPC(20:1)>LPC(20:1/0:0)_and_LPC(0:0/20:1) | LPC | 550.3868 | 550.3868 | 0.5 | 2.04 | 239.3 | 242.4 | -1.3 |
| 27 | LPC(20:0)>LPC(20:0/0:0)_and_LPC(0:0/20:0) | LPC | 552.4027 | 552.4025 | 1.6 | 1.98 | 240.8 | 246.4 | -2.3 |
| 28 | LPC(22:6)>LPC(22:6/0:0)_and_LPC(0:0/22:6) | LPC | 568.3398 | 568.3397 | 0.0 | 1.82 | 243.1 | 236.4 | 2.8 |
| 29 | LPC(22:6)>LPC(22:6/0:0)_and_LPC(0:0/22:6) | LPC | 568.3399 | 568.3396 | 0.1 | 1.89 | 243.3 | 235.2 | 3.5 |
| 30 | LPC(22:5)>LPC(22:5/0:0)_and_LPC(0:0/22:5) | LPC | 570.3555 | 570.3553 | 1.1 | 3.13 | 243.6 | 236.4 | 3.0 |
| 31 | LPC(22:4)>LPC(22:4/0:0)_and_LPC(0:0/22:4) | LPC | 572.3710 | 572.3710 | 0.1 | 2.16 | 245.7 | 238.6 | 3.0 |
| 32 | LPC(22:0)>LPC(22:0/0:0)_and_LPC(0:0/22:0) | LPC | 580.4339 | 580.4346 | 0.2 | 1.95 | 252.0 | 253.9 | -0.7 |
| 33 | PC(28:0) | PC | 678.5071 | 678.5068 | 0.5 | 1.77 | 274.6 | 275.2 | -0.2 |
| 34 | PC(30:1) | PC | 704.5226 | 704.5225 | 0.2 | 1.84 | 283.1 | 277.2 | 2.1 |
| 35 | PC(30:0) | PC | 706.5380 | 706.5382 | 0.3 | 3.53 | 280.0 | 280.5 | -0.2 |
| 36 | PC(O-32:2) | PC | 716.5584 | 716.5588 | 0.3 | 4.82 | 284.5 | 282.4 | 0.7 |
| 37 | PC(31:1) | PC | 718.5385 | 718.5383 | 0.5 | 4.17 | 291.8 | 279.7 | 4.3 |
| 38 | PC(O-32:1) | PC | 718.5747 | 718.5748 | 2.6 | 4.88 | 283.5 | 285.2 | -0.6 |
| 39 | PC(O-32:0) | PC | 720.5905 | 720.5898 | 3.3 | 4.33 | 284.3 | 287.9 | -1.2 |
| 40 | PC(32:3) | PC | 728.5229 | 728.5231 | 1.0 | 4.05 | 285.4 | 277.1 | 3.0 |
| 41 | PC(32:2) | PC | 730.5385 | 730.5386 | 0.1 | 3.88 | 285.4 | 279.6 | 2.1 |
| 42 | PC(32:1) | PC | 732.5540 | 732.5543 | 2.5 | 5.69 | 285.3 | 283.3 | 0.7 |
| 43 | PC(32:0) | PC | 734.5697 | 734.5696 | 0.2 | 5.77 | 283.8 | 286.6 | -1.0 |
| 44 | PC(33:2) | PC | 744.5543 | 744.5541 | 2.5 | 5.11 | 290.0 | 280.6 | 3.4 |
| 45 | PC(O-34:2) | PC | 744.5898 | 744.5890 | 0.5 | 4.58 | 289.7 | 287.6 | 0.7 |
| 46 | PC(33:1) | PC | 746.5696 | 746.5697 | 0.8 | 4.79 | 287.6 | 286.7 | 0.3 |
| 47 | PC(O-34:1) | PC | 746.6062 | 746.6054 | 2.1 | 4.54 | 290.8 | 291 | -0.1 |
| 48 | PC(O-34:0) | PC | 748.6220 | 748.6205 | 0.2 | 4.21 | 293.0 | 294.4 | -0.5 |

| | | | | | | | | | |
|---|---|---|---|---|---|---|---|---|---|
| 49 | PC(34:4) | PC | 754.5384 | 754.5381 | 0.4 | 3.67 | 292.5 | 281.9 | 3.8 |
| 50 | PC(34:3) | PC | 756.5540 | 756.5540 | 0.4 | 4.09 | 292.6 | 283.3 | 3.3 |
| 51 | PC(34:2) | PC | 758.5694 | 758.5718 | 0.2 | 3.59 | 290.0 | 286.4 | 1.3 |
| 52 | PC(34:1) | PC | 760.5852 | 760.5870 | 0.2 | 3.7 | 293.1 | 289.7 | 1.2 |
| 53 | PC(34:0) | PC | 762.6010 | 762.5990 | 0.4 | 3.36 | 297.8 | 292.8 | 1.7 |
| 54 | PC(O-36:5) | PC | 766.5751 | 766.5747 | 0.1 | 4.76 | 300.3 | 289.5 | 3.7 |
| 55 | PC(35:4) | PC | 768.5539 | 768.5540 | 0.0 | 4.01 | 297.6 | 284.3 | 4.7 |
| 56 | PC(35:4) | PC | 768.5539 | 768.5540 | 0.1 | 3.9 | 296.9 | 286.2 | 3.7 |
| 57 | PC(O-36:4) | PC | 768.5902 | 768.5895 | 0.4 | 3.62 | 294.8 | 291.7 | 1.1 |
| 58 | PC(35:3) | PC | 770.5694 | 770.5696 | 0.3 | 5.32 | 300.5 | 288 | 4.3 |
| 59 | PC(O-36:3) | PC | 770.6057 | 770.6059 | 0.7 | 4.58 | 296.5 | 291.8 | 1.6 |
| 60 | PC(35:2) | PC | 772.5851 | 772.5857 | 0.3 | 4.15 | 293.2 | 289.4 | 1.3 |
| 61 | PC(O-36:2) | PC | 772.6216 | 772.6210 | 0.3 | 3.9 | 300.0 | 294.3 | 1.9 |
| 62 | PC(35:1) | PC | 774.6008 | 774.6011 | 0.3 | 3.83 | 306.3 | 292.4 | 4.8 |
| 63 | PC(O-36:1) | PC | 774.6371 | 774.6368 | 1.3 | 6.22 | 301.8 | 297.4 | 1.5 |
| 64 | PC(36:5) | PC | 780.5540 | 780.5542 | 1.1 | 5.37 | 297.9 | 287.3 | 3.7 |
| 65 | PC(O-37:5) | PC | 780.5902 | 780.5903 | 0.1 | 4.47 | 299.4 | 290.7 | 3.0 |
| 66 | PC(36:4) | PC | 782.5694 | 782.5701 | 0.0 | 4.14 | 298.1 | 290.4 | 2.7 |
| 67 | PC(36:4) | PC | 782.5696 | 782.5695 | 0.8 | 6.51 | 297.3 | 287.5 | 3.4 |
| 68 | PC(36:3) | PC | 784.5851 | 784.5867 | 1.1 | 5.89 | 295.8 | 291.1 | 1.6 |
| 69 | PC(36:2) | PC | 786.6010 | 786.6026 | 0.0 | 4.26 | 297.1 | 292.7 | 1.5 |
| 70 | PC(36:1) | PC | 788.6166 | 788.6167 | 0.3 | 4.05 | 300.1 | 295.4 | 1.6 |
| 71 | PC(O-38:5) | PC | 794.6051 | 794.6055 | 0.7 | 3.79 | 302.3 | 295.7 | 2.2 |
| 72 | PC(O-38:4) | PC | 796.6217 | 796.6210 | 0.5 | 5.44 | 306.7 | 297.5 | 3.1 |
| 73 | PC(37:2) | PC | 800.6160 | 800.6174 | 0.3 | 4.42 | 301.5 | 295.5 | 2.0 |
| 74 | PC(37:1) | PC | 802.6323 | 802.6332 | 0.3 | 5.9 | 306.4 | 298.3 | 2.7 |
| 75 | PC(36:3) | PC | 806.5695 | 806.5692 | 0.7 | 5.32 | 305.6 | 293.1 | 4.3 |
| 76 | PC(38:6) | PC | 806.5695 | 806.5689 | 0.1 | 4.2 | 305.7 | 291.4 | 4.9 |
| 77 | PC(38:5) | PC | 808.5850 | 808.5848 | 1.4 | 6.26 | 301.8 | 294.2 | 2.6 |
| 78 | PC(38:4) | PC | 810.6008 | 810.6011 | 0.0 | 5.83 | 304.7 | 296.7 | 2.7 |
| 79 | PC(38:3) | PC | 812.6165 | 812.6163 | 0.8 | 5.38 | 307.1 | 297 | 3.4 |
| 80 | PC(38:2) | PC | 814.6321 | 814.6311 | 0.0 | 4.66 | 307.9 | 298.4 | 3.2 |
| 81 | PC(38:1) | PC | 816.6475 | 816.6468 | 0.2 | 4.53 | 307.7 | 300.8 | 2.3 |
| 82 | PC(O-40:7) | PC | 818.6057 | 818.6057 | 1.9 | 6.3 | 313.2 | 298 | 5.1 |
| 83 | PC(39:6) | PC | 820.5850 | 820.5850 | 1.3 | 5.46 | 308.1 | 296.6 | 3.9 |
| 84 | PC(O-40:6) | PC | 820.6211 | 820.6209 | 1.6 | 5.32 | 315.0 | 298.9 | 5.4 |
| 85 | PC(39:4) | PC | 824.6166 | 824.6160 | 1.0 | 6.36 | 310.1 | 299.7 | 3.5 |
| 86 | PC(O-40:4) | PC | 824.6523 | 824.6529 | 0.9 | 5.7 | 321.5 | 302.8 | 6.2 |
| 87 | PC(40:8) | PC | 830.5696 | 830.5690 | 0.1 | 5.03 | 312.5 | 294 | 6.3 |
| 88 | PC(40:7) | PC | 832.5848 | 832.5846 | 1.0 | 4.7 | 312.6 | 296.8 | 5.3 |
| 89 | PC(40:6) | PC | 834.6007 | 834.6002 | 0.4 | 4.21 | 312.0 | 299.5 | 4.2 |
| 90 | PC(42:7) | PC | 838.6320 | 838.6315 | 0.0 | 6.47 | 314.8 | 301.8 | 4.3 |
| 91 | PC(O-42:6) | PC | 848.6526 | 848.6528 | 0.9 | 5.57 | 319.9 | 304.2 | 5.1 |
| 92 | PC(42:9) | PC | 856.5850 | 856.5846 | 0.4 | 5.11 | 321.5 | 297.6 | 8.0 |
| 93 | PC(42:5) | PC | 864.6475 | 864.6473 | 0.7 | 6.44 | 325.8 | 305.5 | 6.7 |
| 94 | PC(42:4) | PC | 866.6638 | 866.6643 | 0.3 | 5.35 | 325.0 | 308.8 | 5.3 |
| 95 | PE(40:5) | PE | 794.5693 | 794.5694 | 0.1 | 4.53 | 303.4 | 291.1 | 4.2 |
| 96 | SM(d32:2) | SM | 673.5282 | 673.5277 | 0.2 | 6.24 | 271.6 | 276.9 | -1.9 |
| 97 | SM(d32:1) | SM | 675.5438 | 675.5447 | 0.1 | 3.92 | 271.5 | 280.5 | -3.2 |

| | | | | | | | | | |
|---|---|---|---|---|---|---|---|---|---|
| 98 | SM(d32:0) | SM | 677.5598 | 677.5582 | 2.0 | 5.2 | 278.2 | 285 | -2.4 |
| 99 | SM(d33:1) | SM | 689.5595 | 689.5599 | 0.5 | 3.04 | 277.3 | 283.8 | -2.3 |
| 100 | SM(d34:2) | SM | 701.5595 | 701.5615 | 4.2 | 4.06 | 278.1 | 283.5 | -1.9 |
| 101 | SM(d34:1) | SM | 703.5750 | 703.5778 | 3.0 | 3.58 | 277.6 | 286.3 | -3.0 |
| 102 | SM(d34:0) | SM | 705.5904 | 705.5891 | 0.7 | 4.81 | 281.2 | 290.1 | -3.1 |
| 103 | SM(d35:2) | SM | 715.5750 | 715.5748 | 0.1 | 4.19 | 286.7 | 286.6 | 0.0 |
| 104 | SM(d35:1) | SM | 717.5906 | 717.5905 | 2.1 | 6.46 | 283.3 | 289.8 | -2.2 |
| 105 | SM(d34:0-OH) | SM | 721.5858 | 721.5842 | 0.2 | 3.24 | 287.1 | 290.4 | -1.1 |
| 106 | SM(d36:3) | SM | 727.5753 | 727.5748 | 1.9 | 3.7 | 285.3 | 286.8 | -0.5 |
| 107 | SM(d36:2) | SM | 729.5908 | 729.5904 | 1.4 | 3.5 | 285.7 | 289.7 | -1.4 |
| 108 | SM(d36:1) | SM | 731.6065 | 731.6062 | 0.7 | 3.1 | 284.2 | 292.6 | -2.9 |
| 109 | SM(d36:0) | SM | 733.6223 | 733.6208 | 0.8 | 3.76 | 293.3 | 295.7 | -0.8 |
| 110 | SM(d37:1) | SM | 745.6221 | 745.6221 | 1.6 | 4.33 | 293.9 | 295.2 | -0.4 |
| 111 | SM(d38:2) | SM | 757.6218 | 757.6216 | 0.2 | 4.43 | 291.0 | 295.6 | -1.6 |
| 112 | SM(d38:1) | SM | 759.6378 | 759.6373 | 0.3 | 3.83 | 292.9 | 298.2 | -1.8 |
| 113 | SM(d39:1) | SM | 773.6533 | 773.6531 | 1.3 | 5.11 | 296.5 | 300.8 | -1.4 |
| 114 | SM(d40:2) | SM | 785.6533 | 785.6531 | 0.7 | 3.72 | 301.7 | 301.2 | 0.2 |
| 115 | SM(d40:1) | SM | 787.6690 | 787.6697 | 0.2 | 5.29 | 302.4 | 303 | -0.2 |
| 116 | SM(d41:2) | SM | 799.6690 | 799.6687 | 0.3 | 5.77 | 303.7 | 302.8 | 0.3 |
| 117 | SM(d41:1) | SM | 801.6846 | 801.6854 | 0.2 | 6.23 | 304.1 | 306.5 | -0.8 |
| 118 | SM(d42:3) | SM | 811.6686 | 811.6685 | 1.1 | 6.54 | 308.7 | 303.9 | 1.6 |
| 119 | SM(d42:2) | SM | 813.6845 | 813.6862 | 0.2 | 5.91 | 309.2 | 306.6 | 0.8 |
| 120 | SM(d43:3) | SM | 825.6846 | 825.6846 | 1.1 | 6.75 | 311.9 | 306 | 1.9 |
| 121 | SM(d43:2) | SM | 827.7000 | 827.6999 | 0.3 | 6.1 | 313.8 | 310.1 | 1.2 |
| 122 | SM(d44:3) | SM | 839.6996 | 839.6997 | 0.2 | 5.8 | 315.6 | 310.6 | 1.6 |
| 123 | TG(44:1)>TG(10:0_16:0_18:1) M + NH3 | TG | 766.6920 | 766.6927 | 0.1 | 6.62 | 296.3 | 299.8 | -1.2 |
| 124 | TG(46:2) M + NH3 | TG | 792.7078 | 792.7083 | 0.1 | 6.24 | 302.3 | 303.3 | -0.3 |
| 125 | TG(46:1)>TG(12:0_16:0_18:1)_and_TG(10:0_18:0_18:1)_and_TG(14:0_16:0_16:1) M + NH3 | TG | 794.7234 | 794.7233 | 0.1 | 6.56 | 302.1 | 305.5 | -1.1 |
| 126 | TG(46:0)>TG(14:0_16:0_16:0)_and_TG(12:0_16:0_18:0) M + NH3 | TG | 796.7388 | 796.7391 | 0.8 | 7.76 | 309.5 | 307.8 | 0.5 |
| 127 | TG(48:3) M + NH3 | TG | 818.7234 | 818.7238 | 0.7 | 7.79 | 309.7 | 307.4 | 0.7 |
| 128 | TG(48:2)>TG(14:0_16:0_18:2)_and_TG(14:0_16:1_18:1) M + NH3 | TG | 820.7391 | 820.7395 | 0.1 | 8.44 | 309.3 | 309.3 | 0.0 |
| 129 | TG(48:1)>TG(16:0_16:0_16:1)_and_TG(14:0_16:0_18:1) M + NH3 | TG | 822.7548 | 822.7547 | 0.1 | 8.07 | 309.4 | 311.6 | -0.7 |
| 130 | TG(49:3) M + NH3 | TG | 832.7389 | 832.7390 | 0.1 | 8.42 | 315.1 | 310.9 | 1.3 |
| 131 | TG(49:2) M + NH3 | TG | 834.7545 | 834.7550 | 0.5 | 8.1 | 315.2 | 312.2 | 1.0 |
| 132 | TG(49:1)>TG(15:0_16:0_18:0)_and_TG(16:0_16:0_17:1)_and_TG(16:0_16:1_17:0) M + NH3 | TG | 836.7702 | 836.7703 | 0.9 | 7.86 | 319.5 | 314.6 | 1.6 |
| 133 | TG(50:5) M + NH3 | TG | 842.7190 | 842.7231 | 0.2 | 8.61 | 315.7 | 310.8 | 1.6 |
| 134 | TG(50:4)>TG(16:1_16:1_18:2)_and_TG(16:1_16:1_18:2) M + NH3 | TG | 844.7388 | 844.7396 | 0.1 | 7.96 | 315.4 | 311.6 | 1.2 |
| 135 | TG(50:3) M + NH3 | TG | 846.7546 | 846.7556 | 0.1 | 8.01 | 316.2 | 313.5 | 0.9 |
| 136 | TG(50:2)>TG(16:0_16:1_18:1)_and_TG(14:0_18:1_18:1) M + NH3 | TG | 848.7701 | 848.7708 | 0.9 | 7.75 | 317.4 | 315.3 | 0.7 |
| 137 | TG(50:1)>TG(16:0_16:0_18:1)_M+NH3 | TG | 850.7858 | 850.7857 | 0.3 | 7.75 | 316.8 | 318 | -0.4 |
| 138 | TG(51:4)>TG(16:1_17:1_18:2)_and_TG(15:0_18:2_18:2)_and_TG(15:1_18:1_18:2) M + NH3 | TG | 858.7545 | 858.7544 | 0.1 | 8.81 | 319.8 | 314.9 | 1.5 |

| | | | | | | | | | |
|---|---|---|---|---|---|---|---|---|---|
| 139 | TG(51:2)>TG(15:0_18:1_18:1)_and_TG(16:0_17:1_18:1)_and_TG(16:1_17:0_18:1) M + NH3 | TG | 862.7857 | 862.7861 | 0.8 | 8.42 | 320.2 | 318.2 | 0.6 |
| 140 | TG(51:1) M + NH3 | TG | 864.8014 | 864.8019 | 1.3 | 9.22 | 321.1 | 320.8 | 0.1 |
| 141 | TG(52:6) M + NH3 | TG | 868.7388 | 868.7387 | 0.8 | 8.42 | 319.4 | 314.8 | 1.5 |
| 142 | TG(52:5)>TG(16:0_18:2_18:3) M + NH3 | TG | 870.7545 | 870.7554 | 0.1 | 7.79 | 320.4 | 316.2 | 1.3 |
| 143 | TG(52:4)>TG(16:0_16:0_20:4)_and_TG(16:1_18:1_18:2)_and_TG(16:0_18:1_18:2) M + NH3 | TG | 872.7701 | 872.7720 | 0.7 | 9.22 | 320.8 | 317.7 | 1.0 |
| 144 | TG(52:3)>TG(16:0_18:1_18:2)_and_TG(16:1_18:1_18:1) M + NH3 | TG | 874.7858 | 874.7875 | 1.0 | 8.79 | 321.5 | 320 | 0.5 |
| 145 | TG(52:2)>TG(16:0_18:1_18:1) M + NH3 | TG | 876.8016 | 876.8023 | 2.0 | 8.48 | 337.1 | 321.9 | 4.7 |
| 146 | TG(52:1) M + NH3 | TG | 878.8170 | 878.8164 | 2.0 | 8.18 | 324.3 | 324 | 0.1 |
| 147 | TG(53:5)>TG(17:1_18:2_18:2) M + NH3 | TG | 884.7701 | 884.7688 | 0.8 | 7.85 | 329.6 | 319.5 | 3.2 |
| 148 | TG(53:4)>TG(17:0_18:2_18:2)_and_TG(17:1_18:1_18:2) M + NH3 | TG | 886.7856 | 886.7852 | 1.2 | 8.12 | 329.7 | 321.1 | 2.7 |
| 149 | TG(53:3)>TG(17:0_18:1_18:2)_and_TG(17:1_18:1_18:1)_and_TG(16:1_18:1_19:1)_and_TG(16:0_18:1_19:2) M + NH3 | TG | 888.8013 | 888.8012 | 0.5 | 7.81 | 330.1 | 323.1 | 2.2 |
| 150 | TG(53:2) M + NH3 | TG | 890.8167 | 890.8168 | 1.0 | 7.94 | 334.1 | 324.9 | 2.8 |
| 151 | TG(54:7)>TG(16:0_16:1_22:6)_and_TG(16:0_18:2_20:5) M + NH3 | TG | 894.7543 | 894.7539 | 0.1 | 7.89 | 328.1 | 318.5 | 3.0 |
| 152 | TG(52:3)>TG(16:0_18:1_18:2)_and_TG(16:1_18:1_18:1) M + NH3 | TG | 896.7701 | 896.7699 | 0.2 | 9 | 329.5 | 320.4 | 2.9 |
| 153 | TG(54:5)>TG(18:1_18:2_18:2)_and_TG(16:0_16:0_22:5)_and_TG(16:0_18:1_20:4) M + NH3 | TG | 898.7858 | 898.7855 | 0.0 | 8.59 | 329.9 | 322 | 2.5 |
| 154 | TG(54:4)>TG(18:1_18:1_18:2) M + NH3 | TG | 900.8012 | 900.8010 | 0.6 | 8.3 | 331.5 | 323.8 | 2.4 |
| 155 | TG(54:3)>TG(18:1/18:1/18:1) M + NH3 | TG | 902.8167 | 902.8167 | 1.4 | 8.02 | 332.1 | 325.8 | 1.9 |
| 156 | TG(54:2)>TG(16:0_18:1_20:1)_and_TG(18:0_18:1_18:1) M + NH3 | TG | 904.8326 | 904.8316 | 0.0 | 7.91 | 335.1 | 327.7 | 2.3 |
| 157 | TG(56:6)>TG(18:1_18:1_20:4)_and_TG(16:0_18:1_22:5)_and_TG(18:1_18:2_20:3) M + NH3 | TG | 924.8013 | 924.8006 | 1.1 | 9.2 | 341.2 | 327.6 | 4.2 |
| 158 | TG(56:5)>TG(16:0_18:1_22:4) M + NH3 | TG | 926.8172 | 926.8162 | 0.2 | 8.87 | 344.6 | 329.2 | 4.7 |
| 159 | TG(56:1) M + NH3 | TG | 934.8792 | 934.8793 | 0.1 | 8.47 | 346.9 | 335.2 | 3.5 |
| 160 | TG(58:10) M + NH3 | TG | 944.7701 | 944.7694 | 0.4 | 8.12 | 343.6 | 326.7 | 5.2 |
| 161 | TG(57:2) M + NH3 | TG | 946.8793 | 946.8787 | 0.6 | 8.36 | 345.6 | 335.8 | 2.9 |
| 162 | TG(57:1)>TG(16:0_18:1_23:0)_and_TG(18:0_18:1_21:0) M + NH3 | TG | 948.8951 | 948.8944 | 1.0 | 8.8 | 350.6 | 337.2 | 4.0 |
| 163 | TG(58:3) M + NH3 | TG | 958.8792 | 958.8787 | 0.6 | 8.79 | 346.3 | 336.4 | 2.9 |
| 164 | TG(58:2) M + NH3 | TG | 960.8950 | 960.8946 | 0.2 | 8.36 | 346.2 | 338.4 | 2.3 |

**Supplementary Table S4. Summary of lipid features with matched $^{Orbi}$CCS and $^{DT}$CCS values from a single LC-Orbitrap experiment (pump speed 68%, resolution setting 240,000). The same dataset present in Fig. 3a-f.**

| # | Name | Type | m/z_this study | m/z_database | mzError_ppm | RT (min) | Orbi_CCS | DT_CCS | % error |
|---|---|---|---|---|---|---|---|---|---|
| 1 | LPC(16:0)>LPC(16:0/0:0)_and_LPC(0:0/16:0) | LPC | 496.33994 | 496.3403 | 0.7 | 2.77 | 218.2 | 231.4 | -5.7 |
| 2 | LPC(18:0)>LPC(18:0/0:0)_and_LPC(0:0/18:0) | LPC | 524.37122 | 524.3716 | 0.7 | 2.77 | 228.4 | 238.8 | -4.3 |
| 3 | LPC(20:0)>LPC(20:0/0:0)_and_LPC(0:0/20:0) | LPC | 552.40259 | 552.4029 | 0.6 | 2.74 | 242.6 | 247 | -1.8 |
| 4 | Car(12:0) | Car | 344.27948 | 344.2782 | 3.7 | 1.23 | 172.1 | 199.5 | -13.7 |
| 5 | Car(16:0) | Car | 400.3421 | 400.3427 | 1.5 | 2.05 | 187.2 | 214.7 | -12.8 |
| 6 | LPC(14:0)>LPC(14:0/0:0)_and_LPC(0:0/14:0) | LPC | 468.3085 | 468.309 | 1.1 | 1.57 | 212.5 | 220.8 | -3.8 |
| 7 | LPC(18:1)>LPC(18:1/0:0)_and_LPC(0:0/18:1) | LPC | 522.35547 | 522.3559 | 0.8 | 2.17 | 229.3 | 233.2 | -1.7 |
| 8 | LPC(19:0)>LPC(19:0/0:0)_and_LPC(0:0/19:0) | LPC | 538.38696 | 538.3872 | 0.4 | 2.54 | 240.0 | 240.5 | -0.2 |
| 9 | PC(32:2) | PC | 730.53845 | 730.5387 | 0.3 | 3.70 | 283.4 | 278.4 | 1.8 |
| 10 | PC(34:1) | PC | 760.58502 | 760.5856 | 0.8 | 4.88 | 292.9 | 287.9 | 1.7 |
| 11 | PC(36:2) | PC | 786.60065 | 786.6012 | 0.7 | 5.11 | 294.2 | 293.3 | 0.3 |
| 12 | PC(32:1) | PC | 732.55389 | 732.5543 | 0.6 | 4.17 | 283.4 | 277.6 | 2.1 |
| 13 | PC(34:2) | PC | 758.56927 | 758.5699 | 0.8 | 4.33 | 290.7 | 280.6 | 3.6 |
| 14 | PC(34:3) | PC | 756.55396 | 756.5543 | 0.4 | 3.90 | 290.2 | 278.2 | 4.3 |
| 15 | PC(35:1) | PC | 774.60089 | 774.6013 | 0.5 | 5.32 | 292.8 | 285.8 | 2.5 |
| 16 | PC(36:1) | PC | 788.61652 | 788.6169 | 0.5 | 5.77 | 294.3 | 289.4 | 1.7 |
| 17 | PC(36:2) | PC | 786.60065 | 786.6012 | 0.7 | 4.96 | 294.2 | 287.1 | 2.5 |
| 18 | PE(O-36:3)>PE(O-18:1/18:2) | PE | 728.55927 | 728.5594 | 0.2 | 5.59 | 286.5 | 273.5 | 4.8 |
| 19 | SM(d34:1) | SM | 703.57489 | 703.5754 | 0.7 | 4.06 | 276.6 | 281.2 | -1.6 |
| 20 | SM(d36:1) | SM | 731.60669 | 731.6067 | 0.0 | 4.81 | 284.1 | 288.4 | -1.5 |
| 21 | SM(d36:2) | SM | 729.59052 | 729.591 | 0.7 | 4.19 | 285.4 | 285.3 | 0.1 |
| 22 | SM(d37:1) | SM | 745.62195 | 745.6223 | 0.5 | 5.29 | 290.8 | 289.8 | 0.4 |
| 23 | SM(d38:1) | SM | 759.63751 | 759.638 | 0.6 | 5.77 | 292.3 | 293.4 | -0.4 |
| 24 | SM(d39:1) | SM | 773.65338 | 773.6536 | 0.3 | 5.97 | 294.9 | 297 | -0.7 |
| 25 | SM(d40:1) | SM | 787.66882 | 787.6693 | 0.6 | 6.54 | 298.5 | 299.1 | -0.2 |
| 26 | SM(d40:2) | SM | 785.65332 | 785.6536 | 0.4 | 5.91 | 298.5 | 296.8 | 0.6 |
| 27 | SM(d41:1) | SM | 801.68457 | 801.6849 | 0.4 | 6.75 | 300.4 | 302.3 | -0.6 |
| 28 | SM(d41:2) | SM | 799.66901 | 799.6693 | 0.4 | 6.34 | 301.1 | 300.1 | 0.3 |
| 29 | SM(d42:2) | SM | 813.68445 | 813.6849 | 0.6 | 6.46 | 304.5 | 302.2 | 0.8 |
| 30 | SM(d42:3) | SM | 811.6687 | 811.6693 | 0.7 | 5.80 | 304.6 | 300.8 | 1.3 |
| 31 | SM(d43:2) | SM | 827.70013 | 827.7006 | 0.6 | 6.81 | 307.8 | 305.7 | 0.7 |

**Supplementary Table S5. Isotopically labeled lipid standards.** Lipid names, CAS numbers, and concentrations of isotopically labeled standards purchased from Avanti Polar Lipids (Alabaster, AL, USA). The internal standard stock solution was prepared in-house and used for quantification and normalization in the reverse-phase UHPLC–MS workflow.

| Lipid Subclass Name | Isotopically Labeled Lipids | CAS Number | Concentration (μg $mL^{-1}$) |
|---|---|---|---|
| Cholesterol Ester | CE (18:1(d7)) | 1416275-35-7 | 350 |
| Cholesterol | Cholesterol-d7 | 83199-47-7 | 100 |
| Diacylglycerol | DG (15:0/18:1(d7)) | 2097561-14-1 | 10 |
| Lysophosphatidylcholine | LPC (18:1(d7)) | 2097561-13-0 | 25 |
| Lysophosphatidylethanolamine | LPE (18:1(d7)) | 2260669-47-2 | 5 |
| Phosphatidylcholine | PC (15:0/18:1(d7)) | 2097561-16-3 | 160 |
| Phosphatidylethanolamine | PE (15:0/18:1(d7)) | 2097561-15-2 | 5 |
| Phosphatidylglycerol | PG (15:0/18:1(d7)) | 2260669-42-7 | 30 |
| Phosphatidylinositol | PI (15:0/18:1(d7)) | 2260669-44-9 | 20 |
| Phosphatidylserine | PS (15:0/18:1(d7)) | 2260669-40-5 | 10 |
| Sphingomyelin | SM (d18:1/(d9)) | 2260669-50-7 | 30 |
| Triacylglycerol | TG (15:0/18:1(d7)/15:0) | 2097561-17-4 | 55 |

**Supplementary Figure S6. Slope and intercept values used to calibrate $^{Orbi}$CCS during an LC experiment.** Panel (a) shows the changing ion gauge pressure reading when the experiment was performed at reduced turbopump speed. Panel (b) and (c) show the corresponding RT-specific calibration factors, $a_{RT}$ and $b_{RT}$ from linear regression analysis between $^{DT}CCS$ values of internal standards and their corresponding $^{Orbi}CCS_{LC,IS}$.

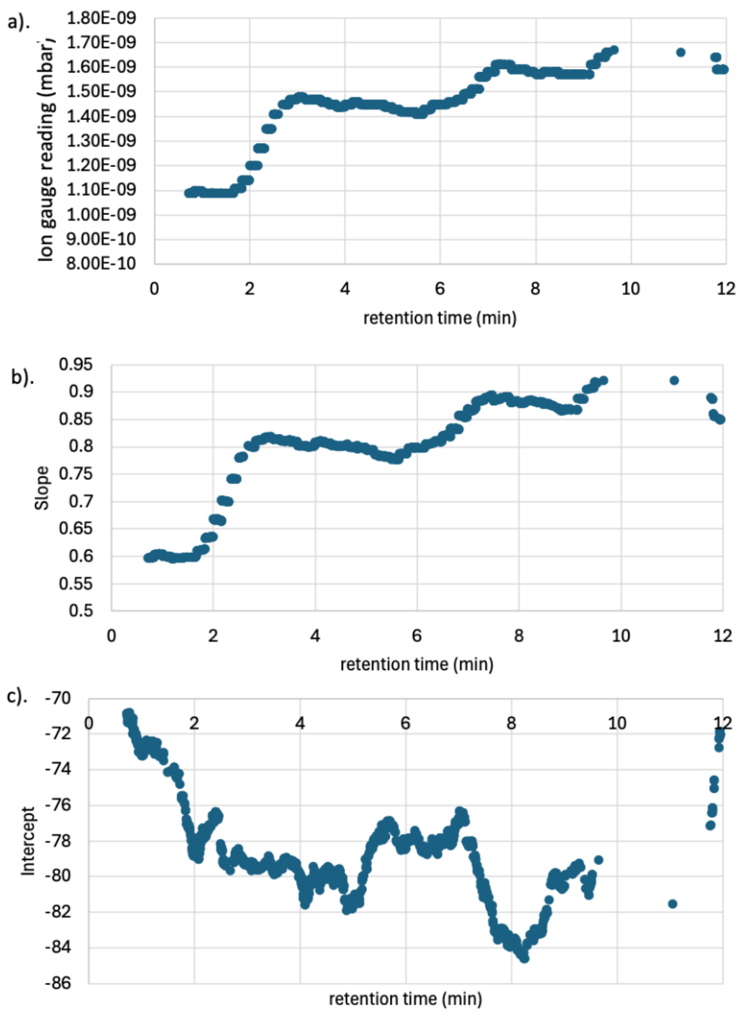

**Supplementary Table S7. Summary of lipid features with matched $^{Orbi}$CCS to reference $^{TIMS}$CCS dataset for experiments conducted at a resolution of 240,000 and turbopump speed 68%.**

| # | m/z_this study | m/z_database | mzError_ppm | $^{Orbi}$CCS_this study | $^{TIMS}$CCS_database | Name_database | Name_This study | % error |
|---|---|---|---|---|---|---|---|---|
| 1 | 818.7234 | 818.7238 | 0.5 | 307.4 | 307.4 | TG 16:1_16:1_16:1 | TG(48:3) M + NH3 | 0.0 |
| 2 | 796.7390 | 796.7391 | 0.1 | 307.9 | 307.8 | TG 14:0_16:0_16:0 | TG(46:0)>TG(14:0_16:0_16:0)_and_TG(12:0_16:0_18:0) M + NH3 | 0.0 |
| 3 | 580.4340 | 580.4346 | 1.1 | 254.0 | 253.9 | LPC 22:0-SN1 | LPC(22:0)>LPC(22:0/0:0)_and_LPC(0:0/22:0) | 0.0 |
| 4 | 846.7546 | 846.7556 | 1.2 | 313.4 | 313.5 | TG 16:1_16:0_18:2 | TG(50:3) M + NH3 | 0.0 |
| 5 | 716.5590 | 716.5588 | 0.2 | 282.3 | 282.4 | PC O-32:2 | PC(O-32:2) | 0.0 |
| 6 | 732.5539 | 732.5543 | 0.5 | 283.4 | 283.3 | PC 16:0_16:1 | PC(32:1) | 0.1 |
| 7 | 872.7703 | 872.7720 | 2.0 | 317.9 | 317.7 | TG 16:0_18:2_18:2 | TG(52:4)>TG(16:0_16:0_20:4)_and_TG(16:1_18:1_18:2)_and_TG(16:0_18:1_18:2) M + NH3 | 0.1 |
| 8 | 746.5698 | 746.5697 | 0.1 | 287.0 | 286.7 | PC 33:1 | PC(33:1) | 0.1 |
| 9 | 796.6216 | 796.6210 | 0.9 | 297.8 | 297.5 | PC O-38:4 | PC(O-38:4) | 0.1 |
| 10 | 820.7391 | 820.7395 | 0.5 | 309.6 | 309.3 | TG 14:0_16:1_18:1 | TG(48:2)>TG(14:0_16:0_18:2)_and_TG(14:0_16:1_18:1) M + NH3 | 0.1 |
| 11 | 794.7234 | 794.7233 | 0.1 | 305.2 | 305.5 | TG 14:0_16:0_16:1 | TG(46:1)>TG(12:0_16:0_18:1)_and_TG(10:0_18:0_18:1)_and_TG(14:0_16:0_16:1) M + NH3 | -0.1 |
| 12 | 774.6009 | 774.6011 | 0.3 | 292.8 | 292.4 | PC 35:1 | PC(35:1) | 0.2 |
| 13 | 792.7077 | 792.7083 | 0.7 | 302.8 | 303.3 | TG 12:0_16:1_18:1 | TG(46:2) M + NH3 | -0.2 |
| 14 | 848.7701 | 848.7708 | 0.8 | 314.7 | 315.3 | TG 16:0_16:1_18:1 | TG(50:2)>TG(16:0_16:1_18:1)_and_TG(14:0_18:1_18:1) M + NH3 | -0.2 |
| 15 | 770.6058 | 770.6059 | 0.1 | 292.4 | 291.8 | PC O-36:3 | PC(O-36:3) | 0.2 |
| 16 | 900.8010 | 900.8010 | 0.1 | 324.4 | 323.8 | TG 18:1_18:1_18:2 | TG(54:4)>TG(18:1_18:1_18:2) M + NH3 | 0.2 |
| 17 | 520.3397 | 520.3405 | 1.5 | 229.6 | 229.1 | LPC 18:2-SN1 | LPC(18:2)>LPC(18:2/0:0)_and_LPC(0:0/18:2) | 0.2 |
| 18 | 811.6687 | 811.6685 | 0.2 | 304.6 | 303.9 | SM d42:3 | SM(d42:3) | 0.2 |
| 19 | 744.5902 | 744.5890 | 1.6 | 288.3 | 287.6 | PC O-34:2 | PC(O-34:2) | 0.2 |
| 20 | 844.7388 | 844.7396 | 0.9 | 312.4 | 311.6 | TG 14:0_18:2_18:2 | TG(50:4)>TG(16:1_16:1_18:2)_and_TG(16:1_16:1_18:2) M + NH3 | 0.3 |

| | | | | | | | | |
|---|---|---|---|---|---|---|---|---|
| 21 | 546.3556 | 546.3547 | 1.6 | 236.2 | 235.4 | LPC 20:3-SN2 | LPC(20:3)>LPC(20:3/0:0)_and_LPC(0:0/20:3) | 0.3 |
| 22 | 768.5903 | 768.5895 | 1.0 | 290.7 | 291.7 | PC O-36:4 | PC(O-36:4) | -0.3 |
| 23 | 902.8168 | 902.8167 | 0.2 | 324.7 | 325.8 | TG 18:1_18:1_18:1 | TG(54:3)>TG(18:0_18:1_18:2)_and_TG(18:1/18:1/18:1)_and_TG(16:0_18:2_20:1) M + NH3 | -0.4 |
| 24 | 788.6165 | 788.6167 | 0.2 | 294.3 | 295.4 | PC 18:0_18:1 | PC(36:1) | -0.4 |
| 25 | 870.7545 | 870.7554 | 1.0 | 317.4 | 316.2 | TG 16:1_18:2_18:2 | TG(52:5)>TG(16:0_18:2_18:3) M + NH3 | 0.4 |
| 26 | 850.7858 | 850.7857 | 0.1 | 316.8 | 318 | TG 16:0_16:0_18:1 | TG(50:1)>TG(16:0_16:0_18:1)_M+NH3 | -0.4 |
| 27 | 678.5070 | 678.5068 | 0.3 | 276.3 | 275.2 | PC 14:0_14:0 | PC(28:0) | 0.4 |
| 28 | 727.5753 | 727.5748 | 0.7 | 285.7 | 286.8 | SM d36:3 | SM(d36:3) | -0.4 |
| 29 | 550.3868 | 550.3868 | 0.1 | 241.4 | 242.4 | LPC 20:1-SN1 | LPC(20:1)>LPC(20:1/0:0)_and_LPC(0:0/20:1) | -0.4 |
| 30 | 830.7236 | 830.7228 | 0.9 | 308.0 | 309.4 | TG 15:1_16:1_18:2 | TG(49:4) M + NH3 | -0.4 |
| 31 | 799.6690 | 799.6687 | 0.3 | 301.4 | 302.8 | SM d41:2 | SM(d41:2) | -0.4 |
| 32 | 862.7860 | 862.7861 | 0.1 | 319.6 | 318.2 | TG 16:0_17:1_18:1 | TG(51:2) M + NH3 | 0.5 |
| 33 | 520.3398 | 520.3402 | 0.7 | 230.1 | 231.2 | LPC 18:2-SN2 | LPC(18:2)>LPC(18:2/0:0)_and_LPC(0:0/18:2) | -0.5 |
| 34 | 874.7858 | 874.7875 | 2.0 | 318.3 | 320 | TG 16:0_18:1_18:2 | TG(52:3)>TG(16:0_18:1_18:2)_and_TG(16:1_18:1_18:1) M + NH3 | -0.5 |
| 35 | 786.6007 | 786.6026 | 2.5 | 294.2 | 292.7 | PC 18:0_18:2 | PC(36:2) | 0.5 |
| 36 | 536.3713 | 536.3720 | 1.2 | 239.0 | 237.8 | LPC 19:1-SN1 | LPC(19:1)>LPC(19:1/0:0)_and_LPC(0:0/19:1) | 0.5 |
| 37 | 794.6052 | 794.6055 | 0.4 | 297.3 | 295.7 | PC O-38:5 | PC(O-38:5) | 0.5 |
| 38 | 827.7000 | 827.6999 | 0.1 | 308.4 | 310.1 | SM d43:2 | SM(d43:2) | -0.6 |
| 39 | 822.7546 | 822.7547 | 0.1 | 309.9 | 311.6 | TG 14:0_16:0_18:1 | TG(48:1)>TG(16:0_16:0_16:1)_and_TG(14:0_16:0_18:1) M + NH3 | -0.6 |
| 40 | 888.8013 | 888.8012 | 0.0 | 324.9 | 323.1 | TG 17:0_18:1_18:2 | TG(53:3)>TG(17:0_18:1_18:2)_and_TG(17:1_18:1_18:1)_and_TG(16:1_18:1_19:1)_and_TG(16:0_18:1_19:2) M + NH3 | 0.6 |
| 41 | 839.6998 | 839.6997 | 0.1 | 312.4 | 310.6 | SM d44:3 | SM(d44:3) | 0.6 |
| 42 | 898.7857 | 898.7855 | 0.2 | 324.0 | 322 | TG 18:1_18:2_18:2 | TG(54:5)>TG(18:1_18:2_18:2)_and_TG(16:0_16:0_22:5)_and_TG(16:0_18:1_20:4) M + NH3 | 0.6 |

| | | | | | | | | |
|---|---|---|---|---|---|---|---|---|
| 43 | 878.8170 | 878.8164 | 0.7 | 321.8 | 324 | TG 16:0_18:0_18:1 | TG(52:1) M + NH3 | -0.7 |
| 44 | 813.6845 | 813.6862 | 2.1 | 304.5 | 306.6 | SM d18:1_24:1 | SM(d42:2) | -0.7 |
| 45 | 868.7387 | 868.7387 | 0.1 | 317.0 | 314.8 | TG 16:1_18:2_18:3 | TG(52:6) M + NH3 | 0.7 |
| 46 | 762.6009 | 762.5990 | 2.5 | 290.6 | 292.8 | PC 18:0_16:0 | PC(34:0) | -0.8 |
| 47 | 825.6845 | 825.6846 | 0.1 | 308.3 | 306 | SM d43:3 | SM(d43:3) | 0.8 |
| 48 | 544.3397 | 544.3397 | 0.1 | 235.6 | 233.8 | LPC 20:4-SN2 | LPC(20:4)>LPC(20:4/0:0)_and_LPC(0:0/20:4) | 0.8 |
| 49 | 824.7708 | 824.7706 | 0.3 | 311.3 | 313.8 | TG 16:0_16:0_16:0 | TG(48:0)>TG(16:0/16:0/16:0)_and_TG(14:0_16:0_18:0) M + NH3 | -0.8 |
| 50 | 546.3555 | 546.3552 | 0.5 | 235.7 | 233.8 | LPC 20:3-SN1 | LPC(20:3)>LPC(20:3/0:0)_and_LPC(0:0/20:3) | 0.8 |
| 51 | 824.6523 | 824.6529 | 0.7 | 305.3 | 302.8 | PC O-40:4 | PC(O-40:4) | 0.8 |
| 52 | 706.5379 | 706.5382 | 0.4 | 278.1 | 280.5 | PC 30:0 | PC(30:0) | -0.8 |
| 53 | 864.8012 | 864.8019 | 0.8 | 323.6 | 320.8 | TG 16:0_17:0_18:1 | TG(51:1) M + NH3 | 0.9 |
| 54 | 784.5850 | 784.5867 | 2.1 | 293.6 | 291.1 | PC 18:1_18:2 | PC(36:3) | 0.9 |
| 55 | 876.8015 | 876.8023 | 1.0 | 319.1 | 321.9 | TG 16:0_18:1_18:1 | TG(52:2)>TG(16:0_18:1_18:1) M + NH3 | -0.9 |
| 56 | 858.7545 | 858.7544 | 0.1 | 317.7 | 314.9 | TG 15:0_18:2_18:2 | TG(51:4)>TG(16:1_17:1_18:2)_and_TG(15:0_18:2_18:2)_and_TG(15:1_18:1_18:2) M + NH3 | 0.9 |
| 57 | 832.7390 | 832.7390 | 0.1 | 308.1 | 310.9 | TG 15:0_16:1_18:2 | TG(49:3) M + NH3 | -0.9 |
| 58 | 785.6533 | 785.6531 | 0.2 | 298.5 | 301.2 | SM d40:2 | SM(d40:2) | -0.9 |
| 59 | 548.3712 | 548.3708 | 0.8 | 238.8 | 236.5 | LPC 20:2-SN1 | LPC(20:2)>LPC(20:2/0:0)_and_LPC(0:0/20:2) | 1.0 |
| 60 | 772.6217 | 772.6210 | 0.9 | 291.4 | 294.3 | PC O-36:2 | PC(O-36:2) | -1.0 |
| 61 | 766.6921 | 766.6927 | 0.8 | 296.8 | 299.8 | TG 12:0_16:0_16:1 | TG(44:1)>TG(10:0_16:0_18:1) M + NH3 | -1.0 |
| 62 | 896.7700 | 896.7699 | 0.0 | 323.6 | 320.4 | TG 18:2_18:2_18:2 | TG(54:6)>TG(16:0_18:2_20:4) M + NH3 | 1.0 |
| 63 | 718.5748 | 718.5748 | 0.0 | 282.3 | 285.2 | PC O-32:1 | PC(O-32:1) | -1.0 |
| 64 | 836.7701 | 836.7703 | 0.2 | 311.3 | 314.6 | TG 15:0_16:0_18:1 | TG(49:1)>TG(15:0_16:0_18:0)_and_TG(16:0_16:0_17:1)_and_TG(16:0_16:1_17:0) M + NH3 | -1.0 |

| | | | | | | | | |
|---|---|---|---|---|---|---|---|---|
| 65 | 746.6063 | 746.6054 | 1.3 | 287.9 | 291 | PC O-34:1 | PC(O-34:1) | -1.1 |
| 66 | 715.5750 | 715.5748 | 0.3 | 283.5 | 286.6 | SM d35:2 | SM(d35:2) | -1.1 |
| 67 | 904.8325 | 904.8316 | 1.1 | 331.2 | 327.7 | TG 18:0_18:1_18:1 | TG(54:2)>TG(16:0_18:1_20:1)_and_TG(18:0_18:1_18:1) M + NH3 | 1.1 |
| 68 | 772.5852 | 772.5857 | 0.7 | 292.5 | 289.4 | PC 35:2 | PC(35:2) | 1.1 |
| 69 | 886.7857 | 886.7852 | 0.6 | 324.6 | 321.1 | TG 17:1_18:1_18:2 | TG(53:4)>TG(17:0_18:2_18:2)_and_TG(17:1_18:1_18:2) M + NH3 | 1.1 |
| 70 | 760.5850 | 760.5870 | 2.6 | 292.9 | 289.7 | PC 16:0_18:1 | PC(34:1) | 1.1 |
| 71 | 734.5698 | 734.5696 | 0.3 | 283.0 | 286.6 | PC 16:0_16:0 | PC(32:0) | -1.3 |
| 72 | 810.6007 | 810.6011 | 0.5 | 300.4 | 296.7 | PC 18:0_20:4 | PC(38:4) | 1.3 |
| 73 | 518.3242 | 518.3248 | 1.1 | 230.8 | 227.9 | LPC 18:3-SN2 | LPC(18:3)>LPC(18:3/0:0)_and_LPC(0:0/18:3) | 1.3 |
| 74 | 882.7543 | 882.7543 | 0.0 | 322.5 | 318.4 | TG 17:1_18:2_18:3 | TG(53:6)>TG(17:2_18:2_18:2)_and_TG(17:1_18:2_18:3) M + NH3 | 1.3 |
| 75 | 538.3870 | 538.3872 | 0.4 | 240.0 | 243.2 | LPC 19:0-SN2 | LPC(19:0)>LPC(19:0/0:0)_and_LPC(0:0/19:0) | -1.3 |
| 76 | 796.5848 | 796.5848 | 0.1 | 297.7 | 293.8 | PC 37:4 | PC(37:4) | 1.3 |
| 77 | 730.5385 | 730.5386 | 0.2 | 283.4 | 279.6 | PC 32:2 | PC(32:2) | 1.3 |
| 78 | 782.5693 | 782.5701 | 1.0 | 294.4 | 290.4 | PC 16:0_20:4 | PC(36:4) | 1.4 |
| 79 | 721.5856 | 721.5842 | 2.0 | 286.3 | 290.4 | SM 34:0;3O | SM(d34:0-OH) | -1.4 |
| 80 | 924.8012 | 924.8006 | 0.6 | 332.3 | 327.6 | TG 18:1_18:1_20:4 | TG(56:6)>TG(18:1_18:1_20:4)_and_TG(16:0_18:1_22:5)_and_TG(18:1_18:2_20:3) M + NH3 | 1.4 |
| 81 | 884.7700 | 884.7688 | 1.4 | 324.1 | 319.5 | TG 17:1_18:2_18:2 | TG(53:5)>TG(17:1_18:2_18:2) M + NH3 | 1.4 |
| 82 | 729.5905 | 729.5904 | 0.1 | 285.4 | 289.7 | SM d18:1_18:1 | SM(d36:2) | -1.5 |
| 83 | 787.6688 | 787.6697 | 1.1 | 298.5 | 303 | SM d40:1 | SM(d40:1) | -1.5 |
| 84 | 814.6320 | 814.6311 | 1.1 | 302.8 | 298.4 | PC 38:2 | PC(38:2) | 1.5 |
| 85 | 745.6220 | 745.6221 | 0.2 | 290.8 | 295.2 | SM d37:1 | SM(d37:1) | -1.5 |
| 86 | 758.5693 | 758.5718 | 3.3 | 290.7 | 286.4 | PC 16:0_18:2 | PC(34:2) | 1.5 |
| 87 | 544.3398 | 544.3399 | 0.0 | 236.1 | 232.6 | LPC 20:4-SN1 | LPC(20:4)>LPC(20:4/0:0)_and_LPC(0:0/20:4) | 1.5 |

| | | | | | | | | |
|---|---|---|---|---|---|---|---|---|
| 88 | 552.4026 | 552.4025 | 0.1 | 242.6 | 246.4 | LPC 20:0-SN1 | LPC(20:0)>LPC(20:0/0:0)_and_LPC(0:0/20:0) | -1.6 |
| 89 | 802.6322 | 802.6332 | 1.3 | 303.1 | 298.3 | PC 37:1 | PC(37:1) | 1.6 |
| 90 | 714.6189 | 714.6185 | 0.6 | 291.1 | 295.9 | CE 22:6 | CE(22:6)M+NH3 | -1.6 |
| 91 | 688.6030 | 688.6026 | 0.7 | 296.3 | 291.4 | CE 20:5 | CE(20:5)M+NH3 | 1.7 |
| 92 | 704.5224 | 704.5225 | 0.2 | 282.0 | 277.2 | PC 30:1 | PC(30:1) | 1.7 |
| 93 | 800.6165 | 800.6174 | 1.1 | 300.7 | 295.5 | PC 37:2 | PC(37:2) | 1.8 |
| 94 | 673.5281 | 673.5277 | 0.7 | 271.8 | 276.9 | SM d32:2 | SM(d32:2) | -1.8 |
| 95 | 744.5540 | 744.5540 | 0.0 | 288.1 | 282.7 | PC 33:2 | PC(33:2) | 1.9 |
| 96 | 748.6220 | 748.6205 | 1.9 | 288.7 | 294.4 | PC O-34:0 | PC(O-34:0) | -1.9 |
| 97 | 733.6217 | 733.6208 | 1.3 | 289.9 | 295.7 | SM d36:0 | SM(d36:0) | -2.0 |
| 98 | 801.6846 | 801.6854 | 1.1 | 300.4 | 306.5 | SM d41:1 | SM(d41:1) | -2.0 |
| 99 | 759.6375 | 759.6373 | 0.3 | 292.3 | 298.2 | SM d38:1 | SM(d38:1) | -2.0 |
| 100 | 718.5383 | 718.5383 | 0.1 | 285.3 | 279.7 | PC 17:0_14:1 | PC(31:1) | 2.0 |
| 101 | 518.3242 | 518.3242 | 0.0 | 230.6 | 226.1 | LPC 18:3-SN1 | LPC(18:3)>LPC(18:3/0:0)_and_LPC(0:0/18:3) | 2.0 |
| 102 | 838.6318 | 838.6315 | 0.3 | 307.9 | 301.8 | PC 40:4 | PC(40:4) | 2.0 |
| 103 | 773.6533 | 773.6531 | 0.2 | 294.7 | 300.8 | SM d39:1 | SM(d39:1) | -2.0 |
| 104 | 816.6475 | 816.6468 | 0.8 | 306.9 | 300.8 | PC 38:1 | PC(38:1) | 2.0 |
| 105 | 820.6212 | 820.6209 | 0.3 | 305.1 | 298.9 | PC O-40:6 | PC(O-40:6) | 2.1 |
| 106 | 720.5904 | 720.5898 | 0.8 | 281.7 | 287.9 | PC O-32:0 | PC(O-32:0) | -2.1 |
| 107 | 717.5906 | 717.5905 | 0.2 | 283.5 | 289.8 | SM d35:1 | SM(d35:1) | -2.2 |
| 108 | 890.8170 | 890.8168 | 0.2 | 331.9 | 324.9 | TG 17:0_18:1_18:1 | TG(53:2) M + NH3 | 2.2 |
| 109 | 728.5228 | 728.5231 | 0.4 | 283.2 | 277.1 | PC 32:3 | PC(32:3) | 2.2 |
| 110 | 494.3242 | 494.3243 | 0.2 | 222.1 | 227.1 | LPC 16:1-SN1 | LPC(16:1)>LPC(16:1/0:0)_and_LPC(0:0/16:1) | -2.2 |
| 111 | 818.6056 | 818.6057 | 0.1 | 304.7 | 298 | PC O-40:7 | PC(O-40:7) | 2.3 |

| | | | | | | | | |
|---|---|---|---|---|---|---|---|---|
| 112 | 834.6008 | 834.6002 | 0.8 | 306.4 | 299.5 | PC 18:0_22:6 | PC(40:6) | 2.3 |
| 113 | 770.5694 | 770.5696 | 0.3 | 294.7 | 288 | PC 35:3 | PC(35:3) | 2.3 |
| 114 | 701.5594 | 701.5615 | 3.0 | 276.9 | 283.5 | SM d18:1_16:1 | SM(d34:2) | -2.3 |
| 115 | 661.5283 | 661.5282 | 0.2 | 271.1 | 277.7 | SM d31:1 | SM(d31:1) | -2.4 |
| 116 | 768.5538 | 768.5540 | 0.3 | 293.0 | 286.2 | PC 35:4 | PC(35:4) | 2.4 |
| 117 | 782.5693 | 782.5695 | 0.2 | 294.4 | 287.5 | PC 18:2_18:2 Cis | PC(36:4) | 2.4 |
| 118 | 808.5848 | 808.5848 | 0.0 | 301.4 | 294.2 | PC 38:5 | PC(38:5) | 2.4 |
| 119 | 756.5540 | 756.5540 | 0.1 | 290.2 | 283.3 | PC 34:3 | PC(34:3) | 2.4 |
| 120 | 780.5539 | 780.5542 | 0.3 | 294.4 | 287.3 | PC 36:5 | PC(36:5) | 2.5 |
| 121 | 824.6164 | 824.6160 | 0.5 | 307.4 | 299.7 | PC 39:4 | PC(39:4) | 2.6 |
| 122 | 568.3398 | 568.3397 | 0.2 | 242.5 | 236.4 | LPC 22:6-SN2 | LPC(22:6)>LPC(22:6/0:0)_and_LPC(0:0/22:6) | 2.6 |
| 123 | 946.8796 | 946.8787 | 1.0 | 344.5 | 335.8 | TG 18:1_18:1_21:0 | TG(57:2) M + NH3 | 2.6 |
| 124 | 836.6160 | 836.6154 | 0.7 | 307.6 | 299.8 | PC 40:5 | PC(40:5) | 2.6 |
| 125 | 647.5125 | 647.5122 | 0.5 | 267.9 | 275 | SM d18:1_12:0 | SM(d30:1) | -2.6 |
| 126 | 960.8952 | 960.8946 | 0.6 | 347.2 | 338.4 | TG 18:1_18:1_22:0 | TG(58:2) M + NH3 | 2.6 |
| 127 | 806.5694 | 806.5692 | 0.3 | 301.0 | 293.1 | PC 38:6 (PC 16:0_22:6) | PC(38:6) | 2.7 |
| 128 | 780.5903 | 780.5903 | 0.0 | 298.6 | 290.7 | PC O-37:5 | PC(O-37:5) | 2.7 |
| 129 | 689.5593 | 689.5599 | 0.8 | 276.0 | 283.8 | SM d33:1 | SM(d33:1) | -2.8 |
| 130 | 930.8481 | 930.8477 | 0.4 | 340.0 | 330.9 | TG 18:1_18:1_20:1 | TG(56:3)>TG(18:1_18:1_20:1) M + NH3 | 2.8 |
| 131 | 944.7701 | 944.7694 | 0.8 | 335.7 | 326.7 | TG 16:0_20:4_22:6 | TG(58:10) M + NH3 | 2.8 |
| 132 | 690.6187 | 690.6187 | 0.1 | 285.7 | 294 | CE 20:4 | CE(20:4)M+NH3 | -2.8 |
| 133 | 754.5383 | 754.5381 | 0.4 | 290.0 | 281.9 | PC 34:4 | PC(34:4) | 2.9 |
| 134 | 731.6067 | 731.6062 | 0.7 | 284.1 | 292.6 | SM d18:1_18:0 | SM(d36:1) | -2.9 |
| 135 | 675.5438 | 675.5447 | 1.4 | 272.3 | 280.5 | SM d32:1 | SM(d32:1) | -2.9 |

| | | | | | | | | |
|---|---|---|---|---|---|---|---|---|
| 136 | 848.6526 | 848.6528 | 0.2 | 313.1 | 304.2 | PC O-42:6 | PC(O-42:6) | 2.9 |
| 137 | 508.3400 | 508.3400 | 0.1 | 225.6 | 232.5 | LPC 17:1-SN2 | LPC(17:1)>LPC(17:1/0:0)_and_LPC(0:0/17:1) | -3.0 |
| 138 | 866.6632 | 866.6643 | 1.4 | 318.2 | 308.8 | PC 42:4 | PC(42:4) | 3.0 |
| 139 | 494.3242 | 494.3244 | 0.3 | 221.5 | 228.5 | LPC 16:1-SN2 | LPC(16:1)>LPC(16:1/0:0)_and_LPC(0:0/16:1) | -3.0 |
| 140 | 948.8956 | 948.8944 | 1.3 | 347.6 | 337.2 | TG 16:0_16:1_25:0 | TG(57:1)>TG(16:0_18:1_23:0)_and_TG(18:0_18:1_21:0) M + NH3 | 3.1 |
| 141 | 794.5695 | 794.5694 | 0.0 | 300.1 | 291.1 | PC 37:5 | PE(40:5) | 3.1 |
| 142 | 522.3556 | 522.3562 | 1.1 | 227.9 | 235.2 | LPC 18:1-SN1 | LPC(18:1)>LPC(18:1/0:0)_and_LPC(0:0/18:1) | -3.1 |
| 143 | 568.3398 | 568.3396 | 0.5 | 242.6 | 235.2 | LPC 22:6-SN1 | LPC(22:6)>LPC(22:6/0:0)_and_LPC(0:0/22:6) | 3.1 |
| 144 | 522.3555 | 522.3554 | 0.2 | 229.3 | 236.9 | LPC 18:1-SN2 | LPC(18:1)>LPC(18:1/0:0)_and_LPC(0:0/18:1) | -3.2 |
| 145 | 958.8795 | 958.8787 | 0.8 | 347.2 | 336.4 | TG 18:1_18:2_22:0 | TG(58:3) M + NH3 | 3.2 |
| 146 | 806.5695 | 806.5689 | 0.7 | 301.0 | 291.4 | PC 38:6 (PC 18:2_20:4) | PC(38:6) | 3.3 |
| 147 | 570.3554 | 570.3553 | 0.2 | 244.3 | 236.4 | LPC 22:5-SN1 | LPC(22:5)>LPC(22:5/0:0)_and_LPC(0:0/22:5) | 3.3 |
| 148 | 703.5749 | 703.5778 | 4.2 | 276.6 | 286.3 | SM d18:1_16:0 | SM(d34:1) | -3.4 |
| 149 | 482.3242 | 482.3243 | 0.4 | 220.4 | 228.3 | LPC 15:0-SN1 | LPC(15:0)>LPC(15:0/0:0)_and_LPC(0:0/15:0) | -3.5 |
| 150 | 832.5845 | 832.5846 | 0.1 | 307.5 | 296.8 | PC 40:7 | PC(40:7) | 3.6 |
| 151 | 705.5903 | 705.5891 | 1.6 | 279.5 | 290.1 | SM d34:0 | SM(d34:0) | -3.7 |
| 152 | 820.5848 | 820.5850 | 0.3 | 307.7 | 296.6 | PC 39:6 | PC(39:6) | 3.8 |
| 153 | 677.5595 | 677.5582 | 1.9 | 274.2 | 285 | SM d32:0 | SM(d32:0) | -3.8 |
| 154 | 572.3710 | 572.3710 | 0.1 | 247.8 | 238.6 | LPC 22:4-SN1 | LPC(22:4)>LPC(22:4/0:0)_and_LPC(0:0/22:4) | 3.8 |
| 155 | 864.6473 | 864.6473 | 0.0 | 318.3 | 305.5 | PC 42:5 | PC(42:5) | 4.2 |
| 156 | 830.5693 | 830.5690 | 0.4 | 307.1 | 294 | PC 20:4_20:4 Cis | PC(40:8) | 4.4 |
| 157 | 524.3710 | 524.3717 | 1.2 | 228.9 | 240.1 | LPC 18:0-SN1 | LPC(18:0)>LPC(18:0/0:0)_and_LPC(0:0/18:0) | -4.7 |
| 158 | 774.6376 | 774.6368 | 1.0 | 283.0 | 297.4 | PC O-36:1 | PC(O-36:1) | -4.8 |

| | | | | | | | | |
|---|---|---|---|---|---|---|---|---|
| 159 | 764.52271 | 764.52264 | 0.1 | 294.0 | 280 | PE 16:0_22:6 | PE(38:6) | 5.0 |
| 160 | 496.33987 | 496.34047 | 1.2 | 220.5 | 232.1 | LPC 16:0-SN1 | LPC(16:0)>LPC(16:0/0:0)_and_LPC(0:0/16:0) | -5.0 |
| 161 | 524.37122 | 524.37145 | 0.4 | 228.4 | 241 | LPC 18:0-SN2 | LPC(18:0)>LPC(18:0/0:0)_and_LPC(0:0/18:0) | -5.2 |
| 162 | 468.30847 | 468.3084 | 0.1 | 212.4 | 224.6 | LPC 14:0-SN1 | LPC(14:0)>LPC(14:0/0:0)_and_LPC(0:0/14:0) | -5.4 |
| 163 | 510.35565 | 510.35546 | 0.4 | 223.1 | 236.4 | LPC 17:0-SN1 | LPC(17:0)>LPC(17:0/0:0)_and_LPC(0:0/17:0) | -5.6 |
| 164 | 468.3085 | 468.30869 | 0.4 | 212.5 | 225.5 | LPC 14:0-SN2 | LPC(14:0)>LPC(14:0/0:0)_and_LPC(0:0/14:0) | -5.8 |
| 165 | 496.33994 | 496.34021 | 0.5 | 218.2 | 233.7 | LPC 16:0-SN2 | LPC(16:0)>LPC(16:0/0:0)_and_LPC(0:0/16:0) | -6.6 |
| 166 | 766.57428 | 766.57459 | 0.4 | 307.0 | 287.5 | PC O-36:5 (PC O-16:1_20:4) | PC(O-36:5) | 6.8 |

**Supplementary Table S8. Summary of lipid features with matched $^{Orbi}$CCS to reference $^{DT}$CCS dataset for experiments conducted at a resolution of 240,000 and turbopump speed 68%.**

| # | m/z_this study | m/z_database | mzError_ppm | $^{Orbi}$CCS_this study | $^{DT}$CCS_database | Name_database | Name_This study | % error |
|---|---|---|---|---|---|---|---|---|
| 1 | 729.5905 | 729.5910 | 0.7 | 285.4 | 285.3 | SM 36:2 | SM(d36:2) | 0.1 |
| 2 | 787.6688 | 787.6693 | 0.6 | 298.5 | 299.1 | SM 40:1 | SM(d40:1) | -0.2 |
| 3 | 538.3870 | 538.3872 | 0.4 | 240.0 | 240.5 | LPC 19:0 | LPC(19:0)>LPC(19:0/0:0)_and_LPC(0:0/19:0) | -0.2 |
| 4 | 786.6007 | 786.6012 | 0.7 | 294.2 | 293.3 | PC (18:1/18:1) (del9-trans) | PC(36:2) | 0.3 |
| 5 | 799.6690 | 799.6693 | 0.4 | 301.1 | 300.1 | SM 41:2 | SM(d41:2) | 0.3 |
| 6 | 745.6220 | 745.6223 | 0.5 | 290.8 | 289.8 | SM 37:1 | SM(d37:1) | 0.4 |
| 7 | 759.6375 | 759.6380 | 0.6 | 292.3 | 293.4 | SM 38:1 | SM(d38:1) | -0.4 |
| 8 | 785.6533 | 785.6536 | 0.4 | 298.5 | 296.8 | SM 40:2 | SM(d40:2) | 0.6 |
| 9 | 801.6846 | 801.6849 | 0.4 | 300.4 | 302.3 | SM 41:1 | SM(d41:1) | -0.6 |
| 10 | 827.7001 | 827.7006 | 0.6 | 307.8 | 305.7 | SM 43:2 | SM(d43:2) | 0.7 |
| 11 | 773.6534 | 773.6536 | 0.3 | 294.9 | 297 | SM 39:1 | SM(d39:1) | -0.7 |
| 12 | 813.6845 | 813.6849 | 0.6 | 304.5 | 302.2 | SM 42:2 | SM(d42:2) | 0.8 |
| 13 | 811.6687 | 811.6693 | 0.7 | 304.6 | 300.8 | SM 42:3 | SM(d42:3) | 1.3 |
| 14 | 731.6067 | 731.6067 | 0.0 | 284.1 | 288.4 | SM 36:1 | SM(d36:1) | -1.5 |
| 15 | 703.5749 | 703.5754 | 0.7 | 276.6 | 281.2 | SM 34:1 | SM(d34:1) | -1.6 |
| 16 | 522.3555 | 522.3559 | 0.8 | 229.3 | 233.2 | LPC 18:1 | LPC(18:1)>LPC(18:1/0:0)_and_LPC(0:0/18:1) | -1.7 |
| 17 | 788.6165 | 788.6169 | 0.5 | 294.3 | 289.4 | PC 36:1 | PC(36:1) | 1.7 |
| 18 | 760.5850 | 760.5856 | 0.8 | 292.9 | 287.9 | PC (18:1(9Z)/16:0) | PC(34:1) | 1.7 |
| 19 | 730.5385 | 730.5387 | 0.3 | 283.4 | 278.4 | PC (16:1/16:1) (del9-cis) | PC(32:2) | 1.8 |
| 20 | 552.4026 | 552.4029 | 0.6 | 242.6 | 247 | 20:0 Lyso PC | LPC(20:0)>LPC(20:0/0:0)_and_LPC(0:0/20:0) | -1.8 |
| 21 | 732.5539 | 732.5543 | 0.6 | 283.4 | 277.6 | PC 32:1 | PC(32:1) | 2.1 |
| 22 | 774.6009 | 774.6013 | 0.5 | 292.8 | 285.8 | PC 35:1 | PC(35:1) | 2.5 |
| 23 | 786.6007 | 786.6012 | 0.7 | 294.2 | 287.1 | PC 36:2 | PC(36:2) | 2.5 |
| 24 | 758.5693 | 758.5699 | 0.8 | 290.7 | 280.6 | PC 34:2 | PC(34:2) | 3.6 |
| 25 | 468.3085 | 468.3090 | 1.1 | 212.5 | 220.8 | LPC 14:0 | LPC(14:0)>LPC(14:0/0:0)_and_LPC(0:0/14:0) | -3.8 |
| 26 | 756.5540 | 756.5543 | 0.4 | 290.2 | 278.2 | PC 34:3 | PC(34:3) | 4.3 |
| 27 | 524.3712 | 524.3716 | 0.7 | 228.4 | 238.8 | 18:0 Lyso PC | LPC(18:0)>LPC(18:0/0:0)_and_LPC(0:0/18:0) | -4.3 |
| 28 | 728.5593 | 728.5594 | 0.2 | 286.5 | 273.5 | PE (O-36:3) | PE(O-36:3)>PE(O-18:1/18:2) | 4.8 |
| 29 | 496.3399 | 496.3403 | 0.7 | 218.2 | 231.4 | 16:0 Lyso PC | LPC(16:0)>LPC(16:0/0:0)_and_LPC(0:0/16:0) | -5.7 |
| 30 | 400.3421 | 400.3427 | 1.5 | 187.2 | 214.7 | Carnitine 16:0 | Car(16:0) | -12.8 |

| | | | | | | | | |
|---|---|---|---|---|---|---|---|---|
| 31 | 344.2795 | 344.2782 | 3.7 | 172.1 | 199.5 | Carnitine 12:0 | Car(12:0) | -13.7 |

**Supplementary Table S9. Summary of lipid features with matched $^{Orbi}$CCS to reference $^{TIMS}$CCS dataset for experiments conducted at resolution of 180,000 and turbopump speed 68%.**

| # | m/z_this study | m/z_database | mzError_ppm | $^{Orbi}$CCS_this study | $^{TIMS}$CCS_database | Name_database | Name_This study | % error |
|---|---|---|---|---|---|---|---|---|
| 1 | 520.3399 | 520.3402 | 0.6 | 231.2 | 231.2 | LPC 18:2-SN2 | LPC(18:2)>LPC(18:2/0:0)_and_LPC(0:0/18:2) | 0.0 |
| 2 | 848.7701 | 848.7708 | 0.9 | 315.3 | 315.3 | TG 16:0_16:1_18:1 | TG(50:2)>TG(16:0_16:1_18:1)_and_TG(14:0_18:1_18:1) M + NH3 | 0.0 |
| 3 | 746.5695 | 746.5697 | 0.3 | 286.8 | 286.7 | PC 33:1 | PC(33:1) | 0.0 |
| 4 | 813.6845 | 813.6862 | 2.1 | 306.8 | 306.6 | SM d18:1_24:1 | SM(d42:2) | 0.1 |
| 5 | 827.7000 | 827.6999 | 0.1 | 310.4 | 310.1 | SM d43:2 | SM(d43:2) | 0.1 |
| 6 | 820.7390 | 820.7395 | 0.5 | 309.0 | 309.3 | TG 14:0_16:1_18:1 | TG(48:2)>TG(14:0_16:0_18:2)_and_TG(14:0_16:1_18:1) M + NH3 | -0.1 |
| 7 | 716.5588 | 716.5588 | 0.1 | 282.7 | 282.4 | PC O-32:2 | PC(O-32:2) | 0.1 |
| 8 | 732.5540 | 732.5543 | 0.4 | 283.7 | 283.3 | PC 16:0_16:1 | PC(32:1) | 0.1 |
| 9 | 874.7858 | 874.7875 | 1.9 | 320.5 | 320 | TG 16:0_18:1_18:2 | TG(52:3)>TG(16:0_18:1_18:2)_and_TG(16:1_18:1_18:1) M + NH3 | 0.2 |
| 10 | 832.7389 | 832.7390 | 0.1 | 311.5 | 310.9 | TG 15:0_16:1_18:2 | TG(49:3) M + NH3 | 0.2 |
| 11 | 744.5903 | 744.5890 | 1.7 | 288.1 | 287.6 | PC O-34:2 | PC(O-34:2) | 0.2 |
| 12 | 762.6009 | 762.5990 | 2.5 | 293.5 | 292.8 | PC 18:0_16:0 | PC(34:0) | 0.2 |
| 13 | 799.6690 | 799.6687 | 0.3 | 302.0 | 302.8 | SM d41:2 | SM(d41:2) | -0.3 |
| 14 | 727.5753 | 727.5748 | 0.7 | 286.0 | 286.8 | SM d36:3 | SM(d36:3) | -0.3 |
| 15 | 792.7076 | 792.7083 | 0.8 | 304.2 | 303.3 | TG 12:0_16:1_18:1 | TG(46:2) M + NH3 | 0.3 |
| 16 | 768.5903 | 768.5895 | 1.0 | 292.6 | 291.7 | PC O-36:4 | PC(O-36:4) | 0.3 |
| 17 | 785.6533 | 785.6531 | 0.2 | 300.3 | 301.2 | SM d40:2 | SM(d40:2) | -0.3 |
| 18 | 794.7233 | 794.7233 | 0.0 | 304.4 | 305.5 | TG 14:0_16:0_16:1 | TG(46:1)>TG(12:0_16:0_18:1)_and_TG(10:0_18:0_18:1)_and_TG(14:0_16:0_16:1) M + NH3 | -0.3 |
| 19 | 774.6008 | 774.6011 | 0.3 | 293.4 | 292.4 | PC 35:1 | PC(35:1) | 0.4 |
| 20 | 850.7858 | 850.7857 | 0.1 | 316.8 | 318 | TG 16:0_16:0_18:1 | TG(50:1)>TG(16:0_16:0_18:1)_M+NH3 | -0.4 |
| 21 | 818.7233 | 818.7238 | 0.6 | 308.6 | 307.4 | TG 16:1_16:1_16:1 | TG(48:3) M + NH3 | 0.4 |

| | | | | | | | | |
|---|---|---|---|---|---|---|---|---|
| 22 | 846.7545 | 846.7556 | 1.4 | 314.8 | 313.5 | TG 16:1_16:0_18:2 | TG(50:3) M + NH3 | 0.4 |
| 23 | 834.7545 | 834.7550 | 0.6 | 310.8 | 312.2 | TG 15:0_16:1_18:1 | TG(49:2) M + NH3 | -0.4 |
| 24 | 715.5750 | 715.5748 | 0.2 | 285.3 | 286.6 | SM d35:2 | SM(d35:2) | -0.5 |
| 25 | 836.7703 | 836.7703 | 0.1 | 316.1 | 314.6 | TG 15:0_16:0_18:1 | TG(49:1)>TG(15:0_16:0_18:0)_and_TG(16:0_16:0_17:1)_and_TG(16:0_16:1_17:0) M + NH3 | 0.5 |
| 26 | 766.6921 | 766.6927 | 0.8 | 298.1 | 299.8 | TG 12:0_16:0_16:1 | TG(44:1)>TG(10:0_16:0_18:1) M + NH3 | -0.6 |
| 27 | 788.6165 | 788.6167 | 0.3 | 297.1 | 295.4 | PC 18:0_18:1 | PC(36:1) | 0.6 |
| 28 | 536.3713 | 536.3720 | 1.3 | 239.2 | 237.8 | LPC 19:1-SN1 | LPC(19:1)>LPC(19:1/0:0)_and_LPC(0:0/19:1) | 0.6 |
| 29 | 796.7390 | 796.7391 | 0.2 | 305.9 | 307.8 | TG 14:0_16:0_16:0 | TG(46:0)>TG(14:0_16:0_16:0)_and_TG(12:0_16:0_18:0) M + NH3 | -0.6 |
| 30 | 796.6216 | 796.6210 | 0.9 | 299.4 | 297.5 | PC O-38:4 | PC(O-38:4) | 0.6 |
| 31 | 862.7858 | 862.7861 | 0.4 | 320.2 | 318.2 | TG 16:0_17:1_18:1 | TG(51:2)>TG(15:0_18:1_18:1)_and_TG(16:0_17:1_18:1)_and_TG(16:1_17:0_18:1) M + NH3 | 0.6 |
| 32 | 794.6057 | 794.6055 | 0.3 | 297.6 | 295.7 | PC O-38:5 | PC(O-38:5) | 0.6 |
| 33 | 772.6217 | 772.6210 | 0.9 | 296.3 | 294.3 | PC O-36:2 | PC(O-36:2) | 0.7 |
| 34 | 872.7701 | 872.7720 | 2.1 | 320.1 | 317.7 | TG 16:0_18:2_18:2 | TG(52:4)>TG(16:0_16:0_20:4)_and_TG(16:1_18:1_18:2)_and_TG(16:0_18:1_18:2) M + NH3 | 0.7 |
| 35 | 706.5378 | 706.5382 | 0.5 | 278.4 | 280.5 | PC 30:0 | PC(30:0) | -0.8 |
| 36 | 830.7236 | 830.7228 | 1.0 | 311.8 | 309.4 | TG 15:1_16:1_18:2 | TG(49:4) M + NH3 | 0.8 |
| 37 | 546.3555 | 546.3547 | 1.5 | 237.3 | 235.4 | LPC 20:3-SN2 | LPC(20:3)>LPC(20:3/0:0)_and_LPC(0:0/20:3) | 0.8 |
| 38 | 678.5070 | 678.5068 | 0.3 | 273.0 | 275.2 | PC 14:0_14:0 | PC(28:0) | -0.8 |
| 39 | 770.6059 | 770.6059 | 0.1 | 294.2 | 291.8 | PC O-36:3 | PC(O-36:3) | 0.8 |
| 40 | 717.5905 | 717.5905 | 0.0 | 287.4 | 289.8 | SM d35:1 | SM(d35:1) | -0.8 |
| 41 | 550.3867 | 550.3868 | 0.1 | 240.3 | 242.4 | LPC 20:1-SN1 | LPC(20:1)>LPC(20:1/0:0)_and_LPC(0:0/20:1) | -0.9 |
| 42 | 822.7546 | 822.7547 | 0.1 | 308.8 | 311.6 | TG 14:0_16:0_18:1 | TG(48:1)>TG(16:0_16:0_16:1)_and_TG(14:0_16:0_18:1) M + NH3 | -0.9 |
| 43 | 757.6218 | 757.6216 | 0.3 | 292.9 | 295.6 | SM d38:2 | SM(d38:2) | -0.9 |

| | | | | | | | | |
|---|---|---|---|---|---|---|---|---|
| 44 | 786.6014 | 786.6026 | 1.5 | 295.5 | 292.7 | PC 18:0_18:2 | PC(36:2) | 1.0 |
| 45 | 746.6062 | 746.6054 | 1.1 | 288.2 | 291 | PC O-34:1 | PC(O-34:1) | -1.0 |
| 46 | 811.6687 | 811.6685 | 0.2 | 306.9 | 303.9 | SM d42:3 | SM(d42:3) | 1.0 |
| 47 | 772.5851 | 772.5857 | 0.8 | 292.5 | 289.4 | PC 35:2 | PC(35:2) | 1.1 |
| 48 | 844.7387 | 844.7396 | 1.0 | 315.0 | 311.6 | TG 14:0_18:2_18:2 | TG(50:4)>TG(16:1_16:1_18:2)_and_TG(16:1_16:1_18:2) M + NH3 | 1.1 |
| 49 | 839.6997 | 839.6997 | 0.1 | 314.0 | 310.6 | SM d44:3 | SM(d44:3) | 1.1 |
| 50 | 784.5851 | 784.5867 | 2.1 | 294.4 | 291.1 | PC 18:1_18:2 | PC(36:3) | 1.1 |
| 51 | 787.6690 | 787.6697 | 1.0 | 299.5 | 303 | SM d40:1 | SM(d40:1) | -1.1 |
| 52 | 580.4338 | 580.4346 | 1.3 | 256.9 | 253.9 | LPC 22:0-SN1 | LPC(22:0)>LPC(22:0/0:0)_and_LPC(0:0/22:0) | 1.2 |
| 53 | 825.6843 | 825.6846 | 0.3 | 309.7 | 306 | SM d43:3 | SM(d43:3) | 1.2 |
| 54 | 902.8169 | 902.8167 | 0.2 | 329.7 | 325.8 | TG 18:1_18:1_18:1 | TG(54:3)>TG(18:0_18:1_18:2)_and_TG(18:1/18:1/18:1)_and_TG(16:0_18:2_20:1) M + NH3 | 1.2 |
| 55 | 548.3712 | 548.3708 | 0.8 | 239.5 | 236.5 | LPC 20:2-SN1 | LPC(20:2)>LPC(20:2/0:0)_and_LPC(0:0/20:2) | 1.2 |
| 56 | 870.7545 | 870.7554 | 1.1 | 320.2 | 316.2 | TG 16:1_18:2_18:2 | TG(52:5)>TG(16:0_18:2_18:3) M + NH3 | 1.3 |
| 57 | 760.5851 | 760.5870 | 2.5 | 293.4 | 289.7 | PC 16:0_18:1 | PC(34:1) | 1.3 |
| 58 | 688.6029 | 688.6026 | 0.5 | 295.1 | 291.4 | CE 20:5 | CE(20:5)M+NH3 | 1.3 |
| 59 | 734.5696 | 734.5696 | 0.0 | 282.9 | 286.6 | PC 16:0_16:0 | PC(32:0) | -1.3 |
| 60 | 718.5747 | 718.5748 | 0.1 | 281.4 | 285.2 | PC O-32:1 | PC(O-32:1) | -1.3 |
| 61 | 810.6010 | 810.6011 | 0.2 | 300.6 | 296.7 | PC 18:0_20:4 | PC(38:4) | 1.3 |
| 62 | 774.6373 | 774.6368 | 0.7 | 293.3 | 297.4 | PC O-36:1 | PC(O-36:1) | -1.4 |
| 63 | 758.5694 | 758.5718 | 3.2 | 290.4 | 286.4 | PC 16:0_18:2 | PC(34:2) | 1.4 |
| 64 | 721.5858 | 721.5842 | 2.2 | 286.3 | 290.4 | SM 34:0;3O | SM(d34:0-OH) | -1.4 |
| 65 | 759.6376 | 759.6373 | 0.4 | 293.9 | 298.2 | SM d38:1 | SM(d38:1) | -1.4 |
| 66 | 714.6188 | 714.6185 | 0.4 | 291.4 | 295.9 | CE 22:6 | CE(22:6)M+NH3 | -1.5 |

| | | | | | | | | |
|---|---|---|---|---|---|---|---|---|
| 67 | 729.5907 | 729.5904 | 0.4 | 285.3 | 289.7 | SM d18:1_18:1 | SM(d36:2) | -1.5 |
| 68 | 730.5385 | 730.5386 | 0.2 | 283.9 | 279.6 | PC 32:2 | PC(32:2) | 1.5 |
| 69 | 809.6527 | 809.6523 | 0.5 | 306.3 | 301.5 | SM d42:4 | SM(d40:1) M + Na | 1.6 |
| 70 | 858.7543 | 858.7544 | 0.1 | 320.0 | 314.9 | TG 15:0_18:2_18:2 | TG(51:4)>TG(16:1_17:1_18:2)_and_TG(15:0_18:2_18:2)_and_TG(15:1_18:1_18:2) M + NH3 | 1.6 |
| 71 | 801.6847 | 801.6854 | 0.9 | 301.5 | 306.5 | SM d41:1 | SM(d41:1) | -1.6 |
| 72 | 546.3555 | 546.3552 | 0.5 | 237.7 | 233.8 | LPC 20:3-SN1 | LPC(20:3)>LPC(20:3/0:0)_and_LPC(0:0/20:3) | 1.7 |
| 73 | 704.5225 | 704.5225 | 0.1 | 281.9 | 277.2 | PC 30:1 | PC(30:1) | 1.7 |
| 74 | 494.3242 | 494.3243 | 0.3 | 223.3 | 227.1 | LPC 16:1-SN1 | LPC(16:1)>LPC(16:1/0:0)_and_LPC(0:0/16:1) | -1.7 |
| 75 | 748.6220 | 748.6205 | 1.9 | 289.4 | 294.4 | PC O-34:0 | PC(O-34:0) | -1.7 |
| 76 | 868.7387 | 868.7387 | 0.0 | 320.2 | 314.8 | TG 16:1_18:2_18:3 | TG(52:6) M + NH3 | 1.7 |
| 77 | 773.6532 | 773.6531 | 0.1 | 295.7 | 300.8 | SM d39:1 | SM(d39:1) | -1.7 |
| 78 | 538.3869 | 538.3872 | 0.5 | 238.8 | 243.2 | LPC 19:0-SN2 | LPC(19:0)>LPC(19:0/0:0)_and_LPC(0:0/19:0) | -1.8 |
| 79 | 782.5694 | 782.5701 | 0.8 | 295.8 | 290.4 | PC 16:0_20:4 | PC(36:4) | 1.9 |
| 80 | 900.8011 | 900.8010 | 0.1 | 329.9 | 323.8 | TG 18:1_18:1_18:2 | TG(54:4)>TG(18:1_18:1_18:2) M + NH3 | 1.9 |
| 81 | 522.3552 | 522.3562 | 1.8 | 230.7 | 235.2 | LPC 18:1-SN1 | LPC(18:1)>LPC(18:1/0:0)_and_LPC(0:0/18:1) | -1.9 |
| 82 | 816.6474 | 816.6468 | 0.8 | 306.8 | 300.8 | PC 38:1 | PC(38:1) | 2.0 |
| 83 | 888.8012 | 888.8012 | 0.0 | 329.5 | 323.1 | TG 17:0_18:1_18:2 | TG(53:3)>TG(17:0_18:1_18:2)_and_TG(17:1_18:1_18:1)_and_TG(16:1_18:1_19:1)_and_TG(16:0_18:1_19:2) M + NH3 | 2.0 |
| 84 | 552.4027 | 552.4025 | 0.3 | 241.3 | 246.4 | LPC 20:0-SN1 | LPC(20:0)>LPC(20:0/0:0)_and_LPC(0:0/20:0) | -2.1 |
| 85 | 812.6165 | 812.6163 | 0.2 | 303.3 | 297 | PC 38:3 | PC(38:3) | 2.1 |
| 86 | 544.3399 | 544.3399 | 0.1 | 237.6 | 232.6 | LPC 20:4-SN1 | LPC(20:4)>LPC(20:4/0:0)_and_LPC(0:0/20:4) | 2.2 |
| 87 | 814.6320 | 814.6311 | 1.1 | 304.9 | 298.4 | PC 38:2 | PC(38:2) | 2.2 |
| 88 | 673.5281 | 673.5277 | 0.6 | 270.9 | 276.9 | SM d32:2 | SM(d32:2) | -2.2 |

| | | | | | | | | |
|---|---|---|---|---|---|---|---|---|
| 89 | 720.5905 | 720.5898 | 0.9 | 281.6 | 287.9 | PC O-32:0 | PC(O-32:0) | -2.2 |
| 90 | 904.8326 | 904.8316 | 1.1 | 334.8 | 327.7 | TG 18:0_18:1_18:1 | TG(54:2)>TG(16:0_18:1_20:1)_and_TG(18:0_18:1_18:1) M + NH3 | 2.2 |
| 91 | 482.3242 | 482.3243 | 0.2 | 223.3 | 228.3 | LPC 15:0-SN1 | LPC(15:0)>LPC(15:0/0:0)_and_LPC(0:0/15:0) | -2.2 |
| 92 | 890.8170 | 890.8168 | 0.2 | 332.1 | 324.9 | TG 17:0_18:1_18:1 | TG(53:2) M + NH3 | 2.2 |
| 93 | 770.5696 | 770.5696 | 0.0 | 294.7 | 288 | PC 35:3 | PC(35:3) | 2.3 |
| 94 | 898.7857 | 898.7855 | 0.2 | 329.5 | 322 | TG 18:1_18:2_18:2 | TG(54:5)>TG(18:1_18:2_18:2)_and_TG(16:0_16:0_22:5)_and_TG(16:0_18:1_20:4) M + NH3 | 2.3 |
| 95 | 802.6323 | 802.6332 | 1.2 | 305.3 | 298.3 | PC 37:1 | PC(37:1) | 2.3 |
| 96 | 701.5595 | 701.5615 | 2.9 | 276.8 | 283.5 | SM d18:1_16:1 | SM(d34:2) | -2.4 |
| 97 | 886.7856 | 886.7852 | 0.5 | 328.8 | 321.1 | TG 17:1_18:1_18:2 | TG(53:4)>TG(17:0_18:2_18:2)_and_TG(17:1_18:1_18:2) M + NH3 | 2.4 |
| 98 | 718.5383 | 718.5383 | 0.0 | 286.4 | 279.7 | PC 17:0_14:1 | PC(31:1) | 2.4 |
| 99 | 518.3242 | 518.3242 | 0.0 | 231.6 | 226.1 | LPC 18:3-SN1 | LPC(18:3)>LPC(18:3/0:0)_and_LPC(0:0/18:3) | 2.4 |
| 100 | 808.5848 | 808.5848 | 0.1 | 301.4 | 294.2 | PC 38:5 | PC(38:5) | 2.4 |
| 101 | 728.5228 | 728.5231 | 0.4 | 283.9 | 277.1 | PC 32:3 | PC(32:3) | 2.4 |
| 102 | 782.5694 | 782.5695 | 0.2 | 295.1 | 287.5 | PC 18:2_18:2 Cis | PC(36:4) | 2.6 |
| 103 | 834.6007 | 834.6002 | 0.6 | 307.4 | 299.5 | PC 18:0_22:6 | PC(40:6) | 2.6 |
| 104 | 733.6217 | 733.6208 | 1.3 | 287.8 | 295.7 | SM d36:0 | SM(d36:0) | -2.7 |
| 105 | 838.6318 | 838.6315 | 0.3 | 309.9 | 301.8 | PC 40:4 | PC(40:4) | 2.7 |
| 106 | 768.5538 | 768.5540 | 0.3 | 294.1 | 286.2 | PC 35:4 | PC(35:4) | 2.8 |
| 107 | 756.5540 | 756.5540 | 0.1 | 291.2 | 283.3 | PC 34:3 | PC(34:3) | 2.8 |
| 108 | 522.3555 | 522.3554 | 0.2 | 230.3 | 236.9 | LPC 18:1-SN2 | LPC(18:1)>LPC(18:1/0:0)_and_LPC(0:0/18:1) | -2.8 |
| 109 | 960.8951 | 960.8946 | 0.5 | 347.9 | 338.4 | TG 18:1_18:1_22:0 | TG(58:2) M + NH3 | 2.8 |
| 110 | 731.6065 | 731.6062 | 0.4 | 284.3 | 292.6 | SM d18:1_18:0 | SM(d36:1) | -2.8 |
| 111 | 824.6524 | 824.6529 | 0.7 | 311.4 | 302.8 | PC O-40:4 | PC(O-40:4) | 2.8 |

| | | | | | | | | |
|---|---|---|---|---|---|---|---|---|
| 112 | 896.7700 | 896.7699 | 0.0 | 329.5 | 320.4 | TG 18:2_18:2_18:2 | TG(54:6)>TG(16:0_18:2_20:4) M + NH3 | 2.8 |
| 113 | 661.5282 | 661.5282 | 0.0 | 269.8 | 277.7 | SM d31:1 | SM(d31:1) | -2.9 |
| 114 | 745.6220 | 745.6221 | 0.2 | 286.7 | 295.2 | SM d37:1 | SM(d37:1) | -2.9 |
| 115 | 780.5539 | 780.5542 | 0.3 | 295.7 | 287.3 | PC 36:5 | PC(36:5) | 2.9 |
| 116 | 508.3400 | 508.3400 | 0.1 | 225.7 | 232.5 | LPC 17:1-SN2 | LPC(17:1)>LPC(17:1/0:0)_and_LPC(0:0/17:1) | -2.9 |
| 117 | 884.7694 | 884.7688 | 0.7 | 328.9 | 319.5 | TG 17:1_18:2_18:2 | TG(53:5)>TG(17:1_18:2_18:2) M + NH3 | 2.9 |
| 118 | 689.5593 | 689.5599 | 0.8 | 275.2 | 283.8 | SM d33:1 | SM(d33:1) | -3.0 |
| 119 | 806.5695 | 806.5692 | 0.3 | 302.1 | 293.1 | PC 38:6 (PC 16:0_22:6) | PC(38:6) | 3.1 |
| 120 | 568.3398 | 568.3397 | 0.2 | 243.7 | 236.4 | LPC 22:6-SN2 | LPC(22:6)>LPC(22:6/0:0)_and_LPC(0:0/22:6) | 3.1 |
| 121 | 754.5383 | 754.5381 | 0.3 | 290.7 | 281.9 | PC 34:4 | PC(34:4) | 3.1 |
| 122 | 894.7543 | 894.7539 | 0.4 | 328.5 | 318.5 | TG 18:2_18:2_18:3 | TG(54:7)>TG(16:0_16:1_22:6)_and_TG(16:0_18:2_20:5) M + NH3 | 3.1 |
| 123 | 864.8010 | 864.8019 | 1.0 | 331.3 | 320.8 | TG 16:0_17:0_18:1 | TG(51:1) M + NH3 | 3.3 |
| 124 | 572.3710 | 572.3710 | 0.1 | 246.5 | 238.6 | LPC 22:4-SN1 | LPC(22:4)>LPC(22:4/0:0)_and_LPC(0:0/22:4) | 3.3 |
| 125 | 958.8795 | 958.8787 | 0.8 | 347.6 | 336.4 | TG 18:1_18:2_22:0 | TG(58:3) M + NH3 | 3.3 |
| 126 | 703.5750 | 703.5778 | 4.1 | 276.7 | 286.3 | SM d18:1_16:0 | SM(d34:1) | -3.4 |
| 127 | 675.5438 | 675.5447 | 1.4 | 271.0 | 280.5 | SM d32:1 | SM(d32:1) | -3.4 |
| 128 | 647.5126 | 647.5122 | 0.6 | 265.6 | 275 | SM d18:1_12:0 | SM(d30:1) | -3.4 |
| 129 | 946.8795 | 946.8787 | 0.9 | 347.3 | 335.8 | TG 18:1_18:1_21:0 | TG(57:2) M + NH3 | 3.4 |
| 130 | 820.6212 | 820.6209 | 0.3 | 309.2 | 298.9 | PC O-40:6 | PC(O-40:6) | 3.4 |
| 131 | 690.6186 | 690.6187 | 0.1 | 283.8 | 294 | CE 20:4 | CE(20:4)M+NH3 | -3.5 |
| 132 | 824.6163 | 824.6160 | 0.5 | 310.2 | 299.7 | PC 39:4 | PC(39:4) | 3.5 |
| 133 | 818.6055 | 818.6057 | 0.2 | 308.5 | 298 | PC O-40:7 | PC(O-40:7) | 3.5 |
| 134 | 834.6009 | 834.5999 | 1.2 | 307.8 | 297.3 | PC 40:6 | PC(40:6) | 3.5 |
| 135 | 570.3554 | 570.3553 | 0.1 | 244.9 | 236.4 | LPC 22:5-SN1 | LPC(22:5)>LPC(22:5/0:0)_and_LPC(0:0/22:5) | 3.6 |

| | | | | | | | | |
|---|---|---|---|---|---|---|---|---|
| 136 | 705.5904 | 705.5891 | 1.8 | 279.7 | 290.1 | SM d34:0 | SM(d34:0) | -3.6 |
| 137 | 806.5695 | 806.5689 | 0.7 | 302.0 | 291.4 | PC 38:6 (PC 18:2_20:4) | PC(38:6) | 3.6 |
| 138 | 568.3399 | 568.3396 | 0.6 | 243.9 | 235.2 | LPC 22:6-SN1 | LPC(22:6)>LPC(22:6/0:0)_and_LPC(0:0/22:6) | 3.7 |
| 139 | 740.5231 | 740.5227 | 0.6 | 288.4 | 278 | PE 16:0_20:4 | PE(36:4) | 3.7 |
| 140 | 780.5904 | 780.5903 | 0.1 | 301.8 | 290.7 | PC O-37:5 | PC(O-37:5) | 3.8 |
| 141 | 677.5595 | 677.5582 | 1.9 | 273.4 | 285 | SM d32:0 | SM(d32:0) | -4.1 |
| 142 | 832.5847 | 832.5846 | 0.1 | 308.9 | 296.8 | PC 40:7 | PC(40:7) | 4.1 |
| 143 | 524.3710 | 524.3717 | 1.2 | 229.8 | 240.1 | LPC 18:0-SN1 | LPC(18:0)>LPC(18:0/0:0)_and_LPC(0:0/18:0) | -4.3 |
| 144 | 510.3556 | 510.3555 | 0.3 | 225.9 | 236.4 | LPC 17:0-SN1 | LPC(17:0)>LPC(17:0/0:0)_and_LPC(0:0/17:0) | -4.4 |
| 145 | 932.8639 | 932.8642 | 0.3 | 348.2 | 333.3 | TG 18:1_18:1_20:0 | TG(56:2) M + NH3 | 4.5 |
| 146 | 924.8012 | 924.8006 | 0.6 | 342.8 | 327.6 | TG 18:1_18:1_20:4 | TG(56:6)>TG(18:1_18:1_20:4)_and_TG(16:0_18:1_22:5)_and_TG(18:1_18:2_20:3) M + NH3 | 4.6 |
| 147 | 876.8016 | 876.8023 | 0.8 | 336.8 | 321.9 | TG 16:0_18:1_18:1 | TG(52:2)>TG(16:0_18:1_18:1) M + NH3 | 4.6 |
| 148 | 866.6629 | 866.6643 | 1.6 | 323.1 | 308.8 | PC 42:4 | PC(42:4) | 4.6 |
| 149 | 468.3085 | 468.3084 | 0.1 | 213.9 | 224.6 | LPC 14:0-SN1 | LPC(14:0)>LPC(14:0/0:0)_and_LPC(0:0/14:0) | -4.8 |
| 150 | 820.5847 | 820.5850 | 0.4 | 311.0 | 296.6 | PC 39:6 | PC(39:6) | 4.8 |
| 151 | 524.3712 | 524.3715 | 0.4 | 229.2 | 241 | LPC 18:0-SN2 | LPC(18:0)>LPC(18:0/0:0)_and_LPC(0:0/18:0) | -4.9 |
| 152 | 948.8954 | 948.8944 | 1.1 | 354.0 | 337.2 | TG 16:0_16:1_25:0 | TG(57:1)>TG(16:0_18:1_23:0)_and_TG(18:0_18:1_21:0) M + NH3 | 5.0 |
| 153 | 496.3399 | 496.3402 | 0.7 | 221.8 | 233.7 | LPC 16:0-SN2 | LPC(16:0)>LPC(16:0/0:0)_and_LPC(0:0/16:0) | -5.1 |
| 154 | 468.3085 | 468.3087 | 0.4 | 214.0 | 225.5 | LPC 14:0-SN2 | LPC(14:0)>LPC(14:0/0:0)_and_LPC(0:0/14:0) | -5.1 |
| 155 | 830.5693 | 830.5690 | 0.4 | 309.1 | 294 | PC 20:4_20:4 Cis | PC(40:8) | 5.1 |
| 156 | 944.7700 | 944.7694 | 0.7 | 344.2 | 326.7 | TG 16:0_20:4_22:6 | TG(58:10) M + NH3 | 5.4 |
| 157 | 764.5227 | 764.5226 | 0.1 | 295.3 | 280 | PE 16:0_22:6 | PE(38:6) | 5.5 |
| 158 | 496.3399 | 496.3405 | 1.2 | 219.3 | 232.1 | LPC 16:0-SN1 | LPC(16:0)>LPC(16:0/0:0)_and_LPC(0:0/16:0) | -5.5 |

| | | | | | | | | |
|---|---|---|---|---|---|---|---|---|
| 159 | 864.6476 | 864.6473 | 0.3 | 322.8 | 305.5 | PC 42:5 | PC(42:5) | 5.7 |
| 160 | 766.5742 | 766.5746 | 0.5 | 307.2 | 287.5 | PC O-36:5 (PC O-16:1_20:4) | PC(O-36:5) | 6.8 |
| 161 | 856.5850 | 856.5846 | 0.4 | 319.1 | 297.6 | PC 42:9 | PC(42:9) | 7.2 |

**Supplementary Table S10. Summary of lipid features with matched $^{Orbi}$CCS to reference $^{DT}$CCS dataset for experiments conducted at resolution of 180,000 and turbopump speed 68%.**

| # | m/z_this study | m/z_database | mzError_ppm | $^{Orbi}$CCS_this study | $^{DT}$CCS_database | Name_database | Name_This study | % error |
|---|---|---|---|---|---|---|---|---|
| 1 | 729.5907 | 729.5910 | 0.4 | 285.3 | 285.3 | SM 36:2 | SM(d36:2) | 0.0 |
| 2 | 787.6690 | 787.6693 | 0.4 | 299.5 | 299.1 | SM 40:1 | SM(d40:1) | 0.2 |
| 3 | 759.6376 | 759.6380 | 0.6 | 293.9 | 293.4 | SM 38:1 | SM(d38:1) | 0.2 |
| 4 | 801.6847 | 801.6849 | 0.3 | 301.5 | 302.3 | SM 41:1 | SM(d41:1) | -0.3 |
| 5 | 773.6534 | 773.6536 | 0.3 | 296.0 | 297 | SM 39:1 | SM(d39:1) | -0.3 |
| 6 | 799.6690 | 799.6693 | 0.4 | 302.0 | 300.1 | SM 41:2 | SM(d41:2) | 0.6 |
| 7 | 538.3869 | 538.3872 | 0.6 | 238.8 | 240.5 | LPC 19:0 | LPC(19:0)>LPC(19:0/0:0)_and_LPC(0:0/19:0) | -0.7 |
| 8 | 786.6014 | 786.6012 | 0.3 | 295.5 | 293.3 | PC (18:1/18:1) (del9-trans) | PC(36:2) | 0.7 |
| 9 | 785.6533 | 785.6536 | 0.4 | 299.9 | 296.8 | SM 40:2 | SM(d40:2) | 1.0 |
| 10 | 745.6220 | 745.6223 | 0.5 | 286.7 | 289.8 | SM 37:1 | SM(d37:1) | -1.1 |
| 11 | 522.3552 | 522.3559 | 1.3 | 230.7 | 233.2 | LPC 18:1 | LPC(18:1)>LPC(18:1/0:0)_and_LPC(0:0/18:1) | -1.1 |
| 12 | 827.7001 | 827.7006 | 0.6 | 309.9 | 305.7 | SM 43:2 | SM(d43:2) | 1.4 |
| 13 | 731.6065 | 731.6067 | 0.3 | 284.3 | 288.4 | SM 36:1 | SM(d36:1) | -1.4 |
| 14 | 813.6845 | 813.6849 | 0.6 | 306.8 | 302.2 | SM 42:2 | SM(d42:2) | 1.5 |
| 15 | 703.5750 | 703.5754 | 0.6 | 276.7 | 281.2 | SM 34:1 | SM(d34:1) | -1.6 |
| 16 | 808.5823 | 808.5832 | 1.1 | 302.3 | 296.9 | PC (18:1/18:1) (del9-trans) | PC(36:2) M + Na | 1.8 |
| 17 | 760.5851 | 760.5856 | 0.7 | 293.4 | 287.9 | PC (18:1(9Z)/16:0) | PC(34:1) | 1.9 |
| 18 | 730.5385 | 730.5387 | 0.3 | 283.9 | 278.4 | PC (16:1/16:1) (del9-cis) | PC(32:2) | 2.0 |
| 19 | 811.6687 | 811.6693 | 0.7 | 306.9 | 300.8 | SM 42:3 | SM(d42:3) | 2.0 |
| 20 | 732.5540 | 732.5543 | 0.5 | 283.7 | 277.6 | PC 32:1 | PC(32:1) | 2.2 |
| 21 | 552.4027 | 552.4029 | 0.5 | 241.3 | 247 | 20:0 Lyso PC | LPC(20:0)>LPC(20:0/0:0)_and_LPC(0:0/20:0) | -2.3 |
| 22 | 774.6008 | 774.6013 | 0.6 | 293.4 | 285.8 | PC 35:1 | PC(35:1) | 2.7 |
| 23 | 788.6165 | 788.6169 | 0.6 | 297.1 | 289.4 | PC 36:1 | PC(36:1) | 2.7 |
| 24 | 546.3533 | 546.3535 | 0.4 | 234.8 | 241.5 | 18:0 Lyso PC | LPC(20:3)>LPC(20:3/0:0)_and_LPC(0:0/20:3) | -2.8 |
| 25 | 786.6007 | 786.6012 | 0.7 | 295.5 | 287.1 | PC 36:2 | PC(36:2) | 2.9 |
| 26 | 518.3220 | 518.3222 | 0.5 | 227.3 | 234.3 | 16:0 Lyso PC | LPC(18:3)>LPC(18:3/0:0)_and_LPC(0:0/18:3) | -3.0 |
| 27 | 468.3085 | 468.3090 | 1.1 | 214.0 | 220.8 | LPC 14:0 | LPC(14:0)>LPC(14:0/0:0)_and_LPC(0:0/14:0) | -3.1 |
| 28 | 809.6527 | 809.6512 | 1.9 | 306.3 | 297.2 | SM 40:1 | SM(d40:1) M + Na | 3.1 |
| 29 | 758.5694 | 758.5699 | 0.7 | 290.4 | 280.6 | PC 34:2 | PC(34:2) | 3.5 |

| | | | | | | | | |
|---|---|---|---|---|---|---|---|---|
| 30 | 524.3712 | 524.3716 | 0.7 | 229.2 | 238.8 | 18:0 Lyso PC | LPC(18:0)>LPC(18:0/0:0)_and_LPC(0:0/18:0) | -4.0 |
| 31 | 496.3399 | 496.3403 | 0.9 | 221.8 | 231.4 | 16:0 Lyso PC | LPC(16:0)>LPC(16:0/0:0)_and_LPC(0:0/16:0) | -4.1 |
| 32 | 756.5540 | 756.5543 | 0.4 | 291.2 | 278.2 | PC 34:3 | PC(34:3) | 4.7 |
| 33 | 742.5382 | 742.5387 | 0.7 | 287.9 | 273.5 | PE (18:1/18:1) (del9-cis) | PC(33:3) | 5.3 |
| 34 | 728.5593 | 728.5594 | 0.1 | 288.0 | 273.5 | PE (O-36:3) | PE(O-36:3)>PE(O-18:1/18:2) | 5.3 |
| 35 | 750.5430 | 750.5413 | 2.3 | 294.4 | 279 | PE (O-36:3) | PE(O-36:3)>PE(O-18:2/18:1) | 5.5 |
| 36 | 400.3421 | 400.3427 | 1.5 | 189.7 | 214.7 | Carnitine 16:0 | Car(16:0) | -11.6 |
| 37 | 344.2795 | 344.2782 | 3.8 | 171.8 | 199.5 | Carnitine 12:0 | Car(12:0) | -13.9 |

**Supplementary Table S11. Summary of lipid features with matched $^{Orbi}$CCS to reference $^{TIMS}$CCS dataset for experiments conducted at a resolution of 120,000 and turbopump speed 68%.**

| # | m/z_this study | m/z_database | mzError_ppm | $^{Orbi}$CCS_this study | $^{TIMS}$CCS_database | Name_database | Name_This study | % error |
|---|---|---|---|---|---|---|---|---|
| 1 | 820.7391 | 820.7395 | 0.5 | 309.3 | 309.3 | TG 14:0_16:1_18:1 | TG(48:2)>TG(14:0_16:0_18:2)_and_TG(14:0_16:1_18:1) M + NH3 | 0.0 |
| 2 | 715.5750 | 715.5748 | 0.2 | 286.7 | 286.6 | SM d35:2 | SM(d35:2) | 0.0 |
| 3 | 520.3397 | 520.3405 | 1.4 | 229.2 | 229.1 | LPC 18:2-SN1 | LPC(20:5)>LPC(20:5/0:0)_and_LPC(0:0/20:5) | 0.0 |
| 4 | 746.6062 | 746.6054 | 1.1 | 290.8 | 291 | PC O-34:1 | PC(O-34:1) | -0.1 |
| 5 | 536.3712 | 536.3720 | 1.5 | 237.6 | 237.8 | LPC 19:1-SN1 | LPC(19:1)>LPC(19:1/0:0)_and_LPC(0:0/19:1) | -0.1 |
| 6 | 864.8014 | 864.8019 | 0.6 | 321.1 | 320.8 | TG 16:0_17:0_18:1 | TG(51:1) M + NH3 | 0.1 |
| 7 | 878.8170 | 878.8164 | 0.6 | 324.3 | 324 | TG 16:0_18:0_18:1 | TG(52:1) M + NH3 | 0.1 |
| 8 | 785.6533 | 785.6531 | 0.2 | 301.7 | 301.2 | SM d40:2 | SM(d40:2) | 0.2 |
| 9 | 706.5380 | 706.5382 | 0.3 | 280.0 | 280.5 | PC 30:0 | PC(30:0) | -0.2 |
| 10 | 787.6690 | 787.6697 | 1.0 | 302.4 | 303 | SM d40:1 | SM(d40:1) | -0.2 |
| 11 | 678.5071 | 678.5068 | 0.5 | 274.6 | 275.2 | PC 14:0_14:0 | PC(28:0) | -0.2 |
| 12 | 546.3559 | 546.3547 | 2.2 | 236.1 | 235.4 | LPC 20:3-SN2 | LPC(20:3)>LPC(20:3/0:0)_and_LPC(0:0/20:3) | 0.3 |
| 13 | 746.5696 | 746.5697 | 0.2 | 287.6 | 286.7 | PC 33:1 | PC(33:1) | 0.3 |
| 14 | 799.6690 | 799.6687 | 0.3 | 303.7 | 302.8 | SM d41:2 | SM(d41:2) | 0.3 |
| 15 | 792.7078 | 792.7083 | 0.6 | 302.3 | 303.3 | TG 12:0_16:1_18:1 | TG(46:2) M + NH3 | -0.3 |
| 16 | 850.7858 | 850.7857 | 0.2 | 316.8 | 318 | TG 16:0_16:0_18:1 | TG(50:1)>TG(16:0_16:0_18:1)_M+NH3 | -0.4 |
| 17 | 745.6221 | 745.6221 | 0.1 | 293.9 | 295.2 | SM d37:1 | SM(d37:1) | -0.4 |
| 18 | 874.7858 | 874.7875 | 2.0 | 321.5 | 320 | TG 16:0_18:1_18:2 | TG(52:3)>TG(16:0_18:1_18:2)_and_TG(16:1_18:1_18:1) M + NH3 | 0.5 |
| 19 | 748.6220 | 748.6205 | 2.0 | 293.0 | 294.4 | PC O-34:0 | PC(O-34:0) | -0.5 |
| 20 | 727.5753 | 727.5748 | 0.8 | 285.3 | 286.8 | SM d36:3 | SM(d36:3) | -0.5 |
| 21 | 796.7388 | 796.7391 | 0.4 | 309.5 | 307.8 | TG 14:0_16:0_16:0 | TG(46:0)>TG(14:0_16:0_16:0)_and_TG(12:0_16:0_18:0) M + NH3 | 0.5 |
| 22 | 718.5747 | 718.5748 | 0.1 | 283.5 | 285.2 | PC O-32:1 | PC(O-32:1) | -0.6 |

| | | | | | | | | |
|---|---|---|---|---|---|---|---|---|
| 23 | 862.7857 | 862.7861 | 0.5 | 320.2 | 318.2 | TG 16:0_17:1_18:1 | TG(51:2)>TG(15:0_18:1_18:1)_and_TG(16:0_17:1_18:1)_and_TG(16:1_17:0_18:1) M + NH3 | 0.6 |
| 24 | 848.7701 | 848.7708 | 0.8 | 317.4 | 315.3 | TG 16:0_16:1_18:1 | TG(50:2)>TG(16:0_16:1_18:1)_and_TG(14:0_18:1_18:1) M + NH3 | 0.7 |
| 25 | 822.7548 | 822.7547 | 0.1 | 309.4 | 311.6 | TG 14:0_16:0_18:1 | TG(48:1)>TG(16:0_16:0_16:1)_and_TG(14:0_16:0_18:1) M + NH3 | -0.7 |
| 26 | 520.3399 | 520.3402 | 0.6 | 229.6 | 231.2 | LPC 18:2-SN2 | LPC(18:2)>LPC(18:2/0:0)_and_LPC(0:0/18:2) | -0.7 |
| 27 | 732.5540 | 732.5543 | 0.3 | 285.3 | 283.3 | PC 16:0_16:1 | PC(32:1) | 0.7 |
| 28 | 744.5898 | 744.5890 | 1.1 | 289.7 | 287.6 | PC O-34:2 | PC(O-34:2) | 0.7 |
| 29 | 818.7234 | 818.7238 | 0.5 | 309.7 | 307.4 | TG 16:1_16:1_16:1 | TG(48:3) M + NH3 | 0.7 |
| 30 | 580.4339 | 580.4346 | 1.2 | 252.0 | 253.9 | LPC 22:0-SN1 | LPC(22:0)>LPC(22:0/0:0)_and_LPC(0:0/22:0) | -0.7 |
| 31 | 716.5584 | 716.5588 | 0.5 | 284.5 | 282.4 | PC O-32:2 | PC(O-32:2) | 0.7 |
| 32 | 801.6846 | 801.6854 | 1.0 | 304.1 | 306.5 | SM d41:1 | SM(d41:1) | -0.8 |
| 33 | 733.6223 | 733.6208 | 2.1 | 293.3 | 295.7 | SM d36:0 | SM(d36:0) | -0.8 |
| 34 | 813.6845 | 813.6862 | 2.1 | 309.2 | 306.6 | SM d18:1_24:1 | SM(d42:2) | 0.8 |
| 35 | 846.7546 | 846.7556 | 1.2 | 316.2 | 313.5 | TG 16:1_16:0_18:2 | TG(50:3) M + NH3 | 0.9 |
| 36 | 546.3553 | 546.3552 | 0.2 | 235.9 | 233.8 | LPC 20:3-SN1 | LPC(20:3)>LPC(20:3/0:0)_and_LPC(0:0/20:3) | 0.9 |
| 37 | 548.3712 | 548.3708 | 0.8 | 238.7 | 236.5 | LPC 20:2-SN1 | LPC(20:2)>LPC(20:2/0:0)_and_LPC(0:0/20:2) | 0.9 |
| 38 | 834.7545 | 834.7550 | 0.7 | 315.2 | 312.2 | TG 15:0_16:1_18:1 | TG(49:2) M + NH3 | 1.0 |
| 39 | 872.7701 | 872.7720 | 2.1 | 320.8 | 317.7 | TG 16:0_18:2_18:2 | TG(52:4)>TG(16:0_16:0_20:4)_and_TG(16:1_18:1_18:2)_and_TG(16:0_18:1_18:2) M + NH3 | 1.0 |
| 40 | 734.5697 | 734.5696 | 0.1 | 283.8 | 286.6 | PC 16:0_16:0 | PC(32:0) | -1.0 |
| 41 | 544.3398 | 544.3397 | 0.3 | 236.3 | 233.8 | LPC 20:4-SN2 | LPC(20:4)>LPC(20:4/0:0)_and_LPC(0:0/20:4) | 1.1 |
| 42 | 768.5902 | 768.5895 | 0.9 | 294.8 | 291.7 | PC O-36:4 | PC(O-36:4) | 1.1 |
| 43 | 794.7234 | 794.7233 | 0.1 | 302.1 | 305.5 | TG 14:0_16:0_16:1 | TG(46:1)>TG(12:0_16:0_18:1)_and_TG(10:0_18:0_18:1)_and_TG(14:0_16:0_16:1) M + NH3 | -1.1 |

| | | | | | | | | |
|---|---|---|---|---|---|---|---|---|
| 44 | 721.5858 | 721.5842 | 2.3 | 287.1 | 290.4 | SM 34:0;3O | SM(d34:0-OH) | -1.1 |
| 45 | 766.6920 | 766.6927 | 0.9 | 296.3 | 299.8 | TG 12:0_16:0_16:1 | TG(44:1)>TG(10:0_16:0_18:1) M + NH3 | -1.2 |
| 46 | 760.5852 | 760.5870 | 2.3 | 293.1 | 289.7 | PC 16:0_18:1 | PC(34:1) | 1.2 |
| 47 | 827.7000 | 827.6999 | 0.1 | 313.8 | 310.1 | SM d43:2 | SM(d43:2) | 1.2 |
| 48 | 844.7388 | 844.7396 | 0.9 | 315.4 | 311.6 | TG 14:0_18:2_18:2 | TG(50:4)>TG(16:1_16:1_18:2)_and_TG(16:1_16:1_18:2) M + NH3 | 1.2 |
| 49 | 720.5905 | 720.5898 | 0.9 | 284.3 | 287.9 | PC O-32:0 | PC(O-32:0) | -1.2 |
| 50 | 758.5694 | 758.5718 | 3.2 | 290.0 | 286.4 | PC 16:0_18:2 | PC(34:2) | 1.3 |
| 51 | 550.3868 | 550.3868 | 0.2 | 239.3 | 242.4 | LPC 20:1-SN1 | LPC(20:1)>LPC(20:1/0:0)_and_LPC(0:0/20:1) | -1.3 |
| 52 | 772.5851 | 772.5857 | 0.8 | 293.2 | 289.4 | PC 35:2 | PC(35:2) | 1.3 |
| 53 | 870.7545 | 870.7554 | 1.1 | 320.4 | 316.2 | TG 16:1_18:2_18:2 | TG(52:5)>TG(16:0_18:2_18:3) M + NH3 | 1.3 |
| 54 | 832.7389 | 832.7390 | 0.1 | 315.1 | 310.9 | TG 15:0_16:1_18:2 | TG(49:3) M + NH3 | 1.3 |
| 55 | 544.3398 | 544.3399 | 0.0 | 235.8 | 232.6 | LPC 20:4-SN1 | LPC(20:4)>LPC(20:4/0:0)_and_LPC(0:0/20:4) | 1.4 |
| 56 | 538.3870 | 538.3872 | 0.4 | 239.8 | 243.2 | LPC 19:0-SN2 | LPC(19:0)>LPC(19:0/0:0)_and_LPC(0:0/19:0) | -1.4 |
| 57 | 729.5908 | 729.5904 | 0.5 | 285.7 | 289.7 | SM d18:1_18:1 | SM(d36:2) | -1.4 |
| 58 | 773.6533 | 773.6531 | 0.3 | 296.5 | 300.8 | SM d39:1 | SM(d39:1) | -1.4 |
| 59 | 868.7388 | 868.7387 | 0.1 | 319.4 | 314.8 | TG 16:1_18:2_18:3 | TG(52:6) M + NH3 | 1.5 |
| 60 | 774.6371 | 774.6368 | 0.4 | 301.8 | 297.4 | PC O-36:1 | PC(O-36:1) | 1.5 |
| 61 | 786.6010 | 786.6026 | 2.1 | 297.1 | 292.7 | PC 18:0_18:2 | PC(36:2) | 1.5 |
| 62 | 858.7545 | 858.7544 | 0.1 | 319.8 | 314.9 | TG 15:0_18:2_18:2 | TG(51:4)>TG(16:1_17:1_18:2)_and_TG(15:0_18:2_18:2)_and_TG(15:1_18:1_18:2) M + NH3 | 1.5 |
| 63 | 836.7702 | 836.7703 | 0.2 | 319.5 | 314.6 | TG 15:0_16:0_18:1 | TG(49:1)>TG(15:0_16:0_18:0)_and_TG(16:0_16:0_17:1)_and_TG(16:0_16:1_17:0) M + NH3 | 1.6 |
| 64 | 757.6218 | 757.6216 | 0.3 | 291.0 | 295.6 | SM d38:2 | SM(d38:2) | -1.6 |
| 65 | 811.6686 | 811.6685 | 0.1 | 308.7 | 303.9 | SM d42:3 | SM(d42:3) | 1.6 |

| | | | | | | | | |
|---|---|---|---|---|---|---|---|---|
| 66 | 788.6166 | 788.6167 | 0.2 | 300.1 | 295.4 | PC 18:0_18:1 | PC(36:1) | 1.6 |
| 67 | 842.7190 | 842.7231 | 4.9 | 315.7 | 310.8 | TG 14:0_18:2_18:3 | TG(50:5) M + NH3 | 1.6 |
| 68 | 839.6996 | 839.6997 | 0.2 | 315.6 | 310.6 | SM d44:3 | SM(d44:3) | 1.6 |
| 69 | 770.6057 | 770.6059 | 0.2 | 296.5 | 291.8 | PC O-36:3 | PC(O-36:3) | 1.6 |
| 70 | 784.5851 | 784.5867 | 2.0 | 295.8 | 291.1 | PC 18:1_18:2 | PC(36:3) | 1.6 |
| 71 | 762.6010 | 762.5990 | 2.6 | 297.8 | 292.8 | PC 18:0_16:0 | PC(34:0) | 1.7 |
| 72 | 518.3242 | 518.3242 | 0.0 | 230.0 | 226.1 | LPC 18:3-SN1 | LPC(18:3)>LPC(18:3/0:0)_and_LPC(0:0/18:3) | 1.7 |
| 73 | 759.6378 | 759.6373 | 0.6 | 292.9 | 298.2 | SM d38:1 | SM(d38:1) | -1.8 |
| 74 | 701.5595 | 701.5615 | 2.8 | 278.1 | 283.5 | SM d18:1_16:1 | SM(d34:2) | -1.9 |
| 75 | 673.5282 | 673.5277 | 0.8 | 271.6 | 276.9 | SM d32:2 | SM(d32:2) | -1.9 |
| 76 | 772.6216 | 772.6210 | 0.8 | 300.0 | 294.3 | PC O-36:2 | PC(O-36:2) | 1.9 |
| 77 | 825.6846 | 825.6846 | 0.1 | 311.9 | 306 | SM d43:3 | SM(d43:3) | 1.9 |
| 78 | 902.8167 | 902.8167 | 0.0 | 332.1 | 325.8 | TG 18:1_18:1_18:1 | TG(54:3)>TG(18:1/18:1/18:1) M + NH3 | 1.9 |
| 79 | 800.6160 | 800.6174 | 1.8 | 301.5 | 295.5 | PC 37:2 | PC(37:2) | 2.0 |
| 80 | 730.5385 | 730.5386 | 0.1 | 285.4 | 279.6 | PC 32:2 | PC(32:2) | 2.1 |
| 81 | 704.5226 | 704.5225 | 0.1 | 283.1 | 277.2 | PC 30:1 | PC(30:1) | 2.1 |
| 82 | 888.8013 | 888.8012 | 0.1 | 330.1 | 323.1 | TG 17:0_18:1_18:2 | TG(53:3)>TG(17:0_18:1_18:2)_and_TG(17:1_18:1_18:1)_and_TG(16:1_18:1_19:1)_and_TG(16:0_18:1_19:2) M + NH3 | 2.2 |
| 83 | 717.5906 | 717.5905 | 0.2 | 283.3 | 289.8 | SM d35:1 | SM(d35:1) | -2.2 |
| 84 | 794.6051 | 794.6055 | 0.5 | 302.3 | 295.7 | PC O-38:5 | PC(O-38:5) | 2.2 |
| 85 | 904.8326 | 904.8316 | 1.2 | 335.1 | 327.7 | TG 18:0_18:1_18:1 | TG(54:2)>TG(16:0_18:1_20:1)_and_TG(18:0_18:1_18:1) M + NH3 | 2.3 |
| 86 | 552.4027 | 552.4025 | 0.3 | 240.8 | 246.4 | LPC 20:0-SN1 | LPC(20:0)>LPC(20:0/0:0)_and_LPC(0:0/20:0) | -2.3 |
| 87 | 816.6475 | 816.6468 | 0.8 | 307.7 | 300.8 | PC 38:1 | PC(38:1) | 2.3 |
| 88 | 689.5595 | 689.5599 | 0.6 | 277.3 | 283.8 | SM d33:1 | SM(d33:1) | -2.3 |

| | | | | | | | | |
|---|---|---|---|---|---|---|---|---|
| 89 | 960.8950 | 960.8946 | 0.4 | 346.2 | 338.4 | TG 18:1_18:1_22:0 | TG(58:2) M + NH3 | 2.3 |
| 90 | 494.3242 | 494.3243 | 0.2 | 221.7 | 227.1 | LPC 16:1-SN1 | LPC(16:1)>LPC(16:1/0:0)_and_LPC(0:0/16:1) | -2.4 |
| 91 | 677.5598 | 677.5582 | 2.3 | 278.2 | 285 | SM d32:0 | SM(d32:0) | -2.4 |
| 92 | 900.8012 | 900.8010 | 0.3 | 331.5 | 323.8 | TG 18:1_18:1_18:2 | TG(54:4)>TG(18:1_18:1_18:2) M + NH3 | 2.4 |
| 93 | 898.7858 | 898.7855 | 0.3 | 329.9 | 322 | TG 18:1_18:2_18:2 | TG(54:5)>TG(18:1_18:2_18:2)_and_TG(16:0_16:0_22:5)_and_TG(16:0_18:1_20:4) M + NH3 | 2.5 |
| 94 | 522.3552 | 522.3562 | 1.9 | 229.2 | 235.2 | LPC 18:1-SN1 | LPC(18:1)>LPC(18:1/0:0)_and_LPC(0:0/18:1) | -2.6 |
| 95 | 808.5850 | 808.5848 | 0.1 | 301.8 | 294.2 | PC 38:5 | PC(38:5) | 2.6 |
| 96 | 782.5694 | 782.5701 | 0.8 | 298.1 | 290.4 | PC 16:0_20:4 | PC(36:4) | 2.7 |
| 97 | 810.6008 | 810.6011 | 0.4 | 304.7 | 296.7 | PC 18:0_20:4 | PC(38:4) | 2.7 |
| 98 | 886.7856 | 886.7852 | 0.5 | 329.7 | 321.1 | TG 17:1_18:1_18:2 | TG(53:4)>TG(17:0_18:2_18:2)_and_TG(17:1_18:1_18:2) M + NH3 | 2.7 |
| 99 | 802.6323 | 802.6332 | 1.1 | 306.4 | 298.3 | PC 37:1 | PC(37:1) | 2.7 |
| 100 | 494.3242 | 494.3244 | 0.3 | 222.1 | 228.5 | LPC 16:1-SN2 | LPC(16:1)>LPC(16:1/0:0)_and_LPC(0:0/16:1) | -2.8 |
| 101 | 890.8167 | 890.8168 | 0.1 | 334.1 | 324.9 | TG 17:0_18:1_18:1 | TG(53:2) M + NH3 | 2.8 |
| 102 | 568.3398 | 568.3397 | 0.3 | 243.1 | 236.4 | LPC 22:6-SN2 | LPC(22:6)>LPC(22:6/0:0)_and_LPC(0:0/22:6) | 2.8 |
| 103 | 896.7701 | 896.7699 | 0.2 | 329.5 | 320.4 | TG 18:2_18:2_18:2 | TG(52:3)>TG(16:0_18:1_18:2)_and_TG(16:1_18:1_18:1) M + NH3 | 2.9 |
| 104 | 731.6065 | 731.6062 | 0.4 | 284.2 | 292.6 | SM d18:1_18:0 | SM(d36:1) | -2.9 |
| 105 | 714.6188 | 714.6185 | 0.4 | 287.3 | 295.9 | CE 22:6 | CE(22:6)M+NH3 | -2.9 |
| 106 | 946.8793 | 946.8787 | 0.7 | 345.6 | 335.8 | TG 18:1_18:1_21:0 | TG(57:2) M + NH3 | 2.9 |
| 107 | 958.8792 | 958.8787 | 0.5 | 346.3 | 336.4 | TG 18:1_18:2_22:0 | TG(58:3) M + NH3 | 2.9 |
| 108 | 572.3710 | 572.3710 | 0.1 | 245.7 | 238.6 | LPC 22:4-SN1 | LPC(22:4)>LPC(22:4/0:0)_and_LPC(0:0/22:4) | 3.0 |
| 109 | 728.5229 | 728.5231 | 0.3 | 285.4 | 277.1 | PC 32:3 | PC(32:3) | 3.0 |
| 110 | 780.5902 | 780.5903 | 0.1 | 299.4 | 290.7 | PC O-37:5 | PC(O-37:5) | 3.0 |
| 111 | 894.7543 | 894.7539 | 0.4 | 328.1 | 318.5 | TG 18:2_18:2_18:3 | TG(54:7)>TG(16:0_16:1_22:6)_and_TG(16:0_18:2_20:5) M + NH3 | 3.0 |

| | | | | | | | | |
|---|---|---|---|---|---|---|---|---|
| 112 | 570.3555 | 570.3553 | 0.3 | 243.6 | 236.4 | LPC 22:5-SN1 | LPC(22:5)>LPC(22:5/0:0)_and_LPC(0:0/ 22:5) | 3.0 |
| 113 | 703.5750 | 703.5778 | 4.0 | 277.6 | 286.3 | SM d18:1_16:0 | SM(d34:1) | -3.0 |
| 114 | 705.5904 | 705.5891 | 1.8 | 281.2 | 290.1 | SM d34:0 | SM(d34:0) | -3.1 |
| 115 | 796.6217 | 796.6210 | 0.9 | 306.7 | 297.5 | PC O-38:4 | PC(O-38:4) | 3.1 |
| 116 | 884.7701 | 884.7688 | 1.4 | 329.6 | 319.5 | TG 17:1_18:2_18:2 | TG(53:5)>TG(17:1_18:2_18:2) M + NH3 | 3.2 |
| 117 | 814.6321 | 814.6311 | 1.2 | 307.9 | 298.4 | PC 38:2 | PC(38:2) | 3.2 |
| 118 | 522.3555 | 522.3554 | 0.2 | 229.4 | 236.9 | LPC 18:1-SN2 | LPC(18:1)>LPC(18:1/0:0)_and_LPC(0:0/ 18:1) | -3.2 |
| 119 | 675.5438 | 675.5447 | 1.4 | 271.5 | 280.5 | SM d32:1 | SM(d32:1) | -3.2 |
| 120 | 756.5540 | 756.5540 | 0.0 | 292.6 | 283.3 | PC 34:3 | PC(34:3) | 3.3 |
| 121 | 482.3243 | 482.3243 | 0.2 | 220.7 | 228.3 | LPC 15:0-SN1 | LPC(15:0)>LPC(15:0/0:0)_and_LPC(0:0/ 15:0) | -3.3 |
| 122 | 812.6165 | 812.6163 | 0.2 | 307.1 | 297 | PC 38:3 | PC(38:3) | 3.4 |
| 123 | 782.5696 | 782.5695 | 0.1 | 297.3 | 287.5 | PC 18:2_18:2 Cis | PC(36:4) | 3.4 |
| 124 | 568.3399 | 568.3396 | 0.6 | 243.3 | 235.2 | LPC 22:6-SN1 | LPC(22:6)>LPC(22:6/0:0)_and_LPC(0:0/ 22:6) | 3.5 |
| 125 | 824.6166 | 824.6160 | 0.8 | 310.1 | 299.7 | PC 39:4 | PC(39:4) | 3.5 |
| 126 | 934.8792 | 934.8793 | 0.2 | 346.9 | 335.2 | TG 16:0_18:1_22:0 | TG(56:1) M + NH3 | 3.5 |
| 127 | 780.5540 | 780.5542 | 0.3 | 297.9 | 287.3 | PC 36:5 | PC(36:5) | 3.7 |
| 128 | 766.5751 | 766.5747 | 0.5 | 300.3 | 289.5 | PC O-36:5 (PC O-16:0_20:5) | PC(O-36:5) | 3.7 |
| 129 | 768.5539 | 768.5540 | 0.2 | 296.9 | 286.2 | PC 35:4 | PC(35:4) | 3.7 |
| 130 | 754.5384 | 754.5381 | 0.4 | 292.5 | 281.9 | PC 34:4 | PC(34:4) | 3.8 |
| 131 | 820.5850 | 820.5850 | 0.0 | 308.1 | 296.6 | PC 39:6 | PC(39:6) | 3.9 |
| 132 | 948.8951 | 948.8944 | 0.8 | 350.6 | 337.2 | TG 16:0_16:1_25:0 | TG(57:1)>TG(16:0_18:1_23:0)_and_TG( 18:0_18:1_21:0) M + NH3 | 4.0 |
| 133 | 834.6007 | 834.6002 | 0.6 | 312.0 | 299.5 | PC 18:0_22:6 | PC(40:6) | 4.2 |
| 134 | 924.8013 | 924.8006 | 0.7 | 341.2 | 327.6 | TG 18:1_18:1_20:4 | TG(56:6)>TG(18:1_18:1_20:4)_and_TG( 16:0_18:1_22:5)_and_TG(18:1_18:2_20: 3) M + NH3 | 4.2 |

| | | | | | | | | |
|---|---|---|---|---|---|---|---|---|
| 135 | 524.3710 | 524.3717 | 1.2 | 229.8 | 240.1 | LPC 18:0-SN1 | LPC(18:0)>LPC(18:0/0:0)_and_LPC(0:0/18:0) | -4.3 |
| 136 | 718.5385 | 718.5383 | 0.3 | 291.8 | 279.7 | PC 17:0_14:1 | PC(31:1) | 4.3 |
| 137 | 770.5694 | 770.5696 | 0.3 | 300.5 | 288 | PC 35:3 | PC(35:3) | 4.3 |
| 138 | 690.6186 | 690.6187 | 0.1 | 280.7 | 294 | CE 20:4 | CE(20:4)M+NH3 | -4.5 |
| 139 | 468.3085 | 468.3084 | 0.1 | 214.2 | 224.6 | LPC 14:0-SN1 | LPC(14:0)>LPC(14:0/0:0)_and_LPC(0:0/14:0) | -4.6 |
| 140 | 524.3713 | 524.3715 | 0.3 | 229.9 | 241 | LPC 18:0-SN2 | LPC(18:0)>LPC(18:0/0:0)_and_LPC(0:0/18:0) | -4.6 |
| 141 | 510.3556 | 510.3555 | 0.2 | 225.4 | 236.4 | LPC 17:0-SN1 | LPC(17:0)>LPC(17:0/0:0)_and_LPC(0:0/17:0) | -4.7 |
| 142 | 926.8172 | 926.8162 | 1.1 | 344.6 | 329.2 | TG 18:0_18:1_20:4 | TG(56:5)>TG(16:0_18:1_22:4) M + NH3 | 4.7 |
| 143 | 876.8016 | 876.8023 | 0.9 | 337.1 | 321.9 | TG 16:0_18:1_18:1 | TG(52:2)>TG(16:0_18:1_18:1) M + NH3 | 4.7 |
| 144 | 774.6008 | 774.6011 | 0.3 | 306.3 | 292.4 | PC 35:1 | PC(35:1) | 4.8 |
| 145 | 468.3085 | 468.3087 | 0.5 | 214.5 | 225.5 | LPC 14:0-SN2 | LPC(14:0)>LPC(14:0/0:0)_and_LPC(0:0/14:0) | -4.9 |
| 146 | 806.5695 | 806.5689 | 0.8 | 305.7 | 291.4 | PC 38:6 (PC 18:2_20:4) | PC(38:6) | 4.9 |
| 147 | 496.3399 | 496.3405 | 1.2 | 220.7 | 232.1 | LPC 16:0-SN1 | LPC(16:0)>LPC(16:0/0:0)_and_LPC(0:0/16:0) | -4.9 |
| 148 | 818.6057 | 818.6057 | 0.1 | 313.2 | 298 | PC O-40:7 | PC(O-40:7) | 5.1 |
| 149 | 848.6526 | 848.6528 | 0.2 | 319.9 | 304.2 | PC O-42:6 | PC(O-42:6) | 5.1 |
| 150 | 944.7701 | 944.7694 | 0.7 | 343.6 | 326.7 | TG 16:0_20:4_22:6 | TG(58:10) M + NH3 | 5.2 |
| 151 | 866.6638 | 866.6643 | 0.6 | 325.0 | 308.8 | PC 42:4 | PC(42:4) | 5.3 |
| 152 | 832.5848 | 832.5846 | 0.3 | 312.6 | 296.8 | PC 40:7 | PC(40:7) | 5.3 |
| 153 | 820.6211 | 820.6209 | 0.2 | 315.0 | 298.9 | PC O-40:6 | PC(O-40:6) | 5.4 |
| 154 | 496.3399 | 496.3402 | 0.7 | 220.4 | 233.7 | LPC 16:0-SN2 | LPC(16:0)>LPC(16:0/0:0)_and_LPC(0:0/16:0) | -5.7 |
| 155 | 824.6523 | 824.6529 | 0.7 | 321.5 | 302.8 | PC O-40:4 | PC(O-40:4) | 6.2 |
| 156 | 830.5696 | 830.5690 | 0.7 | 312.5 | 294 | PC 20:4_20:4 Cis | PC(40:8) | 6.3 |
| 157 | 864.6475 | 864.6473 | 0.2 | 325.8 | 305.5 | PC 42:5 | PC(42:5) | 6.7 |

| | | | | | | | | |
|---|---|---|---|---|---|---|---|---|
| 158 | 856.5850 | 856.5846 | 0.4 | 321.5 | 297.6 | PC 42:9 | PC(42:9) | 8.0 |

**Supplementary Table S12. Summary of lipid features with matched $^{Orbi}$CCS to reference $^{DT}$CCS dataset for experiments conducted at a resolution of 120,000 and turbopump speed 68%.**

| # | m/z_this study | m/z_database | mzError_ppm | $^{Orbi}$CCS_this study | $^{DT}$CCS_database | Name_database | Name_This study | % error |
|---|---|---|---|---|---|---|---|---|
| 1 | 729.5908 | 729.5910 | 0.3 | 285.7 | 285.3 | SM 36:2 | SM(d36:2) | 0.1 |
| 2 | 759.6378 | 759.6380 | 0.3 | 292.9 | 293.4 | SM 38:1 | SM(d38:1) | -0.2 |
| 3 | 773.6533 | 773.6536 | 0.4 | 296.5 | 297 | SM 39:1 | SM(d39:1) | -0.2 |
| 4 | 538.3870 | 538.3872 | 0.4 | 239.8 | 240.5 | LPC 19:0 | LPC(19:0)>LPC(19:0/0:0)_and_LPC(0:0/19:0) | -0.3 |
| 5 | 801.6846 | 801.6849 | 0.3 | 304.1 | 302.3 | SM 41:1 | SM(d41:1) | 0.6 |
| 6 | 787.6690 | 787.6693 | 0.4 | 302.4 | 299.1 | SM 40:1 | SM(d40:1) | 1.1 |
| 7 | 799.6690 | 799.6693 | 0.4 | 303.6 | 300.1 | SM 41:2 | SM(d41:2) | 1.2 |
| 8 | 703.5750 | 703.5754 | 0.6 | 277.6 | 281.2 | SM 34:1 | SM(d34:1) | -1.3 |
| 9 | 786.6010 | 786.6012 | 0.3 | 297.1 | 293.3 | PC (18:1/18:1) (del9-trans) | PC(36:2) | 1.3 |
| 10 | 745.6221 | 745.6223 | 0.3 | 293.9 | 289.8 | SM 37:1 | SM(d37:1) | 1.4 |
| 11 | 731.6065 | 731.6067 | 0.3 | 284.2 | 288.4 | SM 36:1 | SM(d36:1) | -1.5 |
| 12 | 785.6534 | 785.6536 | 0.3 | 301.6 | 296.8 | SM 40:2 | SM(d40:2) | 1.6 |
| 13 | 522.3552 | 522.3559 | 1.4 | 229.2 | 233.2 | LPC 18:1 | LPC(18:1)>LPC(18:1/0:0)_and_LPC(0:0/18:1) | -1.7 |
| 14 | 760.5852 | 760.5856 | 0.5 | 293.1 | 287.9 | PC (18:1(9Z)/16:0) | PC(34:1) | 1.8 |
| 15 | 813.6845 | 813.6849 | 0.5 | 309.2 | 302.2 | SM 42:2 | SM(d42:2) | 2.3 |
| 16 | 730.5385 | 730.5387 | 0.3 | 285.4 | 278.4 | PC (16:1/16:1) (del9-cis) | PC(32:2) | 2.5 |
| 17 | 552.4027 | 552.4029 | 0.5 | 240.8 | 247 | 20:0 Lyso PC | LPC(20:0)>LPC(20:0/0:0)_and_LPC(0:0/20:0) | -2.5 |
| 18 | 827.7001 | 827.7006 | 0.6 | 313.7 | 305.7 | SM 43:2 | SM(d43:2) | 2.6 |
| 19 | 811.6686 | 811.6693 | 0.8 | 308.7 | 300.8 | SM 42:3 | SM(d42:3) | 2.6 |
| 20 | 732.5540 | 732.5543 | 0.4 | 285.3 | 277.6 | PC 32:1 | PC(32:1) | 2.8 |
| 21 | 808.5822 | 808.5832 | 1.2 | 305.7 | 296.9 | PC (18:1/18:1) (del9-trans) | PC(36:2) M + Na | 3.0 |
| 22 | 468.3085 | 468.3090 | 1.1 | 214.2 | 220.8 | LPC 14:0 | LPC(14:0)>LPC(14:0/0:0)_and_LPC(0:0/14:0) | -3.0 |
| 23 | 786.6007 | 786.6012 | 0.6 | 296.0 | 287.1 | PC 36:2 | PC(36:2) | 3.1 |
| 24 | 758.5694 | 758.5699 | 0.7 | 290.0 | 280.6 | PC 34:2 | PC(34:2) | 3.4 |
| 25 | 788.6166 | 788.6169 | 0.4 | 300.1 | 289.4 | PC 36:1 | PC(36:1) | 3.7 |
| 26 | 524.3713 | 524.3716 | 0.6 | 229.9 | 238.8 | 18:0 Lyso PC | LPC(18:0)>LPC(18:0/0:0)_and_LPC(0:0/18:0) | -3.7 |
| 27 | 780.5511 | 780.5519 | 1.0 | 295.0 | 284.2 | PC 34:2 | PC(34:2) M + Na | 3.8 |
| 28 | 744.5543 | 744.5543 | 0.1 | 290.0 | 279.1 | PE (18:1/18:1) (del9-cis) | PC(33:2) | 3.9 |
| 29 | 496.3399 | 496.3403 | 0.9 | 220.4 | 231.4 | 16:0 Lyso PC | LPC(16:0)>LPC(16:0/0:0)_and_LPC(0:0/16:0) | -4.8 |

| | | | | | | | | |
|---|---|---|---|---|---|---|---|---|
| 30 | 756.5540 | 756.5543 | 0.4 | 292.6 | 278.2 | PC 34:3 | PC(34:3) | 5.2 |
| 31 | 728.5591 | 728.5594 | 0.4 | 288.0 | 273.5 | PE (O-36:3) | PE(O-36:3)>PE(O-18:1/18:2) | 5.3 |
| 32 | 806.5695 | 806.5675 | 2.4 | 305.6 | 288.2 | PC 36:3 | PC(36:3) | 6.1 |
| 33 | 774.6008 | 774.6013 | 0.6 | 306.3 | 285.8 | PC 35:1 | PC(35:1) | 7.2 |
| 34 | 400.3421 | 400.3427 | 1.4 | 188.6 | 214.7 | Carnitine 16:0 | Car(16:0) | -12.1 |
| 35 | 344.2795 | 344.2782 | 3.8 | 171.6 | 199.5 | Carnitine 12:0 | Car(12:0) | -14.0 |

**Supplementary Table S13. Summary of lipid features with matched $^{Orbi}$CCS to reference $^{TIMS}$CCS dataset for experiments conducted at a resolution of 90,000 and turbopump speed 68%.**

| # | m/z_this study | m/z_database | mzError_ppm | $^{Orbi}$CCS_this study | $^{TIMS}$CCS_database | Name_database | Name_This study | % error |
|---|---|---|---|---|---|---|---|---|
| 1 | 678.5070 | 678.5068 | 0.3 | 275.1 | 275.2 | PC 14:0_14:0 | PC(28:0) | 0.0 |
| 2 | 834.7545 | 834.7550 | 0.6 | 312.4 | 312.2 | TG 15:0_16:1_18:1 | TG(49:2) M + NH3 | 0.1 |
| 3 | 878.8170 | 878.8164 | 0.7 | 323.7 | 324 | TG 16:0_18:0_18:1 | TG(52:1) M + NH3 | -0.1 |
| 4 | 787.6688 | 787.6697 | 1.1 | 302.7 | 303 | SM d40:1 | SM(d40:1) | -0.1 |
| 5 | 520.3397 | 520.3405 | 1.5 | 229.3 | 229.1 | LPC 18:2-SN1 | LPC(20:5)>LPC(20:5/0:0)_and_LPC(0:0/20:5) | 0.1 |
| 6 | 785.6533 | 785.6531 | 0.2 | 300.8 | 301.2 | SM d40:2 | SM(d40:2) | -0.1 |
| 7 | 536.3712 | 536.3720 | 1.6 | 238.2 | 237.8 | LPC 19:1-SN1 | LPC(19:1)>LPC(19:1/0:0)_and_LPC(0:0/19:1) | 0.2 |
| 8 | 862.7858 | 862.7861 | 0.4 | 318.8 | 318.2 | TG 16:0_17:1_18:1 | TG(51:2)>TG(15:0_18:1_18:1)_and_TG(16:0_17:1_18:1)_and_TG(16:1_17:0_18:1) M + NH3 | 0.2 |
| 9 | 874.7858 | 874.7875 | 2.0 | 320.6 | 320 | TG 16:0_18:1_18:2 | TG(52:3)>TG(16:0_18:1_18:2)_and_TG(16:1_18:1_18:1) M + NH3 | 0.2 |
| 10 | 820.7390 | 820.7395 | 0.5 | 308.6 | 309.3 | TG 14:0_16:1_18:1 | TG(48:2)>TG(14:0_16:0_18:2)_and_TG(14:0_16:1_18:1) M + NH3 | -0.2 |
| 11 | 846.7545 | 846.7556 | 1.3 | 314.2 | 313.5 | TG 16:1_16:0_18:2 | TG(50:3) M + NH3 | 0.2 |
| 12 | 746.5697 | 746.5697 | 0.0 | 285.9 | 286.7 | PC 33:1 | PC(33:1) | -0.3 |
| 13 | 818.7235 | 818.7238 | 0.4 | 308.3 | 307.4 | TG 16:1_16:1_16:1 | TG(48:3) M + NH3 | 0.3 |
| 14 | 848.7701 | 848.7708 | 0.8 | 314.2 | 315.3 | TG 16:0_16:1_18:1 | TG(50:2)>TG(16:0_16:1_18:1)_and_TG(14:0_18:1_18:1) M + NH3 | -0.3 |
| 15 | 827.7000 | 827.6999 | 0.1 | 311.2 | 310.1 | SM d43:2 | SM(d43:2) | 0.4 |
| 16 | 799.6690 | 799.6687 | 0.4 | 303.9 | 302.8 | SM d41:2 | SM(d41:2) | 0.4 |
| 17 | 746.6063 | 746.6054 | 1.2 | 289.9 | 291 | PC O-34:1 | PC(O-34:1) | -0.4 |
| 18 | 732.5540 | 732.5543 | 0.3 | 284.4 | 283.3 | PC 16:0_16:1 | PC(32:1) | 0.4 |
| 19 | 876.8015 | 876.8023 | 0.9 | 320.5 | 321.9 | TG 16:0_18:1_18:1 | TG(52:2)>TG(16:0_18:1_18:1) M + NH3 | -0.4 |
| 20 | 733.6220 | 733.6208 | 1.7 | 294.4 | 295.7 | SM d36:0 | SM(d36:0) | -0.5 |
| 21 | 832.7390 | 832.7390 | 0.1 | 312.3 | 310.9 | TG 15:0_16:1_18:2 | TG(49:3) M + NH3 | 0.5 |
| 22 | 744.5898 | 744.5890 | 1.1 | 289.0 | 287.6 | PC O-34:2 | PC(O-34:2) | 0.5 |

| | | | | | | | | |
|---|---|---|---|---|---|---|---|---|
| 23 | 706.5380 | 706.5382 | 0.3 | 279.1 | 280.5 | PC 30:0 | PC(30:0) | -0.5 |
| 24 | 836.7702 | 836.7703 | 0.2 | 316.2 | 314.6 | TG 15:0_16:0_18:1 | TG(49:1)>TG(15:0_16:0_18:0)_and_TG(16:0_16:0_17:1)_and_TG(16:0_16:1_17:0) M + NH3 | 0.5 |
| 25 | 796.7390 | 796.7391 | 0.2 | 309.3 | 307.8 | TG 14:0_16:0_16:0 | TG(46:0)>TG(14:0_16:0_16:0)_and_TG(12:0_16:0_18:0) M + NH3 | 0.5 |
| 26 | 520.3398 | 520.3402 | 0.7 | 230.0 | 231.2 | LPC 18:2-SN2 | LPC(18:2)>LPC(18:2/0:0)_and_LPC(0:0/18:2) | -0.5 |
| 27 | 792.7076 | 792.7083 | 0.8 | 301.6 | 303.3 | TG 12:0_16:1_18:1 | TG(46:2) M + NH3 | -0.5 |
| 28 | 718.5748 | 718.5748 | 0.0 | 283.4 | 285.2 | PC O-32:1 | PC(O-32:1) | -0.6 |
| 29 | 813.6845 | 813.6862 | 2.1 | 308.6 | 306.6 | SM d18:1_24:1 | SM(d42:2) | 0.7 |
| 30 | 872.7702 | 872.7720 | 2.1 | 319.8 | 317.7 | TG 16:0_18:2_18:2 | TG(52:4)>TG(16:0_16:0_20:4)_and_TG(16:1_18:1_18:2)_and_TG(16:0_18:1_18:2) M + NH3 | 0.7 |
| 31 | 715.5749 | 715.5748 | 0.1 | 288.6 | 286.6 | SM d35:2 | SM(d35:2) | 0.7 |
| 32 | 716.5583 | 716.5588 | 0.7 | 284.6 | 282.4 | PC O-32:2 | PC(O-32:2) | 0.8 |
| 33 | 768.5903 | 768.5895 | 1.0 | 294.1 | 291.7 | PC O-36:4 | PC(O-36:4) | 0.8 |
| 34 | 844.7388 | 844.7396 | 1.0 | 314.2 | 311.6 | TG 14:0_18:2_18:2 | TG(50:4)>TG(16:1_16:1_18:2)_and_TG(16:1_16:1_18:2) M + NH3 | 0.8 |
| 35 | 801.6847 | 801.6854 | 0.9 | 303.9 | 306.5 | SM d41:1 | SM(d41:1) | -0.9 |
| 36 | 727.5753 | 727.5748 | 0.7 | 284.3 | 286.8 | SM d36:3 | SM(d36:3) | -0.9 |
| 37 | 850.7858 | 850.7857 | 0.2 | 315.2 | 318 | TG 16:0_16:0_18:1 | TG(50:1)>TG(16:0_16:0_18:1)_M+NH3 | -0.9 |
| 38 | 822.7546 | 822.7547 | 0.1 | 308.9 | 311.6 | TG 14:0_16:0_18:1 | TG(48:1)>TG(16:0_16:0_16:1)_and_TG(14:0_16:0_18:1) M + NH3 | -0.9 |
| 39 | 858.7545 | 858.7544 | 0.1 | 317.7 | 314.9 | TG 15:0_18:2_18:2 | TG(51:4)>TG(16:1_17:1_18:2)_and_TG(15:0_18:2_18:2)_and_TG(15:1_18:1_18:2) M + NH3 | 0.9 |
| 40 | 720.5540 | 720.5540 | 0.0 | 285.7 | 283.1 | PC 31:0 | PC(31:0) | 0.9 |
| 41 | 546.3558 | 546.3547 | 2.1 | 237.6 | 235.4 | LPC 20:3-SN2 | LPC(20:3)>LPC(20:3/0:0)_and_LPC(0:0/20:3) | 0.9 |
| 42 | 748.6221 | 748.6205 | 2.1 | 291.5 | 294.4 | PC O-34:0 | PC(O-34:0) | -1.0 |
| 43 | 786.6007 | 786.6026 | 2.5 | 295.8 | 292.7 | PC 18:0_18:2 | PC(36:2) | 1.0 |
| 44 | 550.3867 | 550.3868 | 0.1 | 239.9 | 242.4 | LPC 20:1-SN1 | LPC(20:1)>LPC(20:1/0:0)_and_LPC(0:0/20:1) | -1.0 |

| | | | | | | | | |
|---|---|---|---|---|---|---|---|---|
| 45 | 870.7545 | 870.7554 | 1.1 | 319.6 | 316.2 | TG 16:1_18:2_18:2 | TG(52:5)>TG(16:0_18:2_18:3) M + NH3 | 1.1 |
| 46 | 788.6165 | 788.6167 | 0.2 | 298.6 | 295.4 | PC 18:0_18:1 | PC(36:1) | 1.1 |
| 47 | 734.5698 | 734.5696 | 0.2 | 283.4 | 286.6 | PC 16:0_16:0 | PC(32:0) | -1.1 |
| 48 | 811.6686 | 811.6685 | 0.1 | 307.4 | 303.9 | SM d42:3 | SM(d42:3) | 1.2 |
| 49 | 760.5850 | 760.5870 | 2.6 | 293.1 | 289.7 | PC 16:0_18:1 | PC(34:1) | 1.2 |
| 50 | 774.6370 | 774.6368 | 0.3 | 300.9 | 297.4 | PC O-36:1 | PC(O-36:1) | 1.2 |
| 51 | 745.6220 | 745.6221 | 0.1 | 291.7 | 295.2 | SM d37:1 | SM(d37:1) | -1.2 |
| 52 | 580.4340 | 580.4346 | 1.1 | 256.9 | 253.9 | LPC 22:0-SN1 | LPC(22:0)>LPC(22:0/0:0)_and_LPC(0:0/22:0) | 1.2 |
| 53 | 784.5851 | 784.5867 | 2.1 | 294.6 | 291.1 | PC 18:1_18:2 | PC(36:3) | 1.2 |
| 54 | 688.6027 | 688.6026 | 0.2 | 287.8 | 291.4 | CE 20:5 | CE(20:5)M+NH3 | -1.2 |
| 55 | 538.3870 | 538.3872 | 0.2 | 240.1 | 243.2 | LPC 19:0-SN2 | LPC(19:0)>LPC(19:0/0:0)_and_LPC(0:0/19:0) | -1.3 |
| 56 | 772.5850 | 772.5857 | 0.9 | 293.2 | 289.4 | PC 35:2 | PC(35:2) | 1.3 |
| 57 | 839.6997 | 839.6997 | 0.0 | 314.7 | 310.6 | SM d44:3 | SM(d44:3) | 1.3 |
| 58 | 758.5693 | 758.5718 | 3.2 | 290.2 | 286.4 | PC 16:0_18:2 | PC(34:2) | 1.3 |
| 59 | 902.8168 | 902.8167 | 0.2 | 330.3 | 325.8 | TG 18:1_18:1_18:1 | TG(54:3)>TG(18:0_18:1_18:2)_and_TG(18:1/18:1/18:1)_and_TG(16:0_18:2_20:1) M + NH3 | 1.4 |
| 60 | 544.3398 | 544.3397 | 0.2 | 237.1 | 233.8 | LPC 20:4-SN2 | LPC(20:4)>LPC(20:4/0:0)_and_LPC(0:0/20:4) | 1.4 |
| 61 | 720.5905 | 720.5898 | 0.9 | 283.8 | 287.9 | PC O-32:0 | PC(O-32:0) | -1.4 |
| 62 | 482.3241 | 482.3245 | 0.7 | 220.1 | 223.3 | LPE 18:0 SN1 | LPC(15:0)>LPC(15:0/0:0)_and_LPC(0:0/15:0) | -1.4 |
| 63 | 809.6527 | 809.6523 | 0.4 | 305.8 | 301.5 | | SM(d40:1) M + Na | 1.4 |
| 64 | 704.5226 | 704.5225 | 0.1 | 281.2 | 277.2 | PC 30:1 | PC(30:1) | 1.4 |
| 65 | 546.3555 | 546.3552 | 0.6 | 237.3 | 233.8 | LPC 20:3-SN1 | LPC(20:3)>LPC(20:3/0:0)_and_LPC(0:0/20:3) | 1.5 |
| 66 | 888.8013 | 888.8012 | 0.0 | 328.0 | 323.1 | TG 17:0_18:1_18:2 | TG(53:3)>TG(17:0_18:1_18:2)_and_TG(17:1_18:1_18:1)_and_TG(16:1_18:1_19:1)_and_TG(16:0_18:1_19:2) M + NH3 | 1.5 |

| | | | | | | | | |
|---|---|---|---|---|---|---|---|---|
| 67 | 762.6010 | 762.5990 | 2.6 | 297.3 | 292.8 | PC 18:0_16:0 | PC(34:0) | 1.5 |
| 68 | 766.6921 | 766.6927 | 0.8 | 295.2 | 299.8 | TG 12:0_16:0_16:1 | TG(44:1)>TG(10:0_16:0_18:1) M + NH3 | -1.5 |
| 69 | 721.5858 | 721.5842 | 2.2 | 285.9 | 290.4 | SM 34:0;3O | SM(d34:0-OH) | -1.5 |
| 70 | 825.6845 | 825.6846 | 0.1 | 310.7 | 306 | SM d43:3 | SM(d43:3) | 1.5 |
| 71 | 868.7389 | 868.7387 | 0.2 | 319.7 | 314.8 | TG 16:1_18:2_18:3 | TG(52:6) M + NH3 | 1.6 |
| 72 | 794.7235 | 794.7233 | 0.3 | 300.7 | 305.5 | TG 14:0_16:0_16:1 | TG(46:1)>TG(12:0_16:0_18:1)_and_TG(10:0_18:0_18:1)_and_TG(14:0_16:0_16:1) M + NH3 | -1.6 |
| 73 | 729.5906 | 729.5904 | 0.3 | 285.1 | 289.7 | SM d18:1_18:1 | SM(d36:2) | -1.6 |
| 74 | 730.5384 | 730.5386 | 0.2 | 284.0 | 279.6 | PC 32:2 | PC(32:2) | 1.6 |
| 75 | 900.8012 | 900.8010 | 0.3 | 329.0 | 323.8 | TG 18:1_18:1_18:2 | TG(54:4)>TG(18:1_18:1_18:2) M + NH3 | 1.6 |
| 76 | 794.6058 | 794.6055 | 0.5 | 300.5 | 295.7 | PC O-38:5 | PC(O-38:5) | 1.6 |
| 77 | 770.6058 | 770.6059 | 0.0 | 296.6 | 291.8 | PC O-36:3 | PC(O-36:3) | 1.6 |
| 78 | 548.3713 | 548.3708 | 0.9 | 240.4 | 236.5 | LPC 20:2-SN1 | LPC(20:2)>LPC(20:2/0:0)_and_LPC(0:0/20:2) | 1.7 |
| 79 | 898.7857 | 898.7855 | 0.2 | 327.4 | 322 | TG 18:1_18:2_18:2 | TG(54:5)>TG(18:1_18:2_18:2)_and_TG(16:0_16:0_22:5)_and_TG(16:0_18:1_20:4) M + NH3 | 1.7 |
| 80 | 757.6218 | 757.6216 | 0.3 | 290.5 | 295.6 | SM d38:2 | SM(d38:2) | -1.7 |
| 81 | 772.6216 | 772.6210 | 0.8 | 299.5 | 294.3 | PC O-36:2 | PC(O-36:2) | 1.8 |
| 82 | 673.5281 | 673.5277 | 0.7 | 272.0 | 276.9 | SM d32:2 | SM(d32:2) | -1.8 |
| 83 | 796.5847 | 796.5848 | 0.1 | 299.1 | 293.8 | PC 37:4 | PC(37:4) | 1.8 |
| 84 | 494.3242 | 494.3243 | 0.2 | 223.0 | 227.1 | LPC 16:1-SN1 | LPC(16:1)>LPC(16:1/0:0)_and_LPC(0:0/16:1) | -1.8 |
| 85 | 552.4025 | 552.4025 | 0.0 | 241.8 | 246.4 | LPC 20:0-SN1 | LPC(20:0)>LPC(20:0/0:0)_and_LPC(0:0/20:0) | -1.9 |
| 86 | 886.7856 | 886.7852 | 0.4 | 327.2 | 321.1 | TG 17:1_18:1_18:2 | TG(53:4)>TG(17:0_18:2_18:2)_and_TG(17:1_18:1_18:2) M + NH3 | 1.9 |
| 87 | 518.3242 | 518.3242 | 0.1 | 230.4 | 226.1 | LPC 18:3-SN1 | LPC(18:3)>LPC(18:3/0:0)_and_LPC(0:0/18:3) | 1.9 |
| 88 | 810.6008 | 810.6011 | 0.4 | 302.4 | 296.7 | PC 18:0_20:4 | PC(38:4) | 1.9 |

| | | | | | | | | |
|---|---|---|---|---|---|---|---|---|
| 89 | 677.5598 | 677.5582 | 2.3 | 279.4 | 285 | SM d32:0 | SM(d32:0) | -2.0 |
| 90 | 904.8324 | 904.8316 | 0.9 | 334.1 | 327.7 | TG 18:0_18:1_18:1 | TG(54:2)>TG(16:0_18:1_20:1)_and_TG(18:0_18:1_18:1) M + NH3 | 2.0 |
| 91 | 773.6532 | 773.6531 | 0.1 | 294.9 | 300.8 | SM d39:1 | SM(d39:1) | -2.0 |
| 92 | 896.7701 | 896.7699 | 0.2 | 326.8 | 320.4 | TG 18:2_18:2_18:2 | TG(54:6)>TG(16:0_18:2_20:4) M + NH3 | 2.0 |
| 93 | 890.8169 | 890.8168 | 0.1 | 331.7 | 324.9 | TG 17:0_18:1_18:1 | TG(53:2) M + NH3 | 2.1 |
| 94 | 759.6376 | 759.6373 | 0.4 | 291.9 | 298.2 | SM d38:1 | SM(d38:1) | -2.1 |
| 95 | 544.3399 | 544.3399 | 0.1 | 237.5 | 232.6 | LPC 20:4-SN1 | LPC(20:4)>LPC(20:4/0:0)_and_LPC(0:0/20:4) | 2.1 |
| 96 | 816.6475 | 816.6468 | 0.9 | 307.2 | 300.8 | PC 38:1 | PC(38:1) | 2.1 |
| 97 | 508.3400 | 508.3400 | 0.1 | 227.5 | 232.5 | LPC 17:1-SN2 | LPC(17:1)>LPC(17:1/0:0)_and_LPC(0:0/17:1) | -2.1 |
| 98 | 701.5595 | 701.5615 | 2.8 | 277.2 | 283.5 | SM d18:1_16:1 | SM(d34:2) | -2.2 |
| 99 | 796.6217 | 796.6210 | 0.9 | 304.1 | 297.5 | PC O-38:4 | PC(O-38:4) | 2.2 |
| 100 | 960.8951 | 960.8946 | 0.5 | 346.0 | 338.4 | TG 18:1_18:1_22:0 | TG(58:2) M + NH3 | 2.2 |
| 101 | 494.3242 | 494.3244 | 0.3 | 223.3 | 228.5 | LPC 16:1-SN2 | LPC(16:1)>LPC(16:1/0:0)_and_LPC(0:0/16:1) | -2.3 |
| 102 | 782.5693 | 782.5701 | 0.9 | 297.1 | 290.4 | PC 16:0_20:4 | PC(36:4) | 2.3 |
| 103 | 884.7701 | 884.7688 | 1.4 | 327.0 | 319.5 | TG 17:1_18:2_18:2 | TG(53:5)>TG(17:1_18:2_18:2) M + NH3 | 2.3 |
| 104 | 894.7543 | 894.7539 | 0.4 | 326.2 | 318.5 | TG 18:2_18:2_18:3 | TG(54:7)>TG(16:0_16:1_22:6)_and_TG(16:0_18:2_20:5) M + NH3 | 2.4 |
| 105 | 800.6160 | 800.6174 | 1.8 | 302.7 | 295.5 | PC 37:2 | PC(37:2) | 2.4 |
| 106 | 717.5906 | 717.5905 | 0.1 | 282.5 | 289.8 | SM d35:1 | SM(d35:1) | -2.5 |
| 107 | 812.6165 | 812.6163 | 0.2 | 304.8 | 297 | PC 38:3 | PC(38:3) | 2.6 |
| 108 | 958.8795 | 958.8787 | 0.8 | 345.4 | 336.4 | TG 18:1_18:2_22:0 | TG(58:3) M + NH3 | 2.7 |
| 109 | 780.5903 | 780.5903 | 0.0 | 298.5 | 290.7 | PC O-37:5 | PC(O-37:5) | 2.7 |
| 110 | 522.3558 | 522.3562 | 0.7 | 228.6 | 235.2 | LPC 18:1-SN1 | LPC(18:1)>LPC(18:1/0:0)_and_LPC(0:0/18:1) | -2.8 |
| 111 | 647.5126 | 647.5122 | 0.6 | 267.2 | 275 | SM d18:1_12:0 | SM(d30:1) | -2.8 |
| 112 | 946.8795 | 946.8787 | 0.9 | 345.3 | 335.8 | TG 18:1_18:1_21:0 | TG(57:2) M + NH3 | 2.8 |

| | | | | | | | | |
|---|---|---|---|---|---|---|---|---|
| 113 | 572.3711 | 572.3710 | 0.2 | 245.5 | 238.6 | LPC 22:4-SN1 | LPC(22:4)>LPC(22:4/0:0)_and_LPC(0:0/22:4) | 2.9 |
| 114 | 744.5543 | 744.5541 | 0.2 | 288.8 | 280.6 | | PC(33:2) | 2.9 |
| 115 | 689.5595 | 689.5599 | 0.6 | 275.5 | 283.8 | SM d33:1 | SM(d33:1) | -2.9 |
| 116 | 731.6065 | 731.6062 | 0.4 | 284.0 | 292.6 | SM d18:1_18:0 | SM(d36:1) | -2.9 |
| 117 | 814.6321 | 814.6311 | 1.2 | 307.2 | 298.4 | PC 38:2 | PC(38:2) | 2.9 |
| 118 | 802.6322 | 802.6332 | 1.3 | 307.1 | 298.3 | PC 37:1 | PC(37:1) | 3.0 |
| 119 | 675.5438 | 675.5447 | 1.4 | 272.1 | 280.5 | SM d32:1 | SM(d32:1) | -3.0 |
| 120 | 705.5905 | 705.5891 | 1.9 | 281.2 | 290.1 | SM d34:0 | SM(d34:0) | -3.1 |
| 121 | 714.6188 | 714.6185 | 0.5 | 286.8 | 295.9 | CE 22:6 | CE(22:6)M+NH3 | -3.1 |
| 122 | 482.3242 | 482.3243 | 0.3 | 220.9 | 228.3 | LPC 15:0-SN1 | LPC(15:0)>LPC(15:0/0:0)_and_LPC(0:0/15:0) | -3.2 |
| 123 | 782.5694 | 782.5695 | 0.2 | 296.8 | 287.5 | PC 18:2_18:2 Cis | PC(36:4) | 3.2 |
| 124 | 522.3554 | 522.3554 | 0.1 | 229.2 | 236.9 | LPC 18:1-SN2 | LPC(18:1)>LPC(18:1/0:0)_and_LPC(0:0/18:1) | -3.3 |
| 125 | 808.5823 | 808.5848 | 3.2 | 303.9 | 294.2 | PC 38:5 | PC(36:2) M + Na | 3.3 |
| 126 | 703.5749 | 703.5778 | 4.2 | 276.8 | 286.3 | SM d18:1_16:0 | SM(d34:1) | -3.3 |
| 127 | 780.5539 | 780.5542 | 0.3 | 297.0 | 287.3 | PC 36:5 | PC(36:5) | 3.4 |
| 128 | 756.5541 | 756.5540 | 0.1 | 292.8 | 283.3 | PC 34:3 | PC(34:3) | 3.4 |
| 129 | 568.3398 | 568.3397 | 0.2 | 244.7 | 236.4 | LPC 22:6-SN2 | LPC(22:6)>LPC(22:6/0:0)_and_LPC(0:0/22:6) | 3.5 |
| 130 | 570.3554 | 570.3553 | 0.2 | 244.7 | 236.4 | LPC 22:5-SN1 | LPC(22:5)>LPC(22:5/0:0)_and_LPC(0:0/22:5) | 3.5 |
| 131 | 824.6166 | 824.6160 | 0.8 | 310.3 | 299.7 | PC 39:4 | PC(39:4) | 3.5 |
| 132 | 768.5538 | 768.5540 | 0.3 | 296.5 | 286.2 | PC 35:4 | PC(35:4) | 3.6 |
| 133 | 820.5850 | 820.5850 | 0.0 | 307.5 | 296.6 | PC 39:6 | PC(39:6) | 3.7 |
| 134 | 924.8013 | 924.8006 | 0.7 | 339.8 | 327.6 | TG 18:1_18:1_20:4 | TG(56:6)>TG(18:1_18:1_20:4)_and_TG(16:0_18:1_22:5)_and_TG(18:1_18:2_20:3) M + NH3 | 3.7 |
| 135 | 770.5695 | 770.5696 | 0.2 | 299.0 | 288 | PC 35:3 | PC(35:3) | 3.8 |

| | | | | | | | | |
|---|---|---|---|---|---|---|---|---|
| 136 | 806.5694 | 806.5692 | 0.3 | 304.4 | 293.1 | PC 38:6 (PC 16:0_22:6) | PC(38:6) | 3.8 |
| 137 | 754.5384 | 754.5381 | 0.4 | 292.8 | 281.9 | PC 34:4 | PC(34:4) | 3.9 |
| 138 | 948.8952 | 948.8944 | 0.9 | 350.4 | 337.2 | TG 16:0_16:1_25:0 | TG(57:1)>TG(16:0_18:1_23:0)_and_TG(18:0_18:1_21:0) M + NH3 | 3.9 |
| 139 | 834.6007 | 834.6002 | 0.6 | 311.6 | 299.5 | PC 18:0_22:6 | PC(40:6) | 4.0 |
| 140 | 932.8640 | 932.8642 | 0.2 | 346.7 | 333.3 | TG 18:1_18:1_20:0 | TG(56:2) M + NH3 | 4.0 |
| 141 | 568.3399 | 568.3396 | 0.6 | 244.9 | 235.2 | LPC 22:6-SN1 | LPC(22:6)>LPC(22:6/0:0)_and_LPC(0:0/22:6) | 4.1 |
| 142 | 774.6008 | 774.6011 | 0.3 | 304.4 | 292.4 | PC 35:1 | PC(35:1) | 4.1 |
| 143 | 690.6185 | 690.6187 | 0.2 | 281.8 | 294 | CE 20:4 | CE(20:4)M+NH3 | -4.1 |
| 144 | 510.3555 | 510.3555 | 0.1 | 226.4 | 236.4 | LPC 17:0-SN1 | LPC(17:0)>LPC(17:0/0:0)_and_LPC(0:0/17:0) | -4.2 |
| 145 | 468.3085 | 468.3084 | 0.2 | 215.1 | 224.6 | LPC 14:0-SN1 | LPC(14:0)>LPC(14:0/0:0)_and_LPC(0:0/14:0) | -4.2 |
| 146 | 718.5383 | 718.5383 | 0.1 | 292.1 | 279.7 | PC 17:0_14:1 | PC(31:1) | 4.4 |
| 147 | 806.5695 | 806.5689 | 0.7 | 304.3 | 291.4 | PC 38:6 (PC 18:2_20:4) | PC(38:6) | 4.4 |
| 148 | 496.3399 | 496.3405 | 1.1 | 221.7 | 232.1 | LPC 16:0-SN1 | LPC(16:0)>LPC(16:0/0:0)_and_LPC(0:0/16:0) | -4.5 |
| 149 | 468.3085 | 468.3087 | 0.4 | 215.4 | 225.5 | LPC 14:0-SN2 | LPC(14:0)>LPC(14:0/0:0)_and_LPC(0:0/14:0) | -4.5 |
| 150 | 524.3710 | 524.3717 | 1.3 | 229.2 | 240.1 | LPC 18:0-SN1 | LPC(18:0)>LPC(18:0/0:0)_and_LPC(0:0/18:0) | -4.5 |
| 151 | 930.8480 | 930.8477 | 0.2 | 346.3 | 330.9 | TG 18:1_18:1_20:1 | TG(56:3)>TG(18:1_18:1_20:1) M + NH3 | 4.7 |
| 152 | 818.6057 | 818.6057 | 0.0 | 312.1 | 298 | PC O-40:7 | PC(O-40:7) | 4.7 |
| 153 | 848.6525 | 848.6528 | 0.3 | 318.7 | 304.2 | PC O-42:6 | PC(O-42:6) | 4.8 |
| 154 | 866.6639 | 866.6643 | 0.4 | 323.5 | 308.8 | PC 42:4 | PC(42:4) | 4.8 |
| 155 | 524.3713 | 524.3715 | 0.3 | 229.3 | 241 | LPC 18:0-SN2 | LPC(18:0)>LPC(18:0/0:0)_and_LPC(0:0/18:0) | -4.8 |
| 156 | 946.7856 | 946.7851 | 0.6 | 344.8 | 328.5 | TG 18:1_20:4_20:4 | TG(56:6)>TG(18:1_18:1_20:4)_and_TG(16:0_18:1_22:5)_and_TG(18:1_18:2_20:3) M + NH3 | 5.0 |
| 157 | 820.6210 | 820.6209 | 0.1 | 314.2 | 298.9 | PC O-40:6 | PC(O-40:6) | 5.1 |
| 158 | 944.7701 | 944.7694 | 0.7 | 343.7 | 326.7 | TG 16:0_20:4_22:6 | TG(58:10) M + NH3 | 5.2 |

| | | | | | | | | |
|---|---|---|---|---|---|---|---|---|
| 159 | 832.5845 | 832.5846 | 0.1 | 312.5 | 296.8 | PC 40:7 | PC(40:7) | 5.3 |
| 160 | 496.3400 | 496.3402 | 0.4 | 220.9 | 233.7 | LPC 16:0-SN2 | LPC(16:0)>LPC(16:0/0:0)_and_LPC(0:0/16:0) | -5.5 |
| 161 | 864.6475 | 864.6473 | 0.2 | 323.6 | 305.5 | PC 42:5 | PC(42:5) | 5.9 |
| 162 | 824.6524 | 824.6529 | 0.7 | 321.0 | 302.8 | PC O-40:4 | PC(O-40:4) | 6.0 |
| 163 | 830.5695 | 830.5690 | 0.6 | 312.2 | 294 | PC 20:4_20:4 Cis | PC(40:8) | 6.2 |
| 164 | 864.8030 | 864.8019 | 1.2 | 340.9 | 320.8 | TG 16:0_17:0_18:1 | TG(51:1) M + NH3 | 6.3 |
| 165 | 856.5847 | 856.5846 | 0.1 | 320.2 | 297.6 | PC 42:9 | PC(42:9) | 7.6 |
| 166 | 766.5748 | 766.5747 | 0.0 | 320.3 | 289.5 | PC O-36:5 (PC O-16:0_20:5) | PC(O-36:5) | 10.6 |

**Supplementary Table S14. Summary of lipid features with matched $^{Orbi}$CCS to reference $^{DT}$CCS dataset for experiments conducted at a resolution of 90,000 and turbopump speed 68%.**

| # | m/z_this study | m/z_database | mzError_ppm | $^{Orbi}$CCS_this study | $^{DT}$CCS_database | Name_database | Name_This study | % error |
|---|---|---|---|---|---|---|---|---|
| 1 | 729.5906 | 729.5910 | 0.5 | 285.1 | 285.3 | SM 36:2 | SM(d36:2) | -0.1 |
| 2 | 538.3870 | 538.3872 | 0.3 | 240.1 | 240.5 | LPC 19:0 | LPC(19:0)>LPC(19:0/0:0)_and_LPC(0:0/19:0) | -0.2 |
| 3 | 801.6847 | 801.6849 | 0.3 | 303.9 | 302.3 | SM 41:1 | SM(d41:1) | 0.5 |
| 4 | 759.6376 | 759.6380 | 0.6 | 291.9 | 293.4 | SM 38:1 | SM(d38:1) | -0.5 |
| 5 | 745.6220 | 745.6223 | 0.4 | 291.7 | 289.8 | SM 37:1 | SM(d37:1) | 0.7 |
| 6 | 773.6532 | 773.6536 | 0.5 | 294.9 | 297 | SM 39:1 | SM(d39:1) | -0.7 |
| 7 | 786.6007 | 786.6012 | 0.7 | 295.8 | 293.3 | PC (18:1/18:1) (del9-trans) | PC(36:2) | 0.8 |
| 8 | 787.6688 | 787.6693 | 0.6 | 302.7 | 299.1 | SM 40:1 | SM(d40:1) | 1.2 |
| 9 | 799.6690 | 799.6693 | 0.4 | 303.9 | 300.1 | SM 41:2 | SM(d41:2) | 1.3 |
| 10 | 785.6534 | 785.6536 | 0.2 | 301.0 | 296.8 | SM 40:2 | SM(d40:2) | 1.4 |
| 11 | 731.6065 | 731.6067 | 0.3 | 284.0 | 288.4 | SM 36:1 | SM(d36:1) | -1.5 |
| 12 | 703.5749 | 703.5754 | 0.7 | 276.8 | 281.2 | SM 34:1 | SM(d34:1) | -1.6 |
| 13 | 522.3554 | 522.3559 | 0.9 | 229.2 | 233.2 | LPC 18:1 | LPC(18:1)>LPC(18:1/0:0)_and_LPC(0:0/18:1) | -1.7 |
| 14 | 760.5850 | 760.5856 | 0.8 | 293.1 | 287.9 | PC (18:1(9Z)/16:0) | PC(34:1) | 1.8 |
| 15 | 827.7001 | 827.7006 | 0.6 | 311.3 | 305.7 | SM 43:2 | SM(d43:2) | 1.8 |
| 16 | 730.5384 | 730.5387 | 0.4 | 284.0 | 278.4 | PC (16:1/16:1) (del9-cis) | PC(32:2) | 2.0 |
| 17 | 552.4025 | 552.4029 | 0.7 | 241.8 | 247 | 20:0 Lyso PC | LPC(20:0)>LPC(20:0/0:0)_and_LPC(0:0/20:0) | -2.1 |
| 18 | 813.6845 | 813.6849 | 0.5 | 308.6 | 302.2 | SM 42:2 | SM(d42:2) | 2.1 |
| 19 | 811.6686 | 811.6693 | 0.8 | 307.4 | 300.8 | SM 42:3 | SM(d42:3) | 2.2 |
| 20 | 808.5823 | 808.5832 | 1.1 | 303.9 | 296.9 | PC (18:1/18:1) (del9-trans) | PC(36:2) M + Na | 2.4 |
| 21 | 732.5540 | 732.5543 | 0.4 | 284.4 | 277.6 | PC 32:1 | PC(32:1) | 2.4 |
| 22 | 468.3085 | 468.3090 | 1.1 | 215.1 | 220.8 | LPC 14:0 | LPC(14:0)>LPC(14:0/0:0)_and_LPC(0:0/14:0) | -2.6 |
| 23 | 786.6007 | 786.6012 | 0.7 | 295.0 | 287.1 | PC 36:2 | PC(36:2) | 2.8 |
| 24 | 788.6165 | 788.6169 | 0.5 | 298.6 | 289.4 | PC 36:1 | PC(36:1) | 3.2 |
| 25 | 758.5693 | 758.5699 | 0.7 | 290.2 | 280.6 | PC 34:2 | PC(34:2) | 3.4 |
| 26 | 744.5543 | 744.5543 | 0.0 | 288.8 | 279.1 | PE (18:1/18:1) (del9-cis) | PC(33:2) | 3.5 |
| 27 | 524.3713 | 524.3716 | 0.6 | 229.3 | 238.8 | 18:0 Lyso PC | LPC(18:0)>LPC(18:0/0:0)_and_LPC(0:0/18:0) | -4.0 |
| 28 | 496.3400 | 496.3403 | 0.6 | 220.9 | 231.4 | 16:0 Lyso PC | LPC(16:0)>LPC(16:0/0:0)_and_LPC(0:0/16:0) | -4.5 |

| | | | | | | | | |
|---|---|---|---|---|---|---|---|---|
| 29 | 524.3710 | 524.3716 | 1.2 | 229.2 | 240.7 | 2-18:0 Lyso PC | LPC(18:0)>LPC(18:0/0:0)_and_LPC(0:0/18:0) | -4.8 |
| 30 | 756.5541 | 756.5543 | 0.3 | 292.8 | 278.2 | PC 34:3 | PC(34:3) | 5.3 |
| 31 | 728.5593 | 728.5594 | 0.1 | 288.4 | 273.5 | PE (O-36:3) | PE(O-36:3)>PE(O-18:1/18:2) | 5.5 |
| 32 | 774.6008 | 774.6013 | 0.6 | 304.4 | 285.8 | PC 35:1 | PC(35:1) | 6.5 |
| 33 | 400.3421 | 400.3427 | 1.4 | 190.7 | 214.7 | Carnitine 16:0 | Car(16:0) | -11.2 |
| 34 | 344.2795 | 344.2782 | 3.8 | 172.5 | 199.5 | Carnitine 12:0 | Car(12:0) | -13.5 |
